# Bayesian prediction of multivariate ecology from phenotypic data yields new insights into the diets of extant and extinct taxa

**DOI:** 10.1101/2022.05.05.490807

**Authors:** Anna L. Wisniewski, Jonathan A. Nations, Graham J. Slater

## Abstract

Morphology often reflects ecology, enabling the prediction of ecological roles for taxa that lack direct observations such as fossils. In comparative analyses, ecological traits, like diet, are often treated as categorical, which may aid prediction and simplify analyses but ignores the multivariate nature of ecological niches. Futhermore, methods for quantifying and predicting multivariate ecology remain rare. Here, we ranked the relative importance of 13 food items for a sample of 88 extant carnivoran mammals, and then used Bayesian multilevel modeling to assess whether those rankings could be predicted from dental morphology and body size. Traditional diet categories fail to capture the true multivariate nature of carnivoran diets, but Bayesian regression models derived from living taxa have good predictive accuracy for importance ranks. Using our models to predict the importance of individual food items, the multivariate dietary niche, and the nearest extant analogs for a set of data-deficient extant and extinct carnivoran species confirms long-standing ideas for some taxa, but yields new insights about the fundamental dietary niches of others. Our approach provides a promising alternative to traditional dietary classifications. Importantly, this approach need not be limited to diet, but serves as a general framework for predicting multivariate ecology from phenotypic traits.

## Introduction

Interspecific interactions, the structure of communities, and the persistence of lineages through time are all mediated by the ecological niches that each species occupies and the degree of overlap between them (Hayward and Slotow, 2009; Hutchinson, 1957, 1959; Peralta et al., 2020; Pigot et al., 2020; Vannette and Fukami, 2014). Diet is a particularly important axis of niche differentiation; communities of closely related species are often structured along dietary axes (Hertel, 1994; Kiltie, 1988; Pigot et al., 2020) and dietary ecology fosters phenotypic variance as organisms evolve physiological, morphological, and behavioral adaptations to efficiently acquire and process different food resources (Arnold, 1983; Carroll et al., 2004; Holzman et al., 2012; Santana and Dumont, 2009; Vincent et al., 2009). On short time scales, competition for dietary resources can lead to character displacement and divergence in the morphology of ecologically and phylogenetically similar taxa (Grant and Grant, 2006; Pfennig and Pfennig, 2010; Van Valkenburgh and Wayne, 1994), while on macroevolutionary time-scales dietary ecology may promote differential patterns of speciation and extinction as dynamics between competing lineages and resource availability change over time (Burin et al., 2016; Farrell, 1998; Lobato et al., 2014; Poore et al., 2017; Price et al., 2012; Wiens et al., 2015).

Given the importance of diet in eco-evolutionary patterns and processes, the relationship between diet and phenotype is now manifest in many animal systems. For example, the morphology of insect mouthparts (Krenn, 2019), snake fangs (Cleuren et al., 2021), bird beaks (Natale and Slater, 2022; Olsen, 2017; Pigot et al., 2020), mammalian crania and jaws (Janis and Erhardt, 1988; Morales-Garćıa et al., 2021; Spencer, 1995), and vertebrate (especially mammalian) teeth (Christensen and Melstrom, 2021; Melstrom, 2017; Pollock et al., 2022) are all predictive of diet. The performance-mediated relationship between morphology and ecology is particularly important in analyses that include taxa for which ecological data are not directly available such as fossils, which can be critical for robust inference of macroevolutionary parameters (Finarelli and Flynn, 2006; Slater et al., 2012). However, effectively integrating taxa of unknown ecological function into evolutionary studies requires not only that we can construct predictive models to estimate ecological traits from their morphological characters, but also that we can effectively and accurately quantify the range of ecological roles that species may play in the first place.

Problems quickly arise when determining precisely how to categorize diet based on the range of food types that are used and the frequency with which any particular type is consumed. Diet is typically reduced to one of a small number of discrete categories, often based on trophic level (e.g., ‘herbivore’, ‘carnivore’ and ‘omnivore’; Christensen and Melstrom, 2021; Price and Hopkins, 2015; Price et al., 2012). Other studies have expanded this range to include more refined categories, such as ‘browser’, ‘grazer’, ‘mixed feeder’ (Toljagić et al., 2018) and ‘hyper-’, ‘meso-’, ‘hypocarnivore’ (Slater, 2015; Van Valkenburgh, 1988), or even more finely subdivided categories based on variation in foraging behavior and micro-habitat use (Pigot et al., 2020, 2016; Verde Arregoitia and D’Eĺıa, 2021). However, all of these schemes still require that taxa be pigeonholed into a single discrete group. This categorization of diet may be unsatisfactorily simplistic for several reasons. Finely divided categories are often created for a particular taxonomic subset, precluding the application of the same coding scheme across higher-level macroevolutionary studies. For example, a ‘hypercarnivorous’ weasel, rat, and mole may depredate very different animals (e.g. mice, beetles, and worms), leaving few options but to create a large number of categories (e.g. ‘vertebrate hypercarnivore’, ‘soft invertebrate hypercarnivore’, etc). Furthermore, species frequently consume different proportions of foods, often opportunistically, or have diets that cross category boundaries. For example, many squirrel species, such as the Antelope Ground Squirrel, will occasionally eat vertebrates despite seeds making up an overwhelming portion of their diet (Bradley, 1968). Categorizing a squirrel as ‘granivore’ necessarily ignores the less common, but still nutritionally important, vertebrate food items that are consumed, while ‘omnivore’ may be too broad a dietary classification in this case. Pineda-Munoz and Alroy (2014) suggested that diet be classified based on both the primary and secondary food types consumed, with the term ‘generalist’ reserved for taxa in which no individual food type comprises the majority of the diet. Although this scheme is an improvement on previous approaches, it still fails to capture the full breadth of diet variation by ignoring the use of foods of tertiary or lower importance. Grundler and Rabosky (2021) overcame this issue by treating diet as multivariate rather than categorical, and inferred the presence and location of dietary niche shifts over snake phylogeny by using a database of proportional occurrences of food items in the diet of 882 species (Grundler, 2020). However, although some level of unobserved data can be accommodated by this approach, it cannot readily be extended to datasets with large numbers of data-deficient taxa such as those based on fossils.

Ideally, animal diet would be codified in a way that preserves its inherently multidimensional structure, while also being amenable to prediction from the morphology of poorly studied or extinct taxa, and, ultimately, to large-scale macroevolutionary and macroecological analysis. Rojas et al. (2011) and Kissling et al. (2014) apparently independently developed a coding scheme that, rather than assigning taxa to a single discrete diet category, ranks the relative importance of different food types in a species’ diet based on the use of keywords in synoptic reviews and primary ecological studies. The result is that diet is represented not as a single code in a classification scheme but, rather, as a vector of ordinal variables. Rojas et al. (2011) and Kissling et al. (2014) ultimately used their codings to classify taxa to standard specialist categories, while Rojas et al. (2018) used them to develop a univariate continuous variable spanning herbivory to carnivory (see also López-Aguirre et al., 2022). However, the original importance codings present a holistic, multivariate description of diet that may yield previously overlooked insights into the form-function relationship between phenotype and ecology and the structure of dietary niches. For example, a species may experience selective pressure for traits that are associated with efficient acquisition and processing of seasonally or infrequently consumed food items (‘fall-back foods’ Marshall et al., 2009), as these nutritional sources may increase an individual’s fitness when preferred food items are unavailable. Although item importance ranking has several advantages over traditional classification schemes, it is yet to be widely adopted in ecomorphological studies (but see Machado, 2020).

Here, we investigate whether ordinal ranking of food type importance can be predicted from ecomorphological traits using a well-established system with rigorously defined functional variation: the molar dentition of terrestrial (i.e. non-pinniped) members of the mammalian order Carnivora. We seek to understand how different functional aspects of the carnivoran dentition, such as sharpness, complexity, and surface area, correlate with the relative importance of different food items. Carnivora is an ideal group for such a study. The order is taxonomically and ecologically diverse, comprising over 300 living species (Burgin et al., 2018) that occupy a wide range of dietary niches. Extant carnivorans have been well studied, with a wealth of information available regarding their phylogenetic relationships (Eizirik et al., 2010; Nyakatura and Bininda-Emonds, 2012; Slater and Friscia, 2019), dietary diversity (Ewer, 1998), and ecomorphological variation (Van Valkenburgh and Koepfli, 1993; Friscia et al., 2007; Sacco and Van Valkenburgh, 2004; Van Valkenburgh, 1988). Additionally, carnivorans possess a rich fossil record, providing ample opportunity to predict the ecologies of extinct species. We conducted an extensive literature review to evaluate the relative importance of 13 different food items for 88 species of extant carnivoran using the approach of Rojas et al. (2011) and Kissling et al. (2014). We then fit a series of Bayesian multilevel regression models with linear and topographic dental metrics as predictors and the relative importance of diet items as ordinal responses. Finally, we used the best-fitting models for each food item to predict the importance of the 13 food types for a sample of data-deficient extant carnivorans, as well as several species of extinct carnivorans spanning from the Eocene (*∼* 50 myr) to the latest Pleistocene (*∼* 13 kyr).

## Materials and Methods

### Data Collection

#### Morphological data collection

We compiled data for four morphological metrics that collectively describe the grinding area, relief, complexity, and sharpness of the lower molar tooth row from 99 species of extant non-pinniped carnivorans. Species-mean values for Relative Lower Grinding Area (RLGA: Van Valkenburgh, 1988), a measure of the size of the lower molar tooth row dedicated to grinding as opposed to slicing, were obtained from Slater and Friscia (2019) and Friscia et al. (2007). We generated novel topographic metrics from 3D surface scans of carnivoran lower first (m1) and second (m2) molars (see Supplementary Methods for scanning details). Molar scans and measurements were obtained from specimens housed in the mammal collections of the Field Museum of Natural History, Chicago (see Table S1 for specimen numbers). Relief index (RFI: Boyer, 2008) is the ratio of the 3D surface area of a tooth to its 2D planar area and is a measure of topographic relief. Orientation Patch Count rotated (OPCr: Evans et al., 2007; Wilson et al., 2012) measures the complexity of the tooth by counting the number of contiguous patches on the tooth surface that share a common orientation. Because the orientation of each point is sensitive to the alignment of the tooth model to the global coordinate system, counts are calculated over several small rotations of the model and averaged (Wilson et al., 2012). RFI and OPCr were calculated using the molar package (Pampush et al., 2016) for R (R Core Team, 2022). Finally, the Dirichlet Normal Energy (DNE) measures the average curvature of a surface by calculating its “bending energy” (Bunn et al., 2011) and captures overall tooth sharpness. We used “a robustly implemented algorithm for DNE” (ariaDNE: Shan et al., 2019) that is less sensitive to artifacts due to 3D modeling, such as smoothing, than early DNE algorithms, and which is implemented via MATLAB scripts provided by Shan et al. (2019). For carnivoran species without an m2, we assigned a value of zero for all dental topographic measurements. All surface scans are available on the MorphoSource digital repository (project ID: 000501405), and all dental measurement values are available in Table S1

We visualized patterns of covariation in the topographic data by performing a principal components analysis on the covariance matrix of standardized dental data using the prcomp function in the stats package in R (R Core Team, 2022). To evaluate phylogenetic signal in carnivoran dental shape we calculated *K_mult_*(Adams, 2014; Blomberg et al., 2003) using the phylosignal function in the geomorph library (Adams et al., 2021; Baken et al., 2021).

#### Dietary Data Collection

We sourced dietary information for each carnivoran species in our dataset through a review of species accounts and primary ecological studies. We followed Machado’s (2020) coding scheme of canid diets, modified from Kissling et al. (2014), to rank the relative importance of 13 food types: large mammal, small mammal, bird, herptile (reptiles and amphibians), fish, egg, carrion, hard-bodied invertebrate, soft-bodied invertebrate, seed (including nuts), root, fruit, and plant (including leaves and stems). Canids span almost the entire breadth of dietary diversity in carnivorans, and this system is therefore appropriate for the present study. However, food items can easily be modified for subsequent studies, depending on taxonomic and ecological sampling. For example, few extant carnivorans place high importance on leaves or stems, allowing for use of a single broad “plant” type, but this could be split into “grasses”, “herbaceous plants”, “leaves”, and “woody stems” if artiodactyls and perissodactyls were added in subsequent work. We defined large mammals as species with a mean mass greater than five kilograms. Hard and soft-bodied invertebrates were designated as separate food types because sclerotinized and unsclerotinized cuticles have different material properties that require different mechanical solutions to fracture (Freeman, 1979; Strait and Vincent, 1998). We coded larvae as soft-bodied, irrespective of the properties of the adult cuticle.

Following Kissling et al. (2014) and Machado (2020), but with some minor modifications, food types were assigned to ranks based on the use of keywords and phrases in synoptic reviews or primary analyses of diet in the focal species (Table 1). In contrast with Kissling et al. (2014), we avoided broad or superficial sources (e.g., Nowak and Walker, 1999) when coding dietary ranks and attempted to validate primary sources where review papers, such as *Mammalian Species* accounts from The American Society of Mammalogists, were used. A single instance of a species consuming a food type was sufficient to assign it rank 2 (Low Importance), regardless of inferred nutritional importance. Seasonally and regionally important foods were generally considered important in the context of the species’ diet (Porter et al., 2022). We note that this approach may result in higher rankings for some food items, due to individual or population-level specializations (e.g., Bolnick et al., 2003; DeSantis et al., 2022), than are representative of the overall species mean. For this reason, our dietary importance rankings should be conservatively interpreted as describing the fundamental dietary niche of each taxon, rather than the realized niche of any one population (Hutchinson, 1957). In cases where insufficient data were available to assign ranks to food items, such as when a source only listed a set of foods that are eaten or that have been recovered from stomach contents without using keywords to describe the frequency or importance of each, we coded the taxon as unknown. All food importance scores are available in Table S1.

**Table 1:**
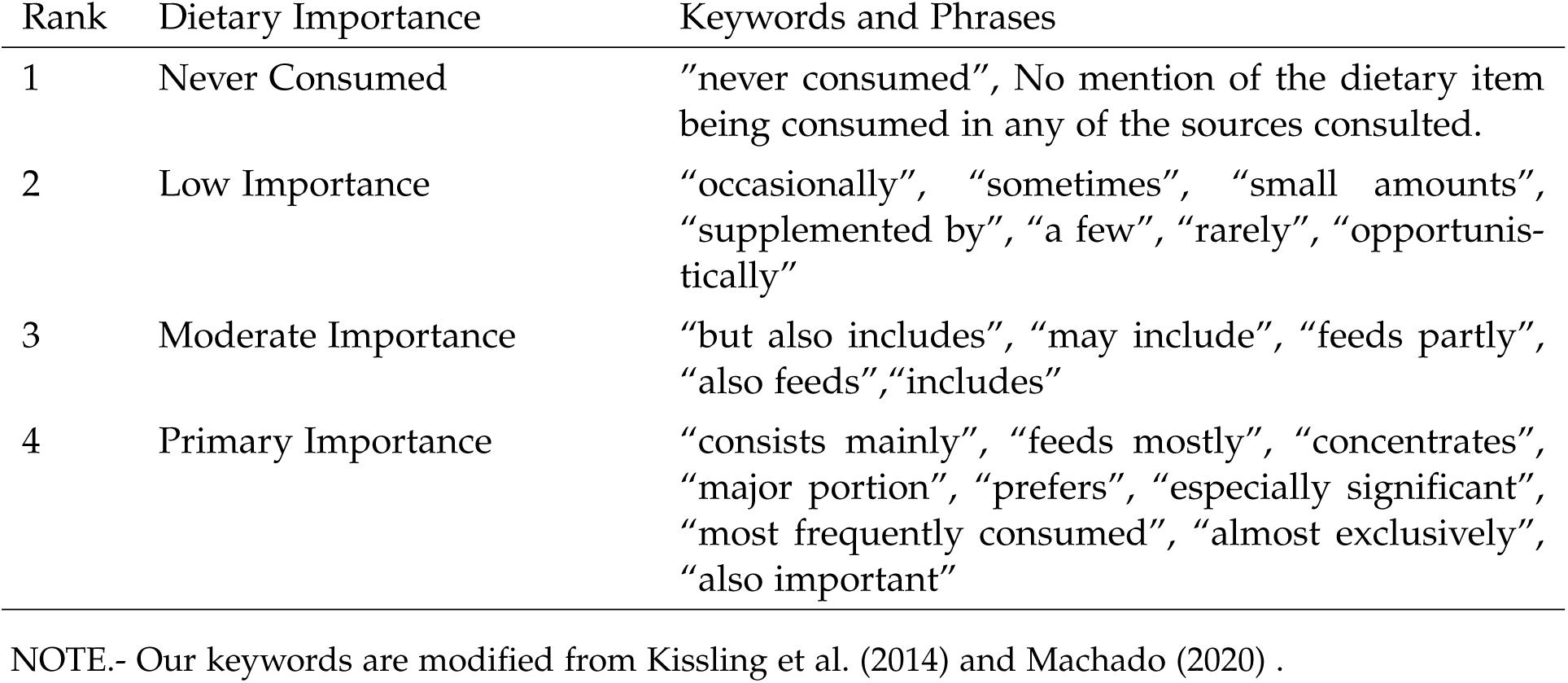
Dietary Importance Ranking Scheme

To visualize general patterns of covariation in dietary item importance, we performed a principal components analysis on the polychoric correlation matrix of importance scores, computed using the polychor function in the polycor library (Fox, 2021). Polychoric correlations differ from Pearson’s correlations in that they do not assume that the input variables themselves are continuously distributed but that they are discrete outcomes of a liability threshold process on a normally distributed latent variable. As such, they are appropriate for ordinating discrete ordered states, such as Likert scores (Holgado-Tello et al., 2010). We then used the mclust library (Scrucca et al., 2016) to determine the optimal number of clusters of taxa in the resulting principal component scores based on a finite Gaussian mixture model, with a maximum of 20 clusters permitted and model selection based on the Bayesian Information Criterion (BIC).

For comparison with previous categorical dietary groupings, we classified the species in our dataset using four often-used carnivoran dietary classification schemes: The PanTHERIA database dietary classification scheme (Jones et al., 2009), which ranks 2161 mammal species into three diet categories (omnivore, herbivore, carnivore); the classification scheme of Van Valkenburgh (1988), which places carnivorans in three dietary categories (hyper-, meso-, and hypocarnivore); the four-category scheme of Pineda-Munoz et al. (2017) that uses secondary dietary categories where an alternate food source comprises a substantial proportion of the diet after the primary food source (in this case, hypercarnivore, hypocarnivore-insectivore, hypocarnivoreherbivore, and mesocarnivore); and diet categories from Animal Diversity Web (https://an imaldiversity.org/, accessed on February 8, 2022), an increasingly popular source of dietary information for comparative analyses (e.g. Goswami et al., 2022; Lomolino et al., 2012; Morales-Garćıa et al., 2021), which places the taxa in our dataset into 6 categories (carnivore, frugivore, herbivore, insectivore, omnivore, and piscivore). We used the Van Valkenburgh (1988) and Pineda-Munoz et al. (2017) classifications reported in the supporting information of Hopkins et al. (2022). We then recomputed optimal clustering schemes in our importance data but with the number of clusters set equal to the number of groups in each of the four discrete dietary schemes, and compared the resulting classifications using the adjusted Rand Index as computed by the adjustedRandIndex function in the mclust library. The Rand Index compares classification schemes based on matches, with its adjusted version accounting for the number of matches that are expected to occur due to chance (Hubert and Arabie, 1985). An adjusted Rand Index of 0 is expected for two completely random classification schemes, while a value of 1 indicates perfect agreement between the two.

### Bayesian multilevel modeling

Dietary rankings are neither continuous predictors nor discrete categories but, instead, are categories with a natural order. Therefore, to test the hypothesis that food type importance can be predicted from dental topography in carnivorans we used ordinal regression, with diet ranking represented as a cumulative distribution (Bürkner and Vuorre, 2019). Like polychoric PCA, cumulative ordinal regression models assume the response variable is drawn from a continuous distribution split by K thresholds that separate K + 1 groups; for 4 dietary rankings, 1-4, the model will estimate 3 thresholds within the total distribution of rankings (Bürkner and Vuorre, 2019). The distances between thresholds in a cumulative distribution need not be equal. We used food item importance rankings as our response variables and the dental topographic metrics for m1 and m2 as our predictors. Body mass can mediate the type of food that a species is capable of attaining and processing with a given occlusal topography (e.g., Carbone et al., 1999, 2007; Radloff and Du Toit, 2004), and we, therefore, modeled the interaction between the natural log of body mass, taken from PanTHERIA (Jones et al., 2009), and the m1 and m2 predictors. Modeling the interaction between body mass and dental morphology allows us to condition our estimates on the effect of body size and simultaneously quantify variation in the effect of tooth topology on diet along the body-size continuum. Species traits may co-vary due to evolutionary history, so we used a phylogenetic correlation matrix generated from the time-scaled molecular phylogeny of Slater and Friscia (2019) as a group-level predictor (de Villemereuil and Nakagawa, 2014; de Villemereuil et al., 2012; Lynch, 1991). This phylogenetic multilevel modeling approach jointly estimates the phylogenetic covariance in the data and conditions the model estimates on this covariance (Fulwood et al., 2021; McElreath, 2020).

Our models took the general form

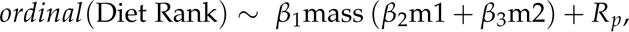

where *β*_1_ represents the effect of body mass on diet category, *β*_2_ and *β*_3_ represent the effect of m1 and m2 topography on diet category, and *R_p_* represents the phylogenetic correlation matrix. The interaction terms *β*_1_ × *β*_2_ and *β*_1_ × *β*_3_ represent the interactions between mass and m1 topography, and mass and m2 topography, respectively. For each dietary item, we fit five ordinal regression models: one for each of the three dental topographic measures, one for RLGA, and one model that included all 4 of the dental metrics. Models were fit using the R package brms (Bü rkner, 2017), an interface for the Bayesian probabilistic programming language Stan (Gelman et al., 2015). Working in a full Bayesian framework allows us to fit these complex ordinal models within a multilevel structure, to provide an intuitive measure of uncertainty in the results (i.e. probability), and to use regularizing priors to minimize overfitting risk. Data processing and post hoc analyses heavily relied on the tidyverse (Wickham et al., 2019), furrr (Vaughan and Dancho, 2022), and tidybayes (Kay, 2021) packages. All models and scripts are available in DataDryad (https://doi.org/10.5061/dryad.pc866t1rg).

#### Prior Predictive Checks

Before fitting the models, we standardized all linear predictors to z-scores as is recommended practice when working with predictors on different scales (Gelman et al., 2020; McElreath, 2020). For each tooth metric, we compared three different prior distributions for each response variable (ranks of food items): a normal distribution, a student-*T* distribution, and a custom Dirichlet distribution. We used prior predictive simulation to determine the best-calibrated parameters for each prior distribution that effectively capture the distribution of the dietary ranks while discouraging unrealistic values (Figure S1). We then used Bayesian leave-one-out (LOO) cross validation (Vehtari et al., 2017) to determine the prior distribution with the highest predictive power for the response variables. Bayesian LOO cross validation uses Pareto smoothed importance sampling to simulate the posterior of the model *n_sample_* times, leaving out a single data point (in our case, a single species) in each refitting. This approach identifies data points (i.e. species) that have an outsized influence on the posterior distributions, rewards accurate prediction while penalizing overfitting, and calculates a LOO score for each model that can be used for model comparison (Vehtari et al., 2017; Yao et al., 2018).

#### Model Fitting

After determining an appropriate prior distribution for each dietary item response, we ran each model for four chains, with 2000 iterations of warm-up and 2000 iterations of sampling. We used a *N* (0, 1) prior on each predictor variable (z-scores of tooth metrics and log(mass), and phylogenetic correlation matrix). Chain convergence was verified with the Gelman-Rubin 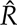 statistic (Gelman and Rubin, 1992). For each food item, we again used LOO cross validation to compare the predictive power of each of the 5 models, then estimated the model weights using stacking (Yao et al., 2018).

#### Model Validation & Predictive Accuracy

We used three approaches to validate our models. First, we used posterior predictive checking via the pp_check command in brms to confirm that the models indeed generated reasonable estimates of the distribution of importance ranks in original dietary data by plotting 500 draws from the posterior along with the empirical counts of the rankings for each food item (Figure S2). Second, we used our LOO model weights to model average over the 5 models for each food item, and then extracted the Pareto-*K* scores for each species for each food item. Pareto-*K* scores are diagnostics calculated by the Pareto smoothed importance sampling algorithm in LOO (Yao et al., 2018). They provide an estimate of how far an individual leave-one-out posterior distribution is from the full distribution, indicating the “importance” of each sample (i.e. each species) to the posterior. A low Pareto-*K* score (<0.3) indicates that removing the sample has no measurable effect on the posterior predictions, while a high Pareto-*K* score (>0.7) indicates that removing a sample has a large effect on the posterior (Vehtari et al., 2017). High Pareto-*K* values indicate overfitting or model misspecification, which both result in models with low predictive ability. Third, we calculated the accuracy of our models for each food item over the entire set of posterior samples. To do this, we extracted the predicted rank for each species per food item from 1000 posterior draws and used the LOO model weights to model-average the predictions over the 5 models for each food item. We then calculated the difference between the posterior mean predicted food item importance ranks and our empirical importance ranks. We calculated two accuracy scores for each food item; one for a difference of zero (posterior mean model-averaged rank = empirical rank) and one for a difference of no more than 1 (0 < |empirical rank - posterior mean model-averaged rank| *≤* 1).

### Predicting food item importance in data-deficient extant and fossil carnivorans

A desirable outcome of any predictive model is the estimation of response variables for observations of an unknown state. We first used our fitted models to predict the importance of each food item to the overall diet for the 11 data-deficient extant taxa in our data set. To further explore the predictive abilities of our models, we generated model-averaged estimates of food item importance for seven fossil carnivorans (Table 2). Molar scans and measurements were obtained from specimens housed in the fossil mammal collections of the Field Museum of Natural History, Chicago, while species mean body mass estimates were taken from the literature (Table 2). The selected fossil taxa span a phylogenetic, body size, and putative ecological breadth that should challenge our models to varying degrees. For example, the sabertoothed felid, *Smilodon fatalis*, and nimravids, *Dinctis felina* and *Hoplophoneus primaevus*, might be expected to exhibit dietary similarities to large extant felids due to their high degree of morphological similarity. Likewise, the paleomustelid *Promartes lepidus* is morphologically comparable to crown group representatives of Mustelidae and so may be expected to exhibit a similar dietary profile. In contrast, the cave bear *Ursus spelaeus* is closely related to the living brown and polar bears but, on the basis of its craniodental morphology (van Heteren et al., 2014), tooth wear patterns (Jones and DeSantis, 2016; Peigné et al., 2009; Pinto-Llona, 2013), dental topographic metrics (Pérez-Ramos et al., 2020), and nitrogen isotopic evidence (Bocherens, 2019; Hilderbrand et al., 1996; Naito et al., 2020; Richards et al., 2008; Robu et al., 2013) has been inferred to be a tough plant specialist, generalized omnivore, or seasonal bone cracker. Finally the daphoenine amphicyonid *Daphoenus* and the stem carnivoran *“Miacis” latidens* have no close extant relatives for comparison. Including these taxa in our sample presents an opportunity to gain new insights into their paleoecology.

**Table 2:**
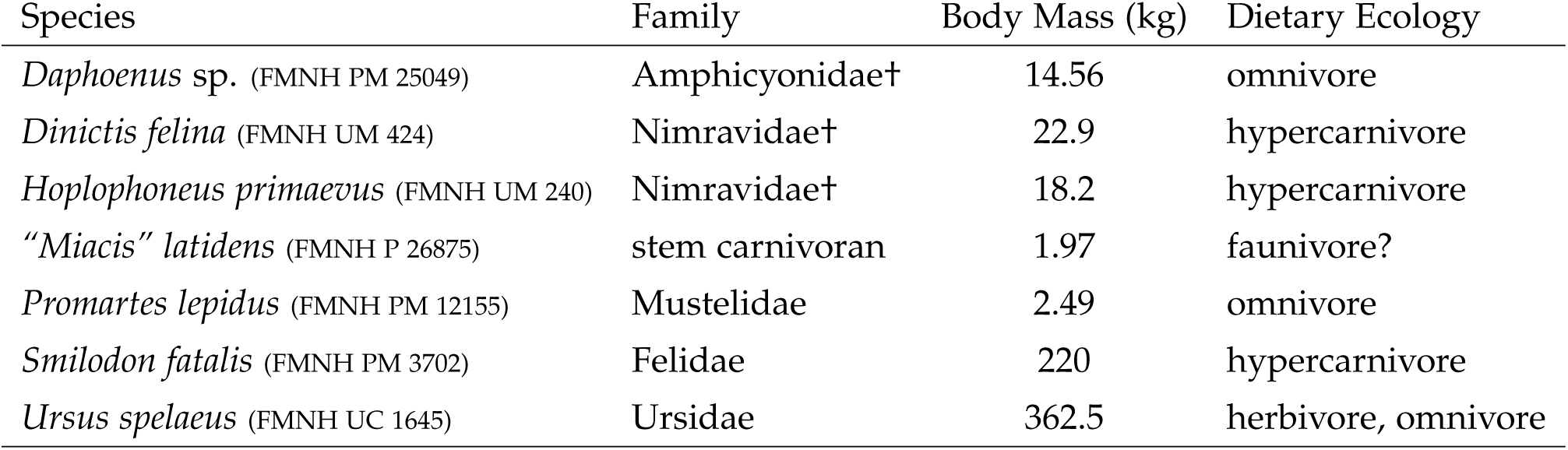
Fossil Taxa Sampled in this Study.

For each model with a non-zero LOO weight, we used the dental and body mass data from each extinct and data-deficient taxon to sample 4000 posterior draws of the expected value of their response variable (diet rank) from the posterior distribution for each food item for each species. Posteriors were multiplied by the LOO model weights and then summed to generate 4000 model-averaged posterior draws. Predictions from these models do not come in the form of a point estimate of the importance rank for each food item for each taxon, as one would obtain from a more traditional discriminant function analysis but, rather, as a vector of probabilities where the *ith* value is the probability that the dietary item has an importance rank *i* or lower to the diet of that taxon. Therefore, for each draw, we computed a Weighted Importance Score (WIS) for the food item as

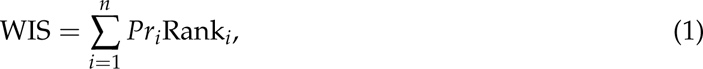

where *Pr_i_* is the probability of the *i*th of the *n* ranks, resulting in 4000 Weighted Importance Scores for each item for each species. The WIS values are continuous on the interval 1-4, rather than integer-valued as in the ordinal dietary data from extant data-rich species. To aid comparison, we performed the same series of steps to generate a distribution of 4000 WIS values for each food item for each data-rich extant species in our sample.

For each draw in our 4000 posterior estimates, we performed a principal components analysis on the correlation matrix of WIS values for data-rich extant species and projected data-deficient extant and extinct species into the resulting space to identify their nearest data-rich neighbors, which we interpret to be the closest dietary analogs. We calculated the frequency with which each data-rich extant species was recovered as the nearest neighbor to each data-deficient and extinct taxon in the 4000 posterior samples, which we interpret as the posterior probability of that data-rich extant taxon being the true nearest neighbor.

## Results

### Morphological Data

Principal components analysis of the dental topographic data yields patterns that conform to functional expectations (Figure 1). PC1 explains 64% of the variance in dental topographic data and separates taxa with large relative grinding areas, complex molars, and tall cusps (negative scores) from those with small grinding areas, relatively simple teeth, and first molars with high relief (positive scores). Qualitatively, this axis appears to separate more “omnivorous” taxa (e.g. bears, negative scores) from more “carnivorous” taxa (e.g. felids, hyaenids, *Cryptoprocta*, positive scores). PC2, which explains 18% of the variance in tooth shape data separates taxa with complex molars (negative scores), from those with sharp molars and high relief (positive scores). This axis seems to be associated with the degree of insectivory, with small insectivorous taxa (herpestids and euplerids, positive scores) separating from taxa with less reliance on insects (negative scores). Broken stick analysis suggests that only the first PC is significant. There is a moderate degree of phylogenetic signal in carnivoran molar morphology that is significantly different from random expectation (*K_mult_*=0.58, *P*=0.001, Effect Size=13.11).

**Figure 1:**
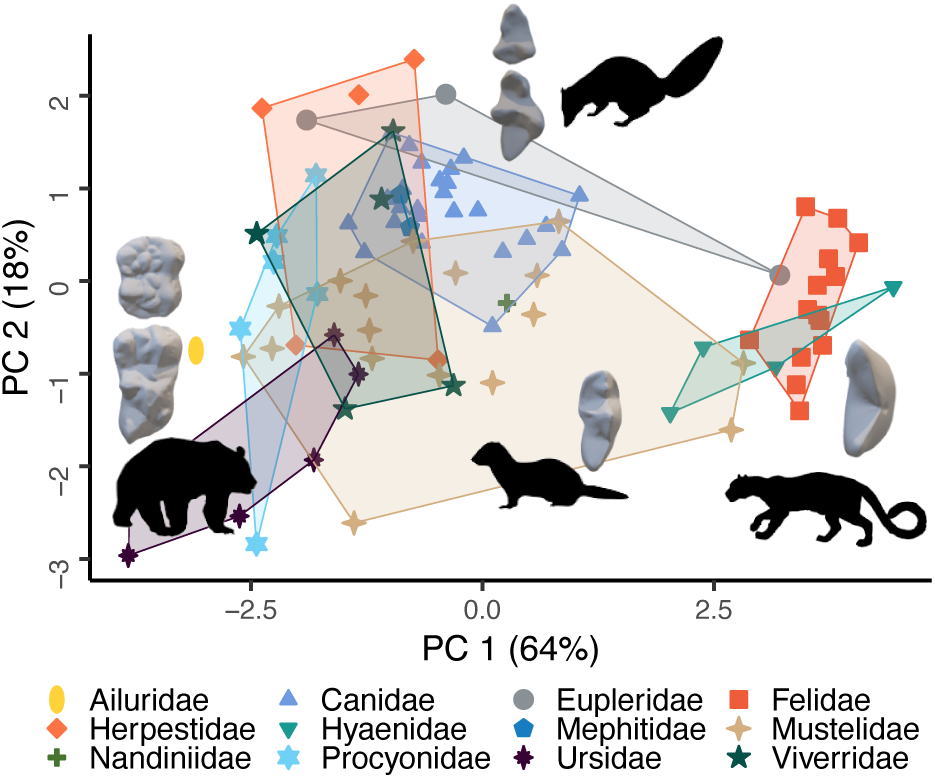
Phylogeny accounts for only a portion of dental morphology. Dental morphology contains some phylogenetic signal (K_mult_ = 0.58) as carnivoran families are loosely organized in morphospace. Families with narrow, sharp m1 molars and no m2 molars cluster in the positive values of PC1, while families with large, complex molars have negative PC1 values. Species’ silhouettes are from PhyloPic and are available under a Public Domain license.

### Dietary Data

Of the 99 species for which we collected morphological measurements, 11 lacked sufficiently detailed descriptions of their dietary habits to score important rankings for each food item. These data-deficient taxa were therefore used to generate predictions (see below). Figure 2 shows the dietary space defined by the first two polychoric PCs of food-item importance scores from the remaining taxa, with plotting symbols corresponding to discrete dietary groupings from four commonly used categorization schemes. The first PC, which accounts for 27.73% of the variance in the data, separates taxa for which large and, to a lesser degree, small mammals are an important component of the diet (positive scores) from those in which hard and soft invertebrates and fruits (negative scores) are important. The second PC, which accounts for 18.07% of the variance, separates taxa that place high importance on small vertebrates (small mammals, birds, herptiles), as well as eggs and carrion (positive scores) from those that consume more plants (negative scores).

**Figure 2:**
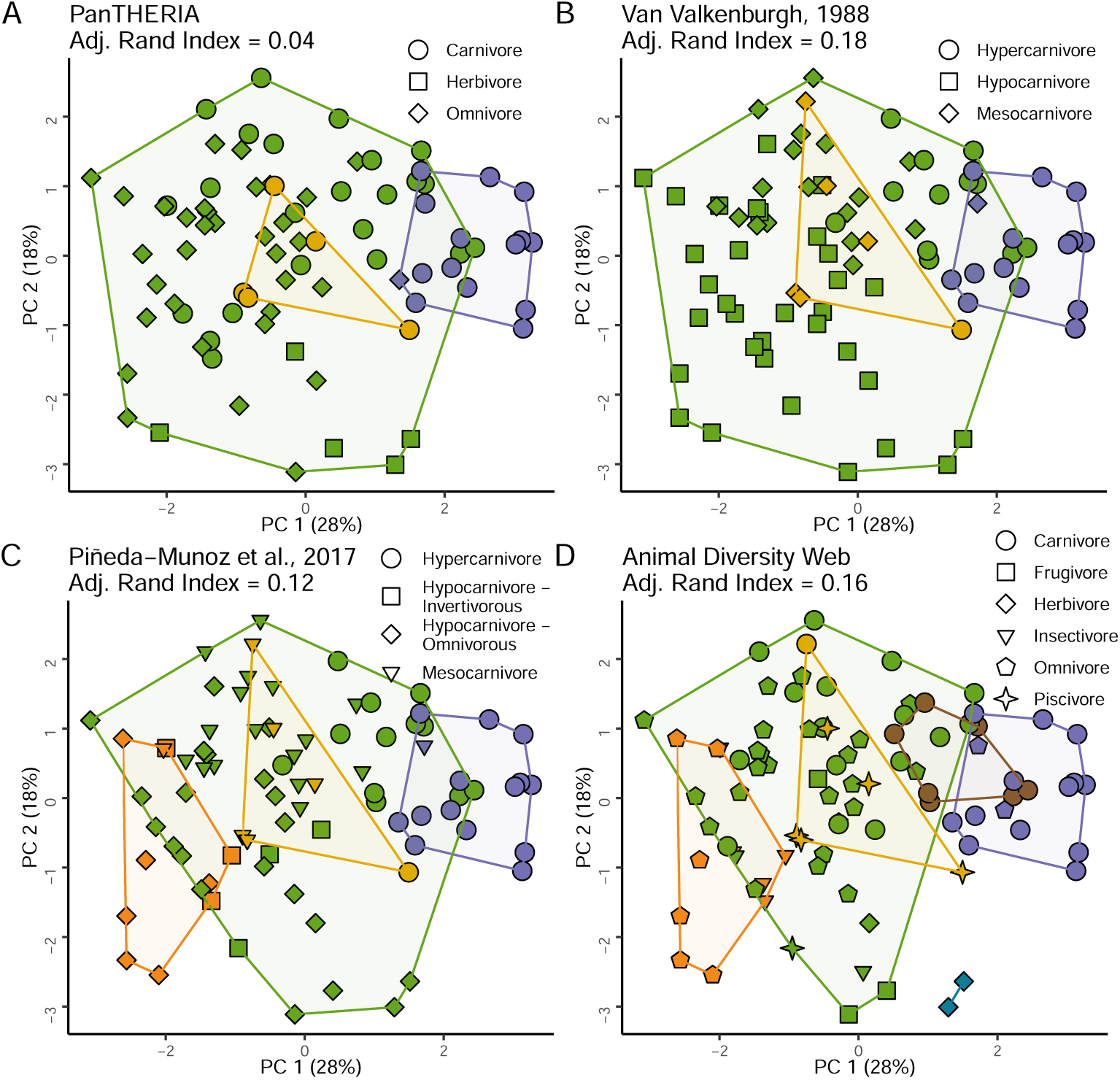
A cluster analysis of polychoric PCA scores generated from multivariate dietary importance data captures some discrete diet categorizations, but misses most. In each plot, the symbols represent the discrete categories assigned by the four *a priori* categorization schemes, the colors represent the multivariate dietary clusters delineated using mixture models in mclust, and the adjusted Rand Index value indicates the match between the two. The number of mclust-delineated diet clusters in each plot was set to the number of groups in the *a priori* categorization schemes **A:** The simplistic *k*=3 category scheme of the PanTHERIA database is an especially poor descriptor of the multivariate diet, with an adjusted Rand Index of 0.04. This mismatch can be visualized by the number of different symbols that share the color green. **B:** The carnivoran-specific *k*=3 scheme of Van Valkenburgh (1988) identifies many of the “hypercarnivore” species cluster (in purple), but the remaining two clusters do not match the “hypocarnivore” or “mesocarnivore” categories. **C:** The *k*=4 *a priori* categories of Pineda-Munoz et al. (2017) split hypocarnivores into two groups, however, these two categories, represented by the square and diamond shapes, are not well separated in multivariate diet space. **D:** The mixture models correctly delineate most of the piscivorous (stars) and herbivorous (diamonds) species from the *k*=6 categorization scheme of Animal Diversity Web, though the remaining four *a priori* categories are not well delineated.

Existing dietary classifications do not appear to conform to multivariate diet data. Mixture-model clustering analyses using the full set of 13 PCs identified a 5 cluster scheme as optimal, based on BIC, though 5 clusters were only minimally preferred (BIC score difference <-2.0) to 3, 4, or 6 clusters, and there is a weak correspondence between the four *a priori* discrete dietary classification schemes and cluster membership when the number of clusters is fixed to be equivalent (Figure 2). The simple PanTHERIA scheme – carnivore, omnivore, herbivore – performed especially poorly (Figure 2A), with an adjusted Rand Index of 0.04. The three remaining schemes match the multivariate dietary clusters slightly better, but adjusted Rand Indicies remain low (*≤* 0.18; Figure 2B–D). Although the *k*=6 clustering analysis does not perform well overall, it does effectively collate 5 of the 6 “piscivorous” species into a cluster, most evident on PC5, and places 2 of the 3 “herbivorous” species into a cluster (Figure 2D).

### Ordinal Models of Food Types

Chains from all Bayesian ordinal models demonstrated appropriate convergence (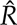 < 1.002). Prior predictive simulations showed that different response distributions (normal, student-*T*, and Dirichlet) each performed best for some food items (Figure S1). All model averaged Pareto-*K* scores are all in the acceptable (0.55 - 0.3, 4% of samples) to good (<0.3, 96% of samples) range (Table S2), indicating that our models are not overfit and have reasonable predictive accuracy. Most of the Pareto-*K* values >0.3 are from the models for root consumption, likely because only 8 of the 88 species in our sample are known to eat any roots, and there are no dental or body mass signatures of root feeding in the diet. Detailed model outputs and visualizations are in Figure S3 and Table S3. Calculations of model weights using LOO scores and model stacking showed that different dental topographic metrics best predict the dietary importance of different food types (Table 3). However, these relationships are often mediated by body size and sometimes in very different ways. To visualize these effects, we plotted the posterior probability of belonging to each dietary importance rank for values of the optimal trait, broken down into low (−1.5 *sd*), mean, and high (+1.5 *sd*) body mass groupings for select dietary items (Figure 3). RLGA strongly decreases with increased consumption of birds (Figure 3A), large mammals (Figure 3D), and small mammals (Figure 3E), but increases with increased consumption of fish. However, due to the strong interaction between mass and RLGA, a diet containing a large proportion of birds is best predicted in small-bodied carnivores (Figure 3A), a diet rich in small mammals is best predicted in small and medium-sized carnivores (Figure 3E), and a diet rich in large mammals is best predicted in large carnivores (Figure 3D), while high importance of fish is best predicted for large-bodied taxa with large grinding areas.

**Figure 3:**
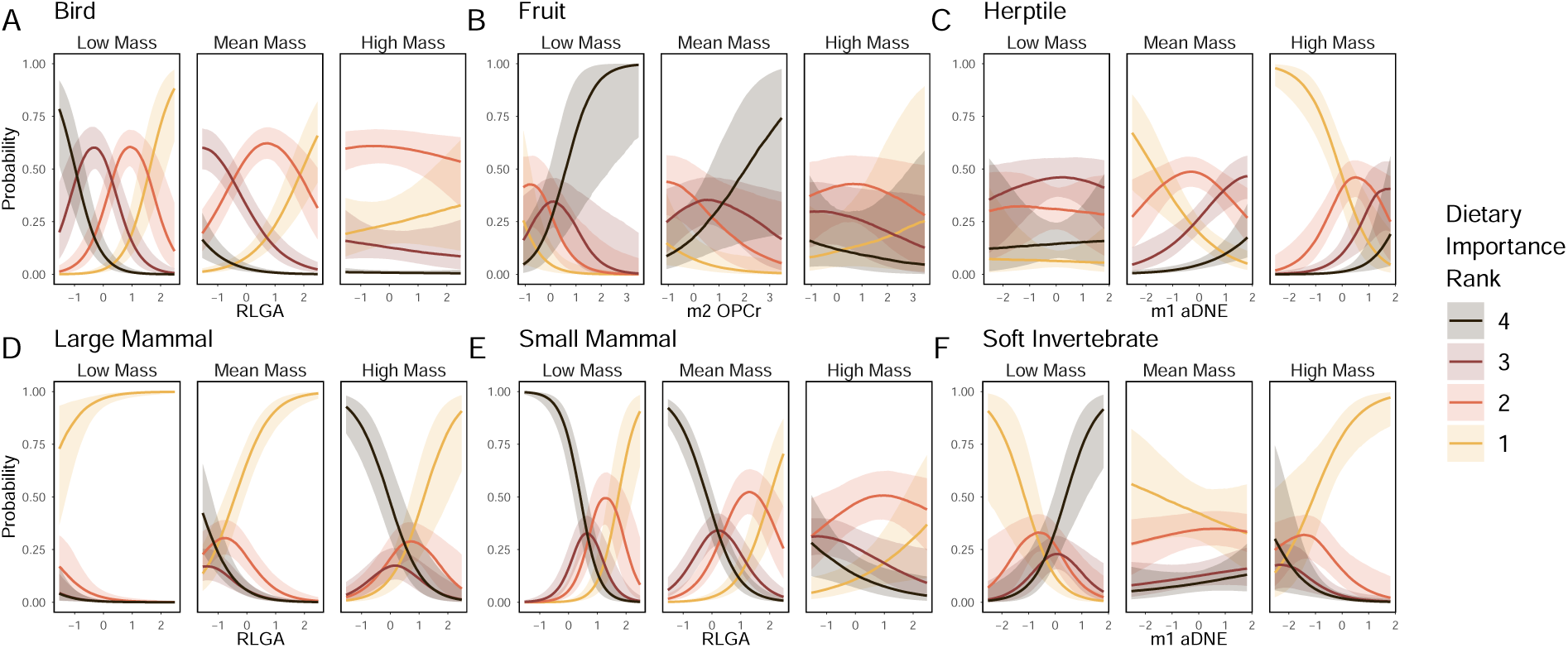
The influence of dental morphology, mass, and their interaction on dietary prediction strongly varies between dietary items. Plots depict the 80% prediction intervals for each of the 4 dietary ranks at low mass, average mass, and high mass. RLGA measurements can effectively predict the amount of bird (**A**) and small mammal (**E**) in the diet of small-bodied carnivorans, however, these predictions become much less effective for large-bodied species. However, RLGA can effectively predict the amount of large mammal consumed across all body sizes (**D**). m1 DNE effectively predicts herptiles and soft invertebrates in the diet of large carnivorans, however, it is more difficult soft invertebrates for average-sized species and herptiles for smaller species (**C**, and **F**). A high m2 OPCr value in small carnivorans means a high probability of eating fruit, though the predictive power decreases with increasing mass. (**B**).

**Table 3:**
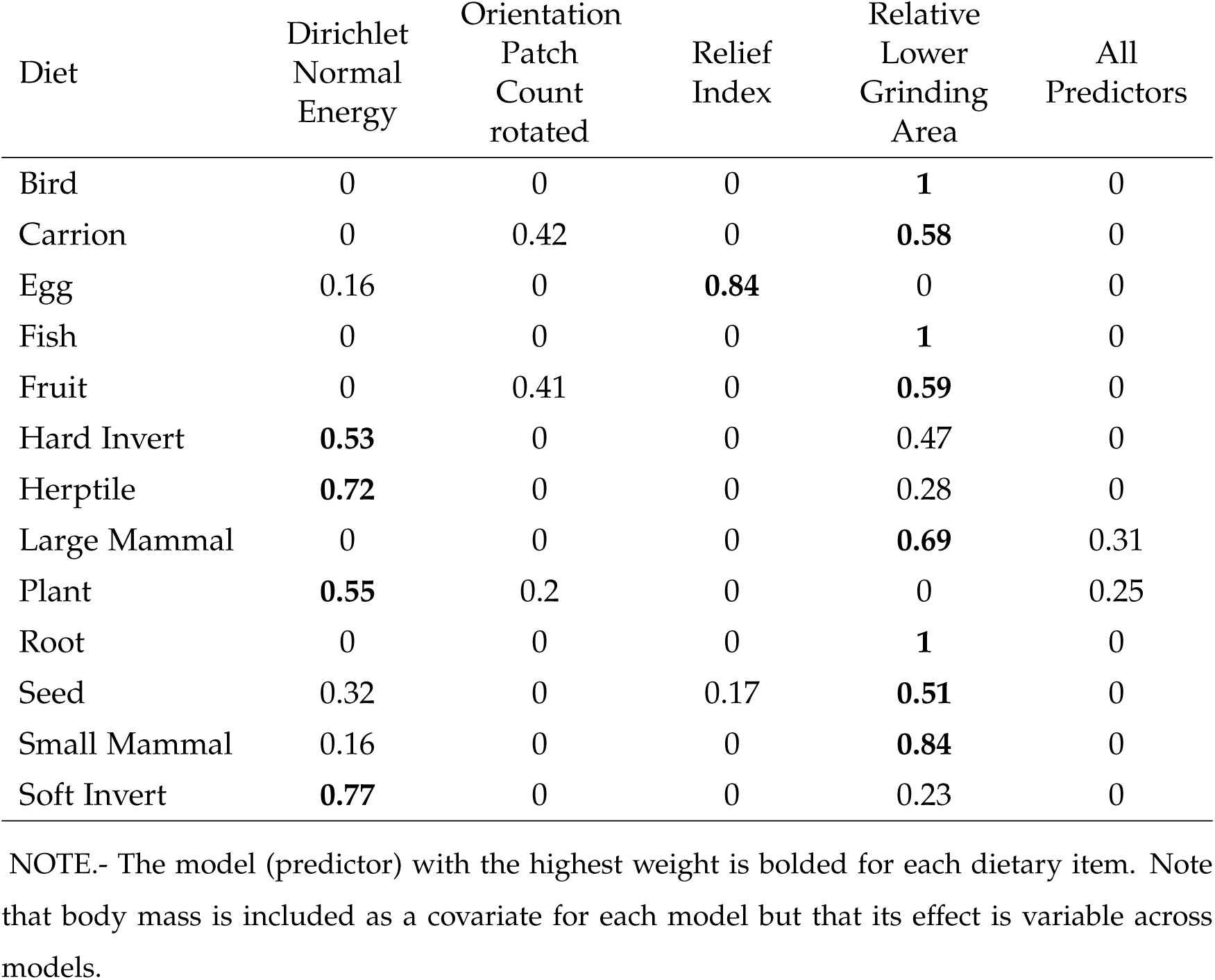
Model Support (LOO weight from model stacking) for Ordinal Regressions of Dietary Item Importance Ranks (rows) on Dental Traits (columns).

| Diet | Dirichlet<br>Normal<br>Energy | Orientation<br>Patch<br>Count<br>rotated | Relief<br>Index | Relative<br>Lower<br>Grinding<br>Area | All<br>Predictors |
| --- | --- | --- | --- | --- | --- |
| Bird | 0 | 0 | 0 | **1** | 0 |
| Carrion | 0 | 0.42 | 0 | **0.58** | 0 |
| Egg | 0.16 | 0 | **0.84** | 0 | 0 |
| Fish | 0 | 0 | 0 | **1** | 0 |
| Fruit | 0 | 0.41 | 0 | **0.59** | 0 |
| Hard Invert | **0.53** | 0 | 0 | 0.47 | 0 |
| Herptile | **0.72** | 0 | 0 | 0.28 | 0 |
| Large Mammal | 0 | 0 | 0 | **0.69** | 0.31 |
| Plant | **0.55** | 0.2 | 0 | 0 | 0.25 |
| Root | 0 | 0 | 0 | **1** | 0 |
| Seed | 0.32 | 0 | 0.17 | **0.51** | 0 |
| Small Mammal | 0.16 | 0 | 0 | **0.84** | 0 |
| Soft Invert | **0.77** | 0 | 0 | 0.23 | 0 |

NOTE.- The model (predictor) with the highest weight is bolded for each dietary item. Note that body mass is included as a covariate for each model but that its effect is variable across models.

We also found that the significance of first and second molar morphology varied across food types. Across the range of carnivoran body mass, a high DNE value on m2, but not m1, increases the probability of plants in the diet. OPCr of the m2, but not m1, is positively associated with the importance of fruit in small carnivorans but here, due to the interaction with mass, high m2 OPCr values are negatively associated with fruit consumption in larger carnivorans (Figure 3B). Similarly, high DNE values for m1 are associated with high importance of soft invertebrates in small-bodied carnivorans, while low DNE values are predictive of high importance of soft invertebrates in large-bodied taxa (Figure 3F). Other models are more complex still. Small-bodied carnivores consume more hard invertebrates than larger species, especially in species with high m1 DNE values. However, an increased lower grinding area is strongly associated with increased hard invertebrate consumption across all body sizes.

Rarer food types present a range of challenges to our predictive models. Specialization on herptiles is uncommon in carnivorans but the best fitting model, incorporating DNE and body mass (Table 3), finds a moderate probability that reptiles and amphibians are important to the diet of small-bodied taxa, regardless of the sharpness of m1 and m2 (Figure S3, Table S3). Infrequent herptile consumption in the empirical diet data therefore lowers the predictive accuracy of our herptile models (Table 4), though precise predictions (exact matches between empirical ranks and posterior mean model-averaged predictions) still remain above 50%. Similarly, while no carnivorans in our data set frequently consume roots, the probability of consuming at least some roots is inferred to robustly increase with increased lower grinding area, particularly in larger species (Figure S3, Table S3). A different effect is seen for plants. Although true plant specialists (i.e. an importance rank for plants of 4) exist in our dataset (e.g., the pandas *Ailuropoda melanoleuca* and *Ailurus fulgens*), they are rare and most taxa use plants modestly. This leads to underprediction of plant importance in these specialist taxa (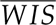 = 3.11 and 2.75, respectively) and overprediction of importance in others, including the more carnivorous polar bear *Ursus maritimus* (empirical plant importance rank = 2, (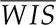 =2.78).

**Table 4:**
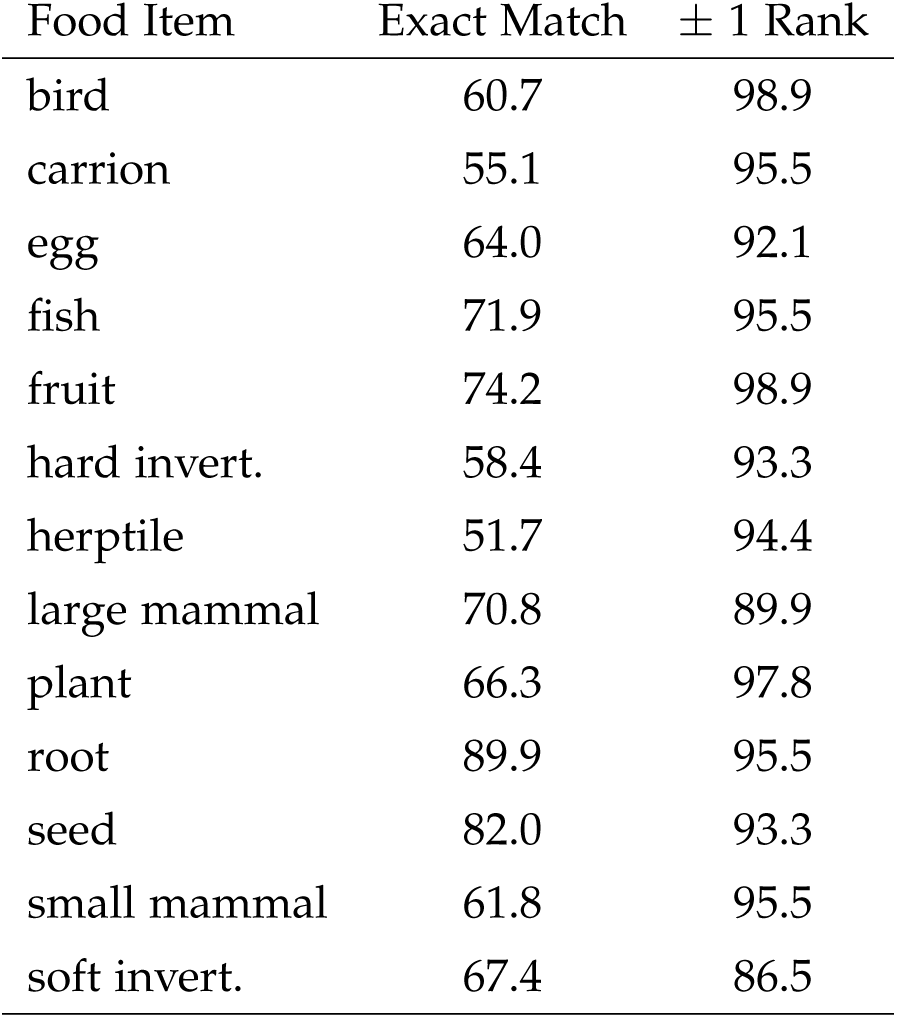
Predictive Accuracy of Ordinal Regression Models of Dietary Importance Ranks

| Food Item | Exact Match | $\pm 1$ Rank |
| --- | --- | --- |
| bird | 60.7 | 98.9 |
| carrion | 55.1 | 95.5 |
| egg | 64.0 | 92.1 |
| fish | 71.9 | 95.5 |
| fruit | 74.2 | 98.9 |
| hard invert. | 58.4 | 93.3 |
| herptile | 51.7 | 94.4 |
| large mammal | 70.8 | 89.9 |
| plant | 66.3 | 97.8 |
| root | 89.9 | 95.5 |
| seed | 82.0 | 93.3 |
| small mammal | 61.8 | 95.5 |
| soft invert. | 67.4 | 86.5 |

NOTE.- The accuracy of the model predictions varies among food items, but is consistently high. Most models estimate within 1 rank of the empirical importance rank over 90% of the time. To generate these estimates, we took 1000 draws from the model-averaged posterior distributions from each food item, then calculated the accuracy of each predicted rank. These values are the mean accuracy scores of the 1000 draws.

### Predicting Dietary Importance in Data-deficient Extant and Fossil Carnivorans

Dietary importance predictions for the 11 data-deficient extant taxa are consistent with limited knowledge regarding their diets. Most data-deficient taxa are small-bodied members of clades that are traditionally considered to exhibit more generalized diets (e.g. Herpestidae, Viverridae), and posterior mean Weighted Importance Scores (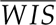) reflect this, with non-negligible contributions (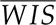 *>* 2.0) of birds, hard and soft invertebrates, fruits, herptiles, and small mammals inferred for most (Figure 4A). Carrion, eggs, fish, plants, roots, and seeds are identified as relatively low-importance foods (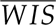 *<* 2.0) for most of these taxa. Among more specialized taxa, the bay cat *Catopuma badia* and marbled cat *Pardofelis marmorata* are confidently predicted to be obligate small mammal feeders (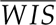 *>* 3.9) and to include a large proportion of birds in their diet (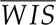 *>* 3.2). These two cats are also the only taxa for which large mammals are inferred to ever be consumed, though the importance of this food resource is low (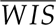 2.1-2.3). The cat-like banded linsang *Prionodon linsang* is predicted to be similarly reliant on small mammals (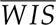 = 3.91) but, unlike the cats, hard (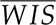 = 3.45) and soft (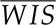 = 3.09) invertebrates are also inferred to comprise important components of its diet. Soft invertebrates are predicted to feature heavily (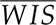 *>* 3.0) in the diets of banded palm civet *Hemigalus derbyanus* and Sunda stink badger *Mydaus javanensis*, as well as the short-tailed mongoose *Herpestes brachyurus*. Due to the small body size of many of these data-deficient taxa, predictions for the importance of some food items, herptiles, in particular, were relatively uniform across species.

**Figure 4:**
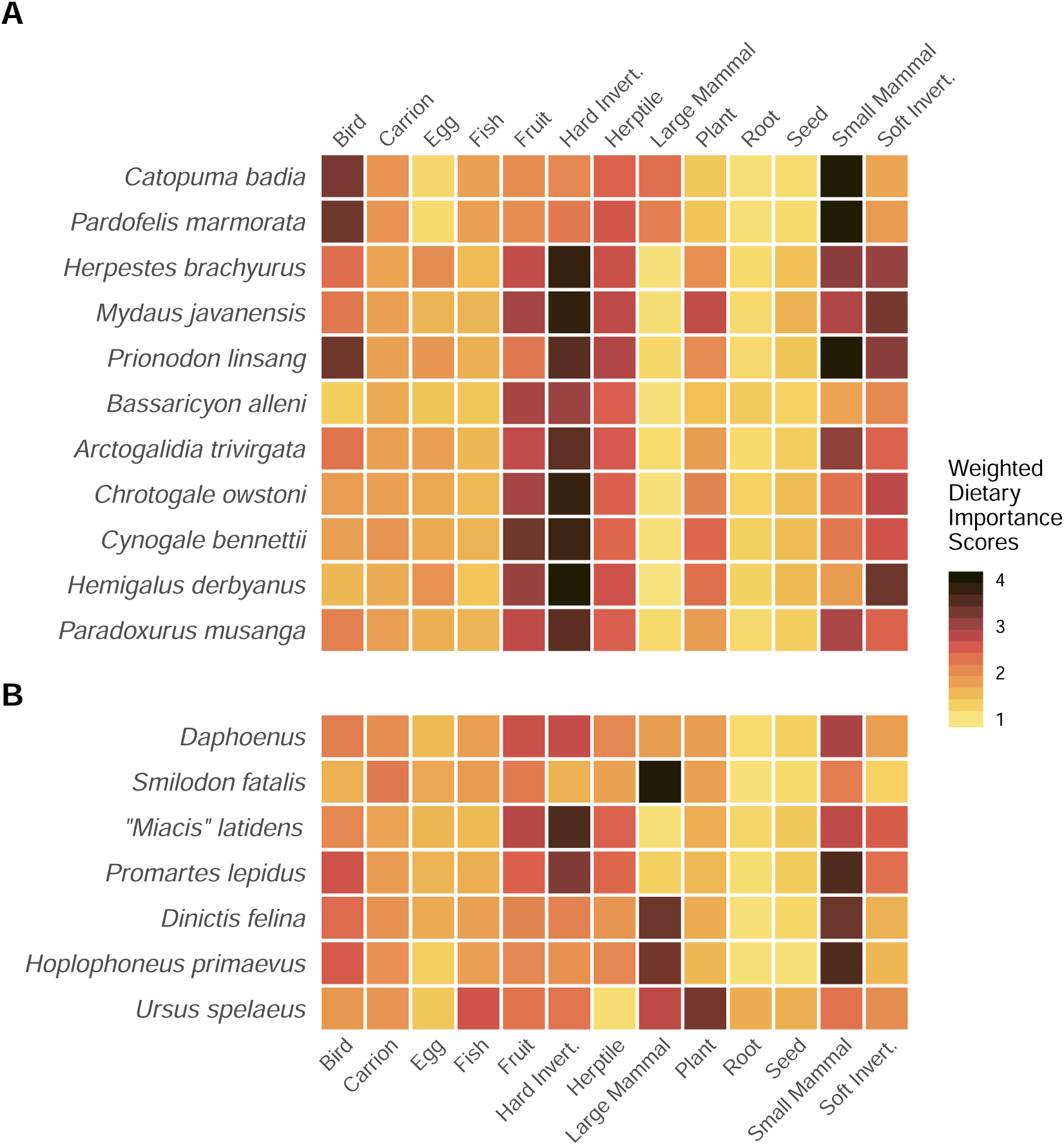
Ordinal models of dietary items can be used to predict the diet of understudied or extinct species. Estimates of the diet rankings of **A**) 11 carnivoran species that lack robust dietary data, and **B**) 7 fossil taxa use the models for each dietary item, weighted by the LOO stacking scores, and the data-deficient species’ dental metrics and mass as input. Heatmap reports the posterior mean weighted importance score (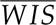, see text for calculation). Scores range continuously between 1, meaning that an item is predicted to never be consumed, and 4, where a food item is of primary importance to the diet of that species. Species are ordered by Family (See Table 5). Some dietary items, such as bird, small mammal, and soft invertebrate, have heterogeneous predictions across species, demonstrating that molar morphology and mass influence these predictions. Other diet items such as egg, root, and carrion have similar and ambiguous predictions across all species, suggesting a low predictive power for these taxa.

**Table 5:** The three most common Nearest Neighbors for Data Deficient Extant Taxa and Fossil Taxa in Multivariate Diet Space.

| Taxon | Family | Neighbor 1 | Neighbor 2 | Neighbor 3 |
| --- | --- | --- | --- | --- |
| *Catopuma badia* | Felidae | *Leopardus wiedii* 15% | *Leptailurus serval* 14% | *Felis silvestris* 14% |
| *Pardofelis marmorata* | Felidae | *Prionailurus bengalensis* 17% | *Leopardus wiedii* 15% | *Leopardus serval* 14% |
| *Herpestes brachyurus* | Herpestidae | *Vulpes zerda* 13% | *Mephitis mephitis* 12% | *Bassariscus astutus* 11% |
| *Mydaus javanensis* | Mephitidae | *Bassariscus astutus* 23% | *Viverricula indica* 20% | *Conepatus mesoleucus* 9% |
| *Prionodon linsang* | Prionodontidae | *Spilogale putorius* 17% | *Vulpes zerda* 14% | *Vulpes velox* 11% |
| *Bassaricyon alleni* | Procyonidae | *Bdeogale jacksoni* 28% | *Potos flavus* 21% | *Arctictis binturong* 13% |
| *Arctogalidia trivirgata* | Viverridae | *Conepatus mesoleucus* 7% | *Hydrictis maculicollis* 6% | *Atilax paludinosus* 6% |
| *Chrotogale owstoni* | Viverridae | *Otocyon megalotis* 13% | *Ichneumia albicauda* 10% | *Procyon lotor* 9% |
| *Cynogale bennettii* | Viverridae | *Otocyon megalotis* 16% | *Nasua nasua* 14% | *Procyon lotor* 13% |
| *Hemigalus derbyanus* | Viverridae | *Nasuella olivacea* 37% | *Nasua nasua* 17% | *Bdeogale jacksoni* 16% |
| *Paradoxurus musanga* | Viverridae | *Lycalopex vetulus* 10% | *Paguma larvata* 9% | *Conepatus mesoleucus* 8% |
| *Daphoenus* | Amphicyonidae | *Arctictis binturong* 14% | *Canis aureus* 7% | *Canis adustus* 7% |
| *Smilodon fatalis* | Felidae | *Panthera leo* 46% | *Crocuta crocuta* 23% | *Canis lupus* 10% |
| *"Miacis" latidens* | "Miacidae" | *Conepatus mesoleucus* 10% | *Paguma larvata* 9% | *Lycalopex vetulus* 8% |
| *Promartes lepidus* | ?Mustelidae | *Vulpes velox* 9% | *Nandinia binotata* 9% | *Atilax paludinosus* 8% |
| *Dinictis felina* | Nimravidae | *Lycaon pictus* 18% | *Cuon alpinus* 13% | *Uncia uncia* 8% |
| *Hoplophoneus primaevus* | Nimravidae | *Neofelis nebulosa* 13% | *Lycaon pictus* 13% | *Cuon alpinus* 11% |
| *Ursus spelaeus* | Ursidae | *Ursus maritimus* 41% | *Ailuropoda melanoleuca* 22% | *Tremarctos ornatus* 11% |

NOTE.- Neighbors are listed in order by the percentage of nearest neighbor matches from 4000 posterior predictions (in parentheses).

Projecting data-deficient extant species into the multivariate diet space for data-rich species illuminates dietary affinities between taxa that transcend phylogenetic and biogeographic boundaries. The “cat-like” linsang does not cluster with felids at all, but is closest to the spotted skunk (*Spilogale putorius*), as well as a selection of small canids, mustelids, euplerids, and herpestids (latter three not shown in Table 5), that are united in placing high importance on a combination of small vertebrate and invertebrate prey. Among other results, it is notable that the nearest neighbors of the three Southeast Asian hemigaline viverrids (*Chrotogale*, *Cynogale*, and *Hemigalus*) largely come from the American Procyonidae (raccoons and relatives). Another noteworthy feature is that the density of the occupied region of dietary space can be inferred from the number and frequency of nearest neighbors in the posterior sample. For example, the hooded skunk *Conepatus mesoleucas* is most frequently recovered as the nearest neighbor to the small-toothed palm civet *Arctogalidia trivirgata* but with a posterior probability of only 0.07. With numerous other taxa inferred as nearest neighbors at posterior probability *<*0.1, *A. trivirgata* apparently resides in a densely occupied region of diet space. In contrast, *Hemigalus derbyanus* appears to reside in a far less occupied region of diet space, close to the two coati species (> 50% of total matches).

Model-averaged predictions for the relative importance of different food items in the diet of some extinct carnivorans (Figure 4B) are largely consistent with previous work. We infer that large mammals were an extremely important (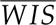 = 3.92) component of the diet of the machairodont felid *Smilodon fatalis* and the nimravids *Dinictis felina* (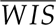 =3.32) and *Hoplophoneus primaevus* (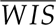 =3.27). Small mammals are inferred to have been of comparable importance to large mammals in the diets of the two nimravids (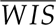 = 3.31 and 3.53, respectively), but not for *Smilodon* (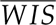 = 2.12). Other animal-derived protein sources, such as fish and herptiles, may have been occasionally consumed by these taxa, but they are generally inferred to be less important (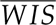 *<* 2.5).

The amphicyonid *Daphoenus* is inferred to have had a rather broad diet in which no single food item dominated but where small mammals (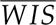 = 2.88), hard invertebrates (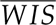 = 2.66), and fruits (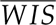 = 2.62) were all consumed frequently. Plants are predicted to have been the most important component of the diet of the cave bear *Ursus speleaus*, with a 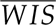 of 3.27 that exceeds that of the the extant bamboo specialist pandas. However, we also infer that fish (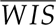 = 2.56) and large mammals (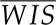 =2.72) may have been regularly consumed, with fruits (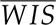 =2.25), hard invertebrates (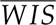=2.22) and small mammals (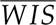 =2.25) all consumed at least occasionally.

The paleodiet of the two other taxa in our sample has not been as thoroughly studied. We infer that *“Miacis” latidens*, a small-bodied stem carnivoran from the Eocene of North America, relied on hard-bodied invertebrates (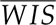 =3.53), with fruit, soft invertebrates, and small vertebrates also contributing occasionally (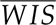 = 2.03-2.77) to its diet. *Promartes lepidus* is inferred to have primarily consumed small mammals (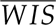 =3.53), though herptiles, birds, fruit, and invertebrates all yield non-negligible probabilities of being consumed at least occasionally (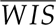 = 2.28-2.60).

Projecting fossil species in the multivariate diet space of extant carnivorans again yields a mix of intuitive and surprising results. The lion *Panthera leo* and the spotted hyaena *Crocuta crocuta*, both of which place high importance on large mammal prey, are inferred to be nearest neighbors to the large sabertoothed cat *Smilodon*. The nimravids are most frequently recovered closest to two hypercarnivorous canids, the African wild dog (*Lycaon pictus*), and dhole (*Cuon alpinus*), and two smaller pantherine felids that are united in placing similar importance on large and small mammalian prey. *Daphoenus* clusters close to extant canids and mustelids that tend to exhibit more generalized diets, though its most frequent nearest neighbor is a frugivorous viverrid, the binturong *Arctictis bintutong*. However, the posterior probabilities that any of these taxa are the nearest dietary neighbor to *Daphoenus* are much lower (PP = 0.7-0.14) than for the sabertoothed taxa (PP = 0.1-0.46), indicating that the amphicyonid probably occupied an area of diet space that is more densely packed by extant species than did the sabertooths. “*Miacis*” *latidens* and *Promartes lepidus* fall closest to a suite of small-bodied carnivorans spanning the families Canidae, Herpestidae, Mephitidae, Mustelidae, and Viverridae. The most frequent neighbors of “*Miacis*” *latidens* are taxa that are typically considered “insectivorous” or “generalists”, while the most frequent neighbors of *Promartes* are small taxa that tend to specialize on small vertebrate prey. However, the low frequencies of the top three neighbors (0.08-0.1) again indicate occupation of a densely packed region of diet space.

The cave bear yields the most unexpected set of nearest neighbors. Despite plants being the most important food item based on weighted importance scores, the most carnivorous extant bear, the polar bear *Ursus maritimus*, is recovered as the nearest neighbor to the cave bear in 41% of posterior samples. The giant panda *Ailuropoda melanoleuca*, a taxon typically considered to be strictly herbivorous, is the next most frequent nearest neighbor, being recovered as such in 22% of posterior samples. That the two most frequently recovered nearest neighbors are so ecologically disparate, yet account for a combined 63% of the posterior sample, may suggest that the *Ursus spelaeus* occupied a unique region of diet space, relative to extant carnivorans. However, the difficulty of accurately predicting the importance of plants to the diet of carnivorans based on our sample means that this interpretation should be made cautiously, especially as the relatively herbivorous Andean bear *Tremarctos ornatus* rounds out the top 3 most probable nearest neighbors with a posterior probability of 0.11.

## Discussion

Diet is a fundamental life history trait, defining an organism’s basic biology as well as its role in a community. However, diet is also a complex, multidimensional trait, and efforts to condense the diversity and frequency of food consumption into discrete dietary categories necessarily omit critical information regarding the realized dietary composition of a given species. We have demonstrated that dietary item importance rankings, phenotypic traits associated with food processing, and Bayesian multilevel ordinal modeling can be leveraged to validate the relationship between traits and food item importance, and to predict the dietary composition of extinct and understudied extant taxa without the need to condense multivariate dietary data into one of a few discrete categories. Our findings illuminate a nuanced perspective on dietary diversity by demonstrating the true multivariate nature of individual species’ diets, while simultaneously revealing the loss of important ecological and biological information that follows the discrete categorization of complex traits. Additionally, our results highlight how the complex interactions between focal dietary traits, such as molar shape and structure, and peripheral traits, such as body mass, are critical to fully understanding a species’ dietary niche.

### Diet is a Multivariate Trait and Should Be Analyzed as Such

The question of how to best quantify or categorize mammalian diet has lived long in the literature. Simple classification schemes that crudely reflect trophic level (carnivore, omnivore, herbivore) continue to be widely used in comparative studies (e.g., Fabre et al., 2017; Price and Hopkins, 2015; Price et al., 2012; Rowe et al., 2016; Santana et al., 2011; Evans et al., 2007), despite an awareness that two species in the same category may use dietary items of very different sizes, material properties, nutritional qualities, and phylogenetic affinity (Pineda-Munoz and Alroy, 2014). Eisenberg (1981) provided one of the first attempts to finely describe the full range of diversity in mammalian feeding behavior using a classification scheme with 16 states, each of which was based on a dominant food item (i.e. a specialization). These categorical states have been further refined over time by workers specializing on more restricted clades, each of which may exhibit its own range of unique predatory and dietary behaviors (Fulwood et al., 2021; Kienle et al., 2017; Slater et al., 2010; Toljagić et al., 2018; Verde Arregoitia and D’Eĺıa, 2021; Williams and Kay, 2001; Boyer, 2008; Slater, 2015; Van Valkenburgh, 1988). Still, it is apparent that most mammals make use of a mixture of food types and that dietary variation is more continuously distributed than the most complicated categorical classifications are able to permit (Pineda-Munoz and Alroy, 2014). Indeed, our ordination of dietary importance data for carnivorans revealed that even species traditionally categorized as “carnivores” occupy a broad swath of dietary space, and do not cluster into natural groupings (Figure 2). Attempts to project dietary variation into a single univariate, quantitative trait (e.g., López-Aguirre et al., 2022; Rojas et al., 2018) may also lead to information loss; although the first principal component of our importance scores appears to represent a carnivory-herbivory continuum, as in Rojas et al. (2018), finer-scale patterns of dietary variation, such as a dominant use of invertebrate prey or fish, occur on subsequent axes and are missed if we focus only on the first or the set of “significant” PCs.

Characterizing diet in a more natural, quantitative fashion still poses considerable challenges, particularly when expanding consideration to taxa that lack detailed ecological data. In one attempt to address this problem, Grundler and Rabosky (2020) proposed a novel comparative method in which the proportional utilization of a finite suite of resource types is modeled as a multinomial distribution that can evolve over the branches of a phylogenetic tree. Under this model, observational data on the frequency with which resources are used by a given species do not represent the diet itself but, rather, are draws from the multinomial distribution (i.e. the diet) allocated to that branch of the phylogeny, which is, in turn, estimated by the model. Using a large database of dietary observations taken from Grundler (2020), Grundler and Rabosky (2021) found evidence that snakes rapidly expanded in dietary diversity and complexity during the early Cenozoic from a likely insectivorous ancestor. While this approach holds much promise for modeling the evolution of complex traits on phylogenies of extant taxa, it appears to be of more restricted applicability to extinct lineages, where observational data on dietary item use are not typically available and “diet” must be estimated from proxies. Data-deficient extant taxa may also pose considerable problems for this method; although variation in sampling quality can be explicitly accommodated (Grundler and Rabosky, 2020), usable observational data are simply lacking for a large number of taxa (Gainsbury et al., 2018). In this respect, the more qualitative but widely applicable approach of Rojas et al. (2011) and Kissling et al. (2014) that we use here is the one that we think holds particular promise. We acknowledge that the use of keywords to rank the importance of food items is not without problems of its own, and in particular that the use of appropriate literature resources is paramount for quality control of data (Gainsbury et al., 2018). It is also likely that the items listed by Rojas et al. (2011) and Kissling et al. (2014) are too broad for describing dietary variation in some clades, where they may fail to capture fine-scale patterns of dietary niche partitioning (McNaughton et al., 1986; Pineda-Munoz and Alroy, 2014; Machado, 2020). Nonetheless, this coding scheme captures major patterns of dietary variation without the need to pigeonhole taxa into arbitrary specialist groupings while, critically, providing a means for estimating dietary item importance in taxa of unknown ecology, including fossils. This particular flexibility is paramount in making informed macroevolutionary and paleoecological inferences (Finarelli and Flynn, 2006; Slater et al., 2012).

### Predicting Multivariate Diet from Multivariate Morphology: Challenges and Future Potential

Predicting dietary item importance rankings from morphological data presents novel challenges, but the flexibility of Bayesian multilevel modeling suggests that these methods hold much promise for future work in functional ecology. Past efforts to link morphology and diet have relied on multivariate classification methods, such as discriminant function analysis, to identify linear combinations of traits that maximally distinguish among groups and to classify species of unknown ecology (Hopkins et al., 2022; Boyer, 2008; Friscia et al., 2007; Sacco and Van Valkenburgh, 2004). However, because discriminant functions classify unknowns into one of the sets of grouping variables present in the training set, it is not possible to identify novel ecologies among the set of data-deficient taxa, even though it is reasonable to expect that some extinct taxa may have belonged to dietary niches that are unoccupied by the Recent fauna. Some of the predicted dietary item importance scores we obtain for fossil carnivorans, such as the predicted high importance of large mammals in the diets of sabertoothed cats and nimravids are entirely consistent with results that might have been obtained from a traditional discriminant function analysis using dietary categories (e.g., Van Valkenburgh, 2007, 1988), but other results yield nuanced insights into dietary paleoecology. For example, although relatively generalized diets are inferred for the three small-to-medium sized taxa *Daphoenus*, *“Miacis” latidens*, and *Promartes lepidus*, our approach allows us to identify a more even importance over the suite of food items in the *Daphoenus*, a greater emphasis on hard invertebrates in “*M*.” *latidens*, and a greater importance of small mammals for *P. lepidus*. It is notable that such inference is not possible using standard statistical toolkits and categorical dietary data, and suggests the potential for further clarification of dietary ecology in extinct taxa.

The cave bear presents an altogether different outcome and emphasizes that broader comparative datasets for extant taxa may be necessary to fully leverage our approach. The combination of low *δ*^15^N and *δ*^13^C values from bones and teeth across the species’ range strongly suggest an exclusively herbivorous niche in which forbs and grasses dominated the diet (Bocherens, 2019; Naito et al., 2020). The occlusal topography of mammalian molars exhibits a strong signal of herbivory across diverse clades of mammals (Evans et al., 2007), and we should in principle be able to detect such a diet here (see also Pérez-Ramos et al., 2020). Our results partially match this expectation, with plants predicted to be the most important component of cave bear diet, but with a variety of vertebrate prey predicted to be at least occasionally consumed (Fig. 4B). However, although many carnivorans use some plant materials in their diets, few in our dataset place high importance (rank = 4) on them and this limits the accuracy of our quantitative predictions for this food item (Table 4). For example, among extant taxa, we predict relatively similar weighted importance scores for plants for the bamboo specialist pandas *Ailurus* and *Ailuropoda* and the arctic carnivore polar bear which, by virtue of its ursine ancestry, possesses relatively large blunt molars in comparison with other carnivoran lineages (Sacco and Van Valkenburgh, 2004). The effects of this prediction error may not be restricted to the prediction of individual food item importance scores, but may also be propagated to our nearest neighbor comparisons, where the cave bear is recovered intermediate to the polar bear and giant panda. There has been considerable debate among isotope paleoecologists regarding whether animal protein could have been a regionally important component of cave bear diet (Hilderbrand et al. 1996, Richards et al. 2008, Robu et al. 2013; but see Bocherens 2019, Naito et al. 2020). It is possible that our results lend support to the idea that this taxon occupied a novel portion of carnivoran dietary space in which plants dominated the diet but animal prey could have been a regionally or seasonally important fallback food (Porter et al., 2022). Our models will benefit from an expanded sampling of mammals beyond Carnivora to increase the representation of species that place high importance on underrepresented food items such as plants which should, in turn, improve the accuracy and precision of our dietary item importance estimates and aid in comparisons of the ecological niches of extant and extinct species.

Multinomial Bayesian regressions that allow for the joint estimation and incorporation of phylogenetic signal in residual error have been recently employed for dietary prediction in fossil primates (Fulwood et al., 2021), and we have extended this approach not only by treating importance ranks as ordered variables, but also by including body size as a covariate. Body size alone is a poor predictor of mammalian dietary categories in comparison to functional trait metrics (Grossnickle, 2020), but our results suggest that body size may still modulate the form-function relationship between dental morphology and diet (Figure 3). Such a claim is not without precedent. Fulwood et al. (2021) did not include size as a covariate in their multinomial models but did note that, among primates in general, the teeth of folivores and insectivores resemble one another so closely that body mass must typically be used to discriminate between them (Kay, 1975). Similar effects are apparent in our carnivoran data. For example, lower grinding area (RLGA) effectively predicts the relative importance of large mammals, small mammals, and birds, which are food items with similar mechanical properties. However, large mammals appear to be a less important component of small and medium-bodied carnivoran diets, regardless of RLGA value (Figure 3D), while RLGA is a poor predictor of the importance of birds and small mammals in the diets of large-bodied carnivorans (Figure 3A, E), consistent with the idea that energetic constraints enforce a strong effect on prey size in mammals (Carbone et al., 1999, 2007). More strikingly, high values of DNE for m1, suggestive of a tooth with multiple tall cusps, predict a high importance of soft invertebrates in the diet of small-bodied carnivorans, while the exact opposite relationship is recovered for large-bodied taxa, driven largely by the low profile but more complex (high OPCr) m1 of the sea otter, raccoon, and the bears (Figure 3F). If importance rankings are to provide a fruitful avenue for future investigations of ecological diversity and evolution, then the flexible framework provided by Bayesian multilevel modeling will play a critical role in untangling phylogenetic and allometric relationships between form and function.

Using predictive ordinal modeling to estimate dietary rankings from phenotypic data can help reconstruct dietary partitioning within a community, determine community dietary breadth, and enable explicit tests of ecological redundancies in a probabilistic manner. For example, a persistent question in evolutionary ecology is the role of limiting similarity in community structure (MacArthur and Levins, 1967). Briefly, limiting similarity postulates that two individuals that occupy an identical niche space should not co-occur in space and time. Support of limiting similarity has been equivocal, leading to the emergence of null hypotheses that place no emphasis on species’ adaptations or ecological niche in the formation of communities (Hubbell, 2001; MacArthur and Wilson, 1963). Partitioning of food types is a fundamental aspect of community assembly and maintenance (Grant, 1986; MacArthur and Levins, 1967), and the ranking method presented here holds promise for quantifying diet in both poorly studied extant communities and paleo communities. Generating food consumption probabilities for species within a community naturally segues to comparisons of dietary structure across communities. Examples include comparing dietary structure between communities that occupy similar habitats but vary in species richness or taxonomic structure, or identifying changes in community dietary structure that occur with faunal turnover across space or time. For example, the dietary similarity that we identify between extant hemigaline viverrids of Southeast Asia and procyonids of the Americas suggests incumbency of the latter clade as a potential explanation for why viverrids failed to colonize the New World despite repeated opportunity and an otherwise geographically unconstrained distribution (Hunt, 1996). Baskin (2003) noted that procyonids only became the dominant hypocarnivorous carnivorans in North America after the extinction of phlaocyonine and cynarctine canids (subfamily Borophagine) in the early Hemingfordian and Clarendonian land mammal stages (Early through Middle Miocene), respectively. We can hypothesize that dietary item importance prediction for these canids might yield further similarities to hemigaline viverrids and procyonids, providing additional evidence for the long-term exclusion of feliform hypocarnivores from North America. The approaches outlined in this paper provide a straight-forward way of testing this hypothesis.

The relationship between carnivoran diet and molar structure is well-documented, with numerous examples of how variation in this simple, functional toolkit influences the processing of various food items (Crusafont-Pairó and Truyols-Santonja, 1956; Smits and Evans, 2012; Van Valkenburgh, 2007; Friscia et al., 2007), and serves as a good starting point for building and testing ordinal models of dietary rankings. However, we believe our methodology can be adapted to virtually any system that has sufficient dietary composition data available. One well-studied example is the avian beak, a tool used for both food acquisition and processing (Grant, 1986; Pigot et al., 2020). In this case, different metrics of beak shape, such as length, width, curvature, or keratin thickness, along with other phenotypic traits such as mass or tarsus length, may vary in their ability to predict the quantity of different food types obtained and consumed. Moreover, the method of ordered ranking need not be restricted to diet composition; with importance rankings of locomotor modes, such as swimming, climbing, digging, or flying, models could be constructed to evaluate which phenotypic traits are associated with different foraging strata or microhabitat use. Testing the effect of phenotypic traits on ranked features of diet, behavior, locomotion, microhabitat, or other commonly discretized environmental or biological variables, rather than forcing data into discrete categories, holds great potential for identifying the traits that best predict life history, interactions between multiple traits that are otherwise unobserved, or the presence of correlated traits that may generate noise in categorical analyses.

## Supporting information

Supplementary Methods and Results

## Acknowledgments

We are grateful to Adam Ferguson and Bill Simpson for providing access to specimens. Associate Editor Scott Steppan, Ethan Fullwood, and one anonymous reviewer, as well as David Jablonski, Chris Law, Z. Jack Tseng, provided helpful comments on previous drafts of this manuscript. JAN was funded by an NSF Postdoctoral Fellowship (DBI 2010756).

## Statement of Authorship

All authors contributed equally to conceptualization of the study, and to writing and editing of the manuscript. A.L.W, and G.J.S. generated 3D data. A.L.W. performed dental topographic analyses and coded dietary item importance scores. G.J.S. performed polychoric PCA and nearest neighbor analyses. J.A.N. performed the cluster analyses and Bayesian Ordinal Modeling.

## Data and Code Accessibility Statement

Scripts and datafiles necessary to perform all analyses are available on the Data Dryad digital repository (https://doi.org/10.5061/dryad.pc866t1rg). All tooth surface scan models are available on the Morphosource digital repository (Project ID: 000501405)

