## Supplementary Methods and Results for "Bayesian prediction of multivariate ecology from phenotypic data yields new insights into the diets of extant and extinct taxa"

### Additional Methods

#### *Ecomorphological Data Collection*

We obtained Relative Lower Grinding Area (RLGA, the square root of the combined area of the talonid of the first molar and the entire second molar, relative to the length of the m1 trigonid: Van Valkenburgh, 1988) from Slater and Friscia (2019) and Friscia et al. (2007). This metric has been previously established as a key predictor of categorical dietary strategies in carnivorans (Friscia et al., 2007; Sacco and Van Valkenburgh, 2004; Slater, 2015; Slater and Friscia, 2019; Van Valkenburgh, 1988, 1991) and is therefore used here.

We also generated novel data on the topographic complexity and shape of the first and lower second molars from 3D surface models generated using structured light surface scanners. A subset of specimens was directly scanned with a Capture Mini scanner (3D Systems, North Carolina) at 80 $\mu$m resolution or Comet L3D (Carl Zeiss, Germany) scanner at 18 – 30 $\mu$m point spacing. For the remainder, molds of the lower dentition were made using light and medium body polyvinyl siloxane (Derby Dental Laboratory, Louisville, KY) which were then cast in dental stone and scanned using the CometL3D scanner. 3D models were edited in Geomagic Design X (3D systems, North Carolina). First, we extracted individual lower first (m1) and second (m2) molars from the tooth row by cropping along the enamel-cementum junction. For each molar, we oriented the tooth so that the occlusal plane was facing the positive z-direction and the lingual side was facing the positive x-axis. Right molars were mirrored along the y axis so all molars were left-hand specimens. Each tooth was cleaned to remove artifacts of the scanning process, subjected to an iterative smoothing process, and decimated to 10,000 polygons.

For each molar, we collected three standard measures of dental topography. Relief index (RFI) is the ratio of the 2D planar surface area at the occlusal surface to the 3D surface area above the occlusal surface and is a measure of topographic relief. Teeth with high RFI values tend to belong to species with more insectivorous or folivorous diets, while taxa with low RFI values tend to be frugivorous (Boyer, 2008). Orientation Patch Count rotated (OPCr) (Evans et al., 2007; Wilson et al., 2012) measures the complexity of the tooth by counting the number of contiguous patches on the tooth surface that share a common orientation, and is calculated by grouping points on the surface into patches based on their orientation. Because the orientation of each point is sensitive to the alignment of the tooth model to the global coordinate system, counts are calculated over several small rotations of the model and averaged (Wilson et al., 2012). Species with high OPCr / dental complexity tend to be more herbivorous, while species with low OPCr / dental complexity tend to be more carnivorous (Evans et al., 2007). Dirichlet Normal Energy (DNE) measures the average curvature of a surface by calculating its "bending energy" (Bunn et al., 2011) and captures overall tooth sharpness. High DNE values correlate with insectivory, with intermediate values correlating with folivory and omnivory low values correlated with frugivory.

OPCr and RFI were measured using functions from the R package `molar` (Pampush et al., 2016). We used a robustly implemented algorithm for DNE (ariaDNE) that is less sensitive to artifacts due to 3D modeling, such as smoothing, than earlier DNE algorithms, and which is implemented via `MATLAB` scripts provided by Shan et al. (2019). For carnivoran species without an m2, we assigned a value of zero for all dental topographic measurements.

### Supplemental Figures

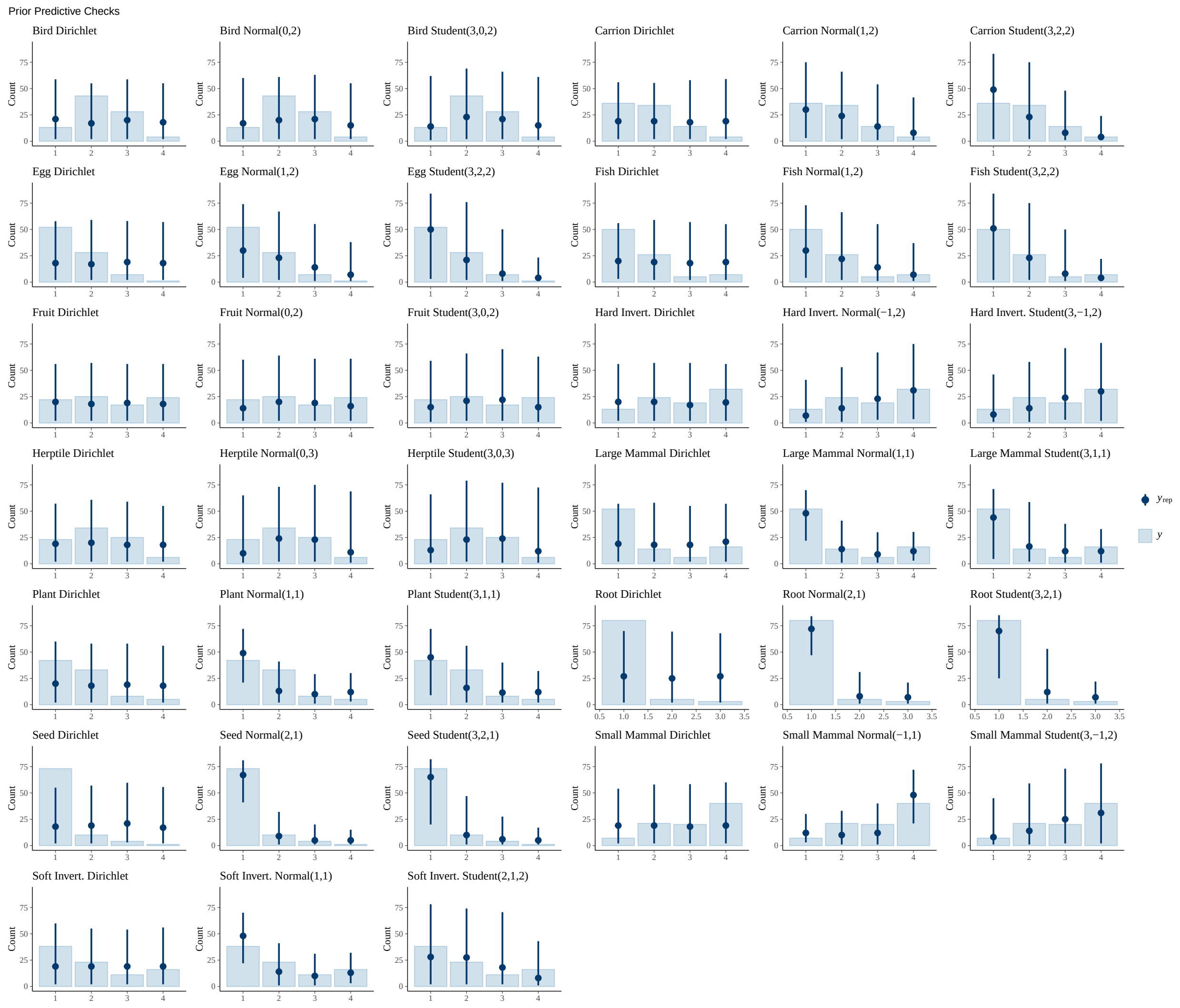


Figure S1: Results from prior predictive simulations of Normal, Student-$T$, and Dirichlet distributions for each food item. The box plots represent the empirical count of each ranking for each food item (n=89 total). The point intervals represent the 89% probability intervals of the prior distributions. The Normal and Student-$T$ distributions that best match the empirical counts of each food item rank are shown. We each of these prior distributions in our prior-predictive simulations for each tooth metric for each food item, then compared the results of these models using Leave One Out Cross Validation.

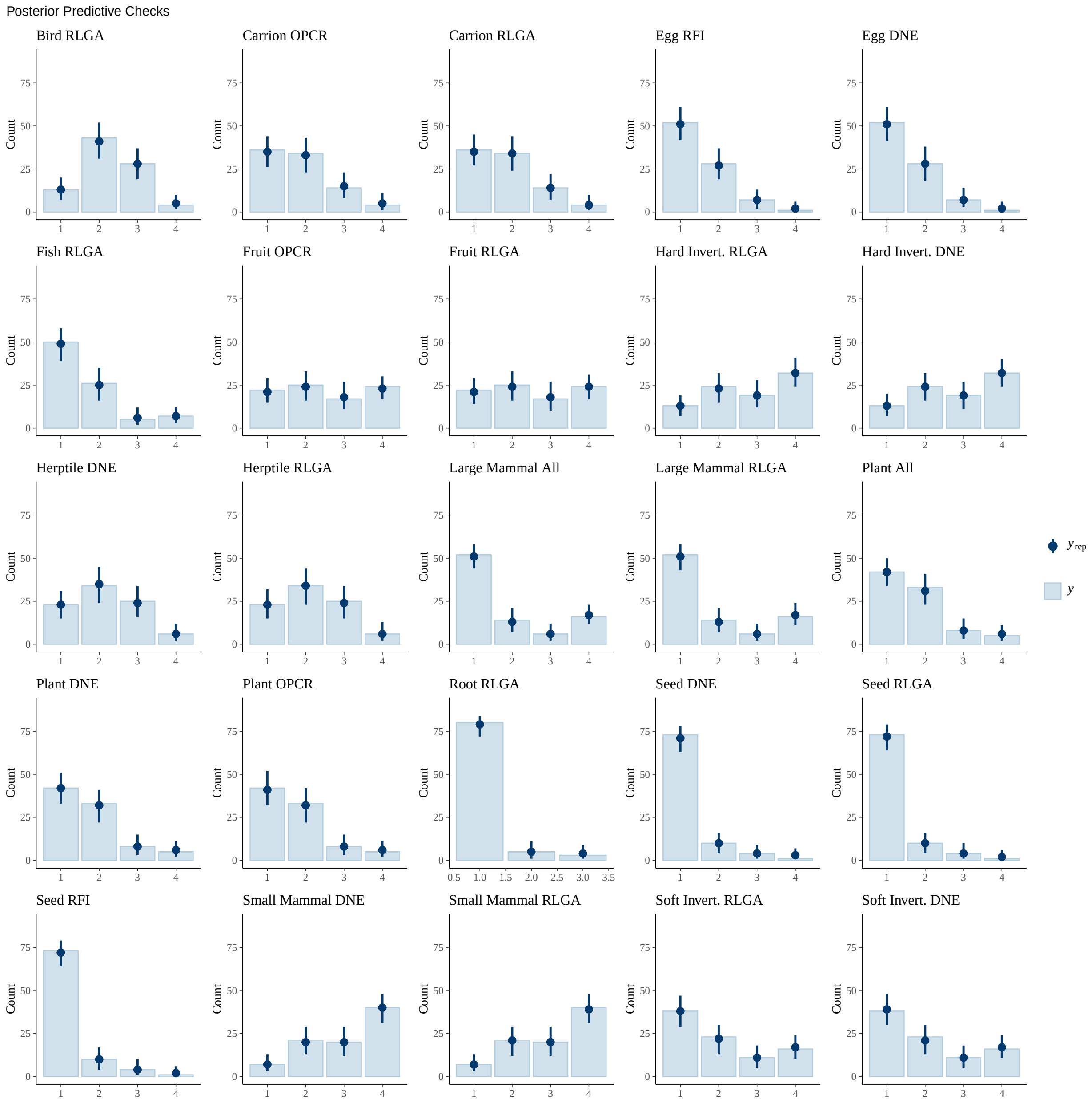


Figure S2: Results from posterior predictive checks from each of the models with a model weight >0. The box plots represent the empirical count of each ranking for each food item (n=89 total). The point intervals represent the 89% probability intervals of the prior distributions. The mean values of the point intervals (dark blue) match the empirical count of ranks (light blue box plot) for each food item, with limited uncertainty, indicating a good model fit to the data.

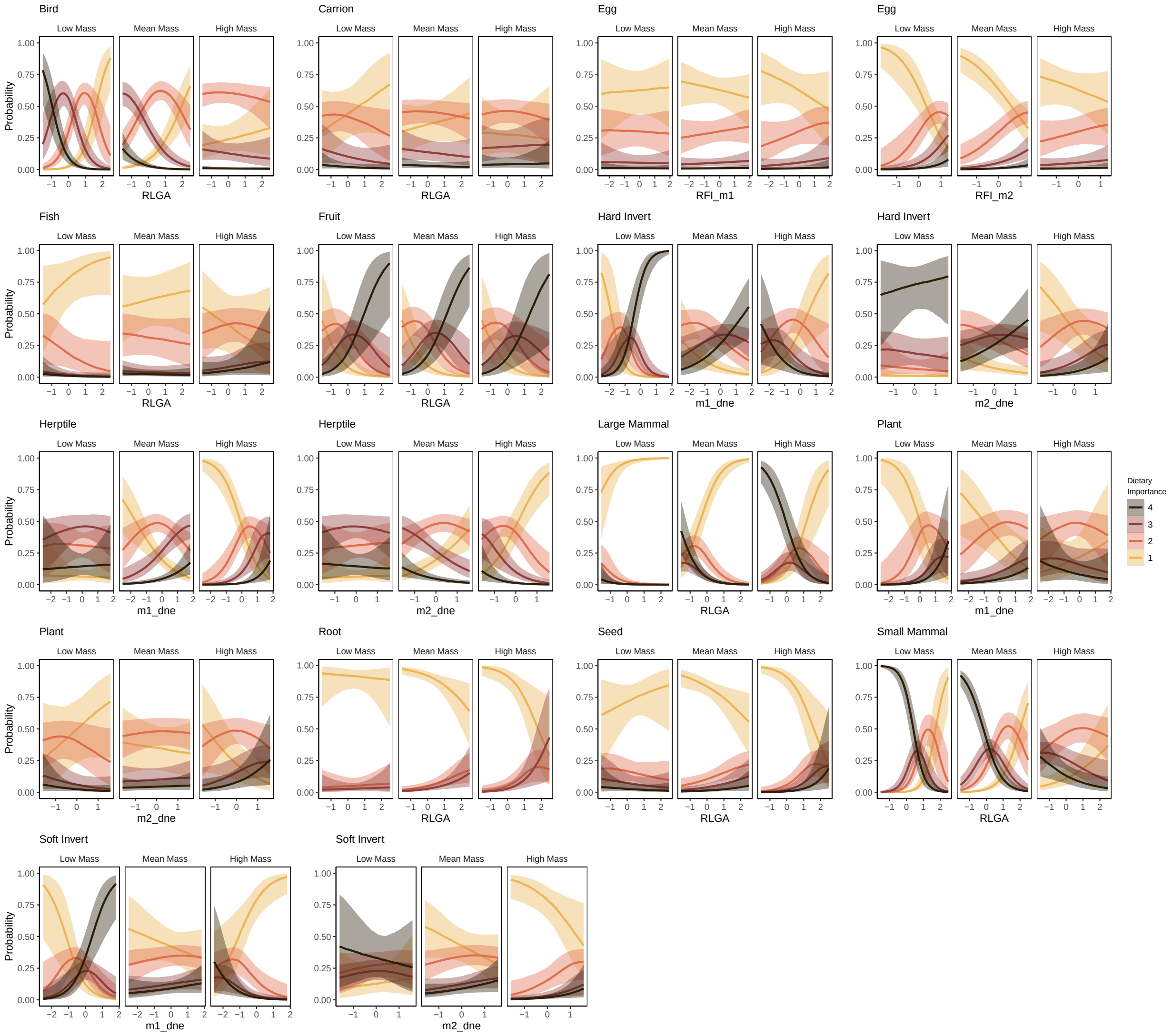


Figure S3: Results from the model for each dietary item with the highest Leave-One-Out cross validation weight. Each triptych plot shows the effect of the dental measurement on the probability of consuming that dietary item at low (-1.5 *sd*), mean, and high (+1.5 *sd*) body mass. The number of plots per dietary item will vary with the dental metric shown. For example, Bird (**S1A**) has only one plot (RLGA is a single value) while Carrion has two (the OPCr of m1 (**S1B**) and m2 (**S1C**)) and Hard Invertebrate has 7 (**S1 I-N**) as 'all metrics' had the highest LOO weight.

### Supplemental Tables

Table S1: Dietary Importance ranks as coded for each extant species across the 13 food items. FMNH is the Field Museum of Natural History Mammals catalog number.

| Species | FMNH | Sources | large mammal | small mammal | bird | herptile | fish | egg | carrion | hard invert | soft invert | seed | fruit | root | plant |
|---|---|---|---|---|---|---|---|---|---|---|---|---|---|---|---|
| *Acinonyx jubatus* | 57826 | Sunquist and Sunquist (2002), Krausman and Morales (2005), Skinner and Chimimba (2005), Craig et al. (2017) | 4 | 2 | 2 | 1 | 1 | 1 | 1 | 1 | 1 | 1 | 1 | 1 | 1 |
| *Ailuropoda melanoleuca* | 31128 | Chorn and Hoffmann (1978); Schaller et al. (1985); Wang et al. (2012) | 1 | 1 | 1 | 1 | 1 | 1 | 2 | 1 | 1 | 1 | 1 | 1 | 4 |
| Ailurus fulgens | 36751 | Choudhury (2001); Glatston et al. (2015); Johnson et al. (1988); Panthi et al. (2012); Pradhan et al. (2001); Roberts and Gittleman (1984) | 1 | 1 | 1 | 1 | 1 | 1 | 1 | 1 | 1 | 1 | 2 | 1 | 4 |
| *Arctictis binturong* | 44726 | Allam et al. (2022); Nakabayashi and Ahmad (2018); Nakabayashi et al. (2017) | 1 | 1 | 1 | 1 | 1 | 1 | 1 | 2 | 1 | 1 | 4 | 1 | 1 |
| *Arctogalidia trivirgata* | 8373 | | NA | NA | NA | NA | NA | NA | NA | NA | NA | NA | NA | NA | NA |
| *Arctonyx collaris* | 39503 | Zhou et al. (2015) | 1 | 2 | 2 | 2 | 1 | 1 | 1 | 4 | 4 | 2 | 4 | 2 | 2 |

| Species | FMNH | Sources | large mammal | small mammal | bird | herptile | fish | egg | carrion | hard invert | soft invert | seed | fruit | root | plant |
|---|---|---|---|---|---|---|---|---|---|---|---|---|---|---|---|
| *Atilax paludinosus* | 27030 | Baker (1992); Do Linh San et al. (2020); Ray (1997) | 2 | 4 | 2 | 4 | 3 | 2 | 2 | 4 | 3 | 2 | 2 | 1 | 1 |
| *Bassaricyon alleni* | 86908 | | NA | NA | NA | NA | NA | NA | NA | NA | NA | NA | NA | NA | NA |
| *Bassariscus astutus* | 51939 | Alexander et al. (1994); Poglayen-Neuwall and Toweill (1988); Rodríguez-Estrella et al. (2000) | 1 | 4 | 2 | 3 | 1 | 1 | 2 | 4 | 4 | 2 | 4 | 1 | 3 |
| *Bdeogale jacksoni* | 85969 | De Luca and Rovero (2006),Kingdon (1977) | 1 | 4 | 2 | 2 | 1 | 2 | 2 | 4 | 2 | 1 | 1 | 1 | 1 |
| *Canis adustus* | 38184 | Sillero-Zubiri et al. (2004), Skinner and Chimimba (2005) | 2 | 4 | 2 | 2 | 1 | 2 | 3 | 4 | 3 | 1 | 4 | 1 | 2 |
| *Canis aureus* | 103909 | Sillero-Zubiri et al. (2004) ,Moehlman and Hayssen (2018) , Jaeger et al. (2007), Markov and Lanszki (2012) | 3 | 4 | 3 | 2 | 2 | 1 | 4 | 2 | 1 | 2 | 3 | 1 | 3 |
| *Canis latrans* | 89416 | Gese et al. (1988); Jensen et al. (2022); Lukasik and Alexander (2011); Sillero-Zubiri et al. (2004) | 3 | 4 | 2 | 2 | 2 | 1 | 3 | 2 | 2 | 1 | 3 | 1 | 3 |

| Species | FMNH | Sources | large mammal | small mammal | bird | herptile | fish | egg | carrion | hard invert | soft invert | seed | fruit | root | plant |
|---|---|---|---|---|---|---|---|---|---|---|---|---|---|---|---|
| *Canis lupus* | 153801 | Lanszki et al. (2012); Sillero-Zubiri et al. (2004); Stahler et al. (2006) | 4 | 3 | 2 | 1 | 2 | 1 | 2 | 1 | 1 | 1 | 3 | 1 | 2 |
| *Canis mesomelas* | 38183 | Walton and Joly (2003), Sillero-Zubiri et al. (2004), Skinner and Chimimba (2005) | 3 | 3 | 3 | 2 | 1 | 1 | 3 | 3 | 3 | 1 | 3 | 1 | 3 |
| *Canis simensis* | 32940 | Sillero-Zubiri et al. (2004), Sillero-Zubiri and Gottelli (1994) | 2 | 4 | 2 | 1 | 1 | 2 | 1 | 1 | 1 | 1 | 1 | 1 | 2 |
| *Caracal caracal* | 105607 | Sunquist and Sunquist (2002), Skinner and Chimimba (2005), Braczkowski et al. (2012), Palmer | 3 | 4 | 4 | 2 | 2 | 1 | 2 | 2 | 1 | 1 | 2 | 1 | 2 |
| *Catopuma badia* | 8378 | | NA | NA | NA | NA | NA | NA | NA | NA | NA | NA | NA | NA | NA |
| *Catopuma temminckii* | 75826 | Kawanishi and Sunquist (2008) Kamler et al. (2020) | 4 | 4 | 3 | 2 | 1 | 1 | 1 | 1 | 1 | 1 | 1 | 1 | 1 |
| *Cerdocyon thous* | 94336 | Lucherini (2015), Berta (1982), Pedó et al. (2006) | 1 | 3 | 3 | 3 | 2 | 2 | 3 | 3 | 1 | 1 | 3 | 1 | 2 |
| *Chrotogale owstoni* | 32556 | | NA | NA | NA | NA | NA | NA | NA | NA | NA | NA | NA | NA | NA |

| Species | FMNH | Sources | large mammal | small mammal | bird | herptile | fish | egg | carrion | hard invert | soft invert | seed | fruit | root | plant |
|---|---|---|---|---|---|---|---|---|---|---|---|---|---|---|---|
| *Chrysocyon brachyurus* | 54406 | Dietz (1985), Machado (2020),Rodden et al. (2004), Santos et al. (2003), Motta-Junior et al. (1996) | 2 | 4 | 3 | 2 | 1 | 2 | 2 | 3 | 1 | 1 | 4 | 1 | 2 |
| *Civettictis civetta* | 155921 | Amiard et al. (2015); Ray (1995); Skinner and Chimimba (2005) | 1 | 3 | 3 | 3 | 2 | 3 | 2 | 4 | 4 | 1 | 4 | 1 | 2 |
| *Conepatus mesoleucus* | 15955 | Helgen (2016*b*), Meaney et al. (2006) | 1 | 2 | 2 | 2 | 1 | 2 | 2 | 4 | 4 | 1 | 3 | 1 | 1 |
| *Crocuta crocuta* | 104022 | Skinner and Chimimba (2005), Di Silvestre et al. (2000), Hayward et al. (2006) | 4 | 2 | 2 | 1 | 2 | 1 | 3 | 2 | 2 | 1 | 2 | 1 | 2 |
| *Crossarchus obscurus* | 43740 | Goldman (1987), Olson (2001) | 1 | 2 | 2 | 2 | 1 | 3 | 1 | 4 | 4 | 1 | 2 | 1 | 1 |
| *Cryptoprocta ferox* | 5655 | Hawkins (2016), Hawkins and Racey (2008), Dollar et al. (2007), Behrens and Barnes (2016) | 2 | 4 | 3 | 3 | 1 | 1 | 1 | 2 | 1 | 1 | 1 | 1 | 1 |
| *Cuon alpinus* | 33500 | Sillero-Zubiri et al. (2004) Cohen (1978) | 4 | 3 | 2 | 2 | 1 | 1 | 2 | 2 | 1 | 1 | 2 | 1 | 2 |

| Species | FMNH | Sources | large mammal | small mammal | bird | herptile | fish | egg | carrion | hard invert | soft invert | seed | fruit | root | plant |
|---|---|---|---|---|---|---|---|---|---|---|---|---|---|---|---|
| *Cynictis penicillata* | 38376 | Herzig-Straschil (1977); Skinner and Chimimba (2005); Taylor and Meester (1993) | 1 | 3 | 3 | 3 | 1 | 2 | 2 | 4 | 4 | 2 | 2 | 1 | 2 |
| *Cynogale bennettii* | 89446 | | NA | NA | NA | NA | NA | NA | NA | NA | NA | NA | NA | NA | NA |
| *Eira barbara* | 21379 | IUCN redlist, Presley (2000), Galef Jr et al. (1976) | 2 | 4 | 2 | 2 | 1 | 1 | 2 | 3 | 3 | 1 | 4 | 1 | 1 |
| *Enhydra lutris* | 78761 | Estes (1980) | 1 | 1 | 1 | 1 | 4 | 1 | 1 | 4 | 4 | 1 | 1 | 1 | 1 |
| *Felis silvestris* | 31127 | Sunquist and Sunquist (2002), Apostolico et al. (2016) | 1 | 4 | 2 | 2 | 2 | 1 | 1 | 2 | 1 | 1 | 2 | 1 | 2 |
| *Fossa fossana* | 15664 | Goodman et al. (2003) | 1 | 3 | 1 | 3 | 1 | 1 | 2 | 4 | 1 | 1 | 1 | 1 | 1 |
| *Galerella sanguinea* | 17810 | Skinner and Chimimba (2005), Graw and Manser (2017) | 1 | 3 | 2 | 3 | 1 | 2 | 1 | 4 | 3 | 1 | 2 | 1 | 1 |
| *Galictis cuja* | 22426 | Yensen and Tarifa (2003) , Diuk-Wasser and Cassini (1998) | 1 | 4 | 3 | 3 | 2 | 1 | 1 | 2 | 2 | 1 | 2 | 1 | 1 |
| *Galidia elegans* | 170876 | | NA | NA | NA | NA | NA | NA | NA | NA | NA | NA | NA | NA | NA |
| *Galidictis fasciata* | 156653 | | NA | NA | NA | NA | NA | NA | NA | NA | NA | NA | NA | NA | NA |

| Species | FMNH | Sources | large mammal | small mammal | bird | herptile | fish | egg | carrion | hard invert | soft invert | seed | fruit | root | plant |
|---|---|---|---|---|---|---|---|---|---|---|---|---|---|---|---|
| *Gulo gulo* | 133755 | Harris and Ogan (1997); Koskela et al. (2013); Pasitschniak-Arts and Larivière (1995); Scrafford and Boyce (2018) | 3 | 4 | 2 | 1 | 2 | 2 | 4 | 1 | 2 | 1 | 2 | 1 | 1 |
| *Helarctos malayanus* | 54201 | Scotson et al. (2017), Fitzgerald and Krausman (2002), Sethy and Chauhan (2018) | 1 | 2 | 1 | 1 | 2 | 1 | 1 | 4 | 4 | 1 | 4 | 2 | 2 |
| *Helogale parvula* | 34201 | Skinner and Chimimba (2005) | 1 | 2 | 1 | 2 | 1 | 2 | 1 | 4 | 4 | 1 | 2 | 1 | 1 |
| *Hemigalus derbyanus* | 68717 | | NA | NA | NA | NA | NA | NA | NA | NA | NA | NA | NA | NA | NA |
| *Herpestes brachyurus* | 88321 | | NA | NA | NA | NA | NA | NA | NA | NA | NA | NA | NA | NA | NA |
| *Hyaena hyaena* | 57973 | Rieger (1981), Kruuk (1976) | 2 | 2 | 3 | 3 | 1 | 1 | 4 | 3 | 2 | 1 | 3 | 1 | 1 |
| *Hydrictis maculicollis* | 125385 | Larivière (2002*a*); Lejeune (1990); Ponsonby (2018); Skinner and Chimimba (2005) | 1 | 2 | 2 | 3 | 4 | 1 | 1 | 4 | 3 | 1 | 1 | 1 | 2 |
| *Ichneumia albicauda* | 26039 | Taylor (1972) | 1 | 2 | 1 | 3 | 1 | 1 | 2 | 4 | 4 | 1 | 2 | 1 | 1 |

| Species | FMNH | Sources | large mammal | small mammal | bird | herptile | fish | egg | carrion | hard invert | soft invert | seed | fruit | root | plant |
|---|---|---|---|---|---|---|---|---|---|---|---|---|---|---|---|
| *Leopardus wiedii* | 52437 | Sunquist and Sunquist (2002) , De Oliveira (1998), Rocha-Mendes et al. (2010) | 1 | 4 | 4 | 3 | 1 | 1 | 1 | 2 | 1 | 1 | 2 | 1 | 1 |
| *Leptailurus serval* | 38192 | Sunquist and Sunquist (2002), | 2 | 4 | 3 | 3 | 1 | 1 | 1 | 3 | 1 | 1 | 1 | 1 | 2 |
| *Lontra canadensis* | 81490 | Larivière and Walton (1998), Buzzell et al. (2014), Knudsen and Hale (1968), Reid et al. (1994), Day et al. (2015), Roberts et al. (2008) | 1 | 2 | 2 | 3 | 4 | 1 | 1 | 4 | 2 | 1 | 2 | 1 | 2 |
| *Lutra lutra* | 39181 | Hung and Law (2016) 2016 | 1 | 3 | 3 | 4 | 4 | 1 | 1 | 4 | 1 | 1 | 2 | 1 | 1 |
| *Lutrogale perspicillata* | 63799 | Ten Hwang and Larivière (2005) | 1 | 3 | 2 | 3 | 4 | 1 | 1 | 3 | 2 | 1 | 1 | 1 | 1 |
| *Lycalopex culpaeus* | 23828 | Sillero-Zubiri et al. (2004), Walker et al. (2007), Guntiñas et al. (2021) | 4 | 4 | 2 | 2 | 1 | 2 | 2 | 2 | 1 | 1 | 3 | 1 | 1 |
| *Lycalopex gymnocercus* | 23819 | Sillero-Zubiri et al. (2004), Lucherini and Luengos Vidal (2008) | 1 | 4 | 3 | 2 | 2 | 1 | 3 | 3 | 3 | 1 | 3 | 1 | 1 |
| *Lycalopex sechurae* | 80967 | Asa and Wallace (1990), Cossíos (2010) | 1 | 3 | 2 | 2 | 1 | 3 | 3 | 3 | 1 | 3 | 4 | 1 | 1 |

| Species | FMNH | Sources | large mammal | small mammal | bird | herptile | fish | egg | carrion | hard invert | soft invert | seed | fruit | root | plant |
|---|---|---|---|---|---|---|---|---|---|---|---|---|---|---|---|
| *Lycalopex vetulus* | 20747 | Dalponte (2009), Sillero-Zubiri et al. (2004) | 1 | 3 | 2 | 2 | 1 | 1 | 1 | 4 | 1 | 1 | 4 | 1 | 1 |
| *Lycaon pictus* | 24326 | Sillero-Zubiri et al. (2004), Skinner and Chimimba (2005) | 4 | 3 | 1 | 1 | 1 | 1 | 2 | 1 | 1 | 1 | 1 | 1 | 1 |
| *Lynx canadensis* | 145825 | Sunquist and Sunquist (2002), Lavoie et al. (2019) | 2 | 4 | 2 | 1 | 1 | 1 | 2 | 1 | 1 | 1 | 1 | 1 | 1 |
| *Martes americana* | 13486 | Clark et al. (1987), Harris and Ogan (1997), Zielinski and Duncan (2004), Nagorsen et al. (1989) | 1 | 4 | 3 | 3 | 3 | 3 | 3 | 3 | 3 | 1 | 3 | 1 | 1 |
| *Mellivora capensis* | 85510 | Begg et al. (2003); Kruuk and Mills (1983); Skinner and Chimimba (2005); Vanderhaar and Ten Hwang (2003) | 1 | 4 | 2 | 4 | 1 | 1 | 2 | 4 | 4 | 1 | 2 | 2 | 1 |
| *Melursus ursinus* | 30253 | Dharaiya et al. (2020); Laurie and Seidensticker (1977); Mewada and Dharaiya (2010) | 1 | 2 | 2 | 2 | 1 | 1 | 2 | 4 | 1 | 2 | 4 | 2 | 2 |
| *Mephitis mephitis* | 51659 | Azevedo et al. (2006); Wade-Smith and Verts (1982) | 1 | 3 | 3 | 3 | 1 | 3 | 1 | 4 | 4 | 1 | 3 | 1 | 1 |

| Species | FMNH | Sources | large mammal | small mammal | bird | herptile | fish | egg | carrion | hard invert | soft invert | seed | fruit | root | plant |
|---|---|---|---|---|---|---|---|---|---|---|---|---|---|---|---|
| *Mungotictis decemlineata* | 176129 | Behrens and Barnes (2016), Rasolofoniaina et al. (2019), Woolaver et al. (2006) | 1 | 2 | 2 | 2 | 1 | 2 | 1 | 4 | 4 | 1 | 1 | 1 | 1 |
| *Mydaus javanensis* | 68730 | | NA | NA | NA | NA | NA | NA | NA | NA | NA | NA | NA | NA | NA |
| *Nandinia binotata* | 161266 | Azevedo et al. (2006); Charles-Dominique (1978); Skinner and Chimimba (2005) | 1 | 2 | 2 | 1 | 1 | 2 | 1 | 2 | 2 | 1 | 4 | 1 | 1 |
| *Nasua nasua* | 70735 | Gompper and Decker (1998), Alves-Costa et al. (2004), Ferreira et al. (2013) | 1 | 2 | 2 | 2 | 2 | 2 | 2 | 4 | 4 | 1 | 4 | 1 | 2 |
| *Nasuella olivacea* | 89236 | Balaguera-Reina et al. (2009), Rodríguez-Bolaños et al. (2000) | 1 | 1 | 1 | 3 | 1 | 1 | 2 | 4 | 4 | 1 | 4 | 2 | 2 |
| *Neofelis nebulosa* | 42583 | Sunquist and Sunquist (2002) | 4 | 4 | 2 | 1 | 2 | 1 | 1 | 1 | 1 | 1 | 1 | 1 | 1 |
| *Neogale vison* | 136259 | Larivière (1999) | 1 | 4 | 3 | 4 | 4 | 2 | 2 | 4 | 2 | 1 | 1 | 1 | 1 |
| *Nyctereutes procyonoides* | 33615 | Ward and Wurster-Hill (1990), Sillero-Zubiri et al. (2004) | 1 | 3 | 3 | 3 | 3 | 3 | 3 | 3 | 3 | 3 | 3 | 3 | 3 |

| Species | FMNH | Sources | large mammal | small mammal | bird | herptile | fish | egg | carrion | hard invert | soft invert | seed | fruit | root | plant |
|---|---|---|---|---|---|---|---|---|---|---|---|---|---|---|---|
| *Otocyon megalotis* | 1469 | Clark (2005); Klare et al. (2011); Skinner and Chimimba (2005) | 1 | 2 | 2 | 2 | 1 | 2 | 2 | 4 | 2 | 2 | 4 | 1 | 2 |
| *Paguma larvata* | 88308 | Iwama et al. (2017), Zhou et al. (2008), Torii (1986) | 1 | 4 | 3 | 2 | 2 | 1 | 1 | 4 | 1 | 1 | 4 | 1 | 2 |
| *Panthera leo* | 75608 | Haas et al. (2005), Sunquist and Sunquist (2002), Skinner and Chimimba (2005) | 4 | 2 | 2 | 2 | 2 | 2 | 2 | 2 | 1 | 1 | 2 | 1 | 2 |
| *Panthera onca* | 21392 | Sunquist and Sunquist (2002), Seymour (1989) | 4 | 4 | 2 | 4 | 2 | 2 | 1 | 2 | 1 | 1 | 2 | 1 | 2 |
| *Panthera pardus* | 60615 | Sunquist and Sunquist (2002) , Stein and Hayssen (2013), Skinner and Chimimba (2005) | 4 | 4 | 2 | 2 | 2 | 1 | 1 | 2 | 1 | 1 | 2 | 1 | 2 |
| *Panthera uncia* | 43113 | Sunquist and Sunquist (2002), Hemmer (1972) | 4 | 3 | 3 | 1 | 1 | 1 | 2 | 1 | 1 | 1 | 1 | 1 | 2 |
| *Paradoxurus musanga* | 8382 | | NA | NA | NA | NA | NA | NA | NA | NA | NA | NA | NA | NA | NA |
| *Parahyaena brunnea* | 34584 | Faure et al. (2019); Skinner and Chimimba (2005) | 2 | 3 | 2 | 2 | 1 | 2 | 4 | 2 | 1 | 1 | 3 | 1 | 1 |
| *Pardofelis marmorata* | 60020 | | NA | NA | NA | NA | NA | NA | NA | NA | NA | NA | NA | NA | NA |

| Species | FMNH | Sources | large mammal | small mammal | bird | herptile | fish | egg | carrion | hard invert | soft invert | seed | fruit | root | plant |
|---|---|---|---|---|---|---|---|---|---|---|---|---|---|---|---|
| *Pekania pennanti* | 6341 | Arthur et al. (1989); Golightly et al. (2012); Powell (1981) | 3 | 4 | 2 | 3 | 1 | 2 | 3 | 2 | 2 | 1 | 3 | 1 | 1 |
| *Potos flavus* | 68882 | Kays (1999), Ford and Hoffmann (1988), | 1 | 1 | 1 | 1 | 1 | 1 | 1 | 2 | 2 | 1 | 4 | 1 | 3 |
| *Prionailurus bengalensis* | 31782 | Sunquist and Sunquist (2002) | 2 | 4 | 4 | 4 | 1 | 1 | 2 | 3 | 2 | 1 | 1 | 1 | 2 |
| *Prionodon linsang* | 8371 | | NA | NA | NA | NA | NA | NA | NA | NA | NA | NA | NA | NA | NA |
| *Procyon cancrivorus* | 95528 | Gatti et al. (2006), Quintela et al. (2014), Martinelli and Volpi (2010) | 1 | 3 | 2 | 3 | 2 | 1 | 1 | 3 | 1 | 1 | 4 | 1 | 1 |
| *Procyon lotor* | 90555 | Lotze and Anderson (1979), Dorney (1954) | 1 | 2 | 2 | 2 | 2 | 2 | 2 | 4 | 1 | 4 | 4 | 1 | 2 |
| *Proteles cristatus* | 211365 | Skinner and Chimimba (2005) | 1 | 1 | 1 | 1 | 1 | 1 | 1 | 4 | 2 | 1 | 1 | 1 | 1 |
| *Pteronura brasiliensis* | 70768 | Noonan et al. (2017), Rosas et al. (1999), Cabral et al. (2010), Silva et al. (2014), | 2 | 2 | 1 | 2 | 4 | 1 | 1 | 2 | 1 | 1 | 1 | 1 | 1 |
| *Puma concolor* | 78097 | Sunquist and Sunquist (2002) | 4 | 4 | 2 | 2 | 1 | 1 | 1 | 1 | 1 | 1 | 1 | 1 | 2 |
| *Speothos venaticus* | 60290 | Sillero-Zubiri et al. (2004), de Mello Beisiegel and Zuercher (2005) | 4 | 4 | 3 | 3 | 1 | 1 | 1 | 1 | 1 | 1 | 2 | 1 | 1 |

| Species | FMNH | Sources | large mammal | small mammal | bird | herptile | fish | egg | carrion | hard invert | soft invert | seed | fruit | root | plant |
|---|---|---|---|---|---|---|---|---|---|---|---|---|---|---|---|
| *Spilogale putorius* | 15056 | Kinlaw et al. (1995), Baker and Baker (1975) | 1 | 4 | 3 | 2 | 1 | 2 | 2 | 4 | 4 | 1 | 2 | 1 | 3 |
| *Taxidea taxus* | 63899 | Azevedo et al. (2006); Helgen (2016*a*); Hoodicoff (2003); Long (1973); Sovada et al. (1999); **?** | 1 | 4 | 3 | 2 | 2 | 3 | 2 | 3 | 2 | 2 | 1 | 1 | 2 |
| *Tremarctos ornatus* | 78678 | Velez-Liendo and García-Rangel (2017), VVela-Vargas et al. (2021), García-Rangel (2012) | 2 | 2 | 2 | 1 | 1 | 1 | 2 | 2 | 2 | 1 | 4 | 1 | 4 |
| *Urocyon cinereoargenteus* | 141988 | Sillero-Zubiri et al. (2004), Fritzell and Haroldson (1982), Hockman and Chapman (1983) | 1 | 4 | 3 | 1 | 1 | 1 | 2 | 4 | 2 | 3 | 4 | 1 | 1 |
| *Ursus arctos* | 65738 | Pasitschniak-Arts (1993) | 4 | 2 | 2 | 1 | 3 | 2 | 3 | 2 | 2 | 2 | 4 | 3 | 4 |
| *Ursus maritimus* | 51473 | DeMaster and Stirling (1981), Gormezano and Rockwell (2013) | 4 | 2 | 2 | 1 | 2 | 2 | 3 | 2 | 2 | 1 | 2 | 1 | 2 |
| *Viverra zibetha* | 36009 | Kanchanasakha (2000), Timmins et al. (2016), Myers (2016) | 1 | 3 | 2 | 2 | 2 | 2 | 1 | 2 | 2 | 1 | 4 | 1 | 1 |

| Species | FMNH | Sources | large mammal | small mammal | bird | herptile | fish | egg | carrion | hard invert | soft invert | seed | fruit | root | plant |
|---|---|---|---|---|---|---|---|---|---|---|---|---|---|---|---|
| *Viverricula indica* | 75864 | Chuang and Lee (1997), Wang and Fuller (2003) | 1 | 4 | 2 | 2 | 2 | 1 | 1 | 4 | 4 | 1 | 4 | 1 | 4 |
| *Vormela peregusna* | 48482 | Gorsuch and Larivière (2005) | 1 | 4 | 3 | 3 | 1 | 1 | 1 | 4 | 1 | 1 | 2 | 1 | 2 |
| *Vulpes chama* | 85881 | Skinner and Chimimba (2005), Sillero-Zubiri et al. (2004) | 1 | 4 | 3 | 3 | 1 | 3 | 2 | 3 | 3 | 1 | 2 | 1 | 2 |
| *Vulpes lagopus* | 26882 | Sillero-Zubiri et al. (2004) , Audet et al. (2002), Macpherson (1969), Fay and Stephenson (1989), Møller Nielsen (1991) | 2 | 4 | 4 | 1 | 3 | 4 | 3 | 3 | 3 | 1 | 2 | 1 | 1 |
| *Vulpes macrotis* | 15041 | Sillero-Zubiri et al. (2004), McGrew (1979) | 1 | 4 | 2 | 2 | 1 | 1 | 2 | 2 | 1 | 1 | 2 | 1 | 2 |
| *Vulpes rueppellii* | 106368 | Sillero-Zubiri et al. (2004), Larivière and Seddon (2001) | 1 | 3 | 3 | 3 | 1 | 1 | 1 | 3 | 1 | 1 | 3 | 1 | 3 |
| *Vulpes velox* | 129301 | Sillero-Zubiri et al. (2004), Egoscue (1979) | 1 | 4 | 3 | 3 | 2 | 2 | 3 | 3 | 2 | 2 | 2 | 1 | 2 |
| *Vulpes vulpes* | 67415 | Larivière and Pasitschniak-Arts (1996),Hockman and Chapman (1983) | 1 | 4 | 3 | 1 | 2 | 2 | 2 | 3 | 3 | 3 | 3 | 1 | 1 |

| Species | FMNH | Sources | large mammal | small mammal | bird | herptile | fish | egg | carrion | hard invert | soft invert | seed | fruit | root | plant |
|---|---|---|---|---|---|---|---|---|---|---|---|---|---|---|---|
| *Vulpes zerda* | 89938 | Sillero-Zubiri et al. (2004), Larivière (2002*b*), Brahmi et al. (2012) | 1 | 3 | 3 | 3 | 1 | 2 | 1 | 4 | 2 | 1 | 3 | 3 | 1 |

Table S2: Model-averaged Pareto-$k$ scores for each species for each food item. Pareto-$k$ scores are calculated using Pareto-smoothed-importance-sampling that is integrated into leave-one-out cross validation in the R package LOO. The scores provide an estimate of the importance of each data point to the posterior distribution. When a single sample is removed during loo cv, and the posterior distribution changes very little, then the Pareto-$k$ score for the sample is low. The larger the influence of a single sample on the overall posterior, the higher the Pareto-$k$ score. If many Pareto-$k$ scores are high, this means the model is learning too much from the data, and it is overfit.

| **Species** | **Food Item** | **Pareto-$k$** |
|---|---|---|
| Acinonyx_jubatus | bird | 0.033 |
| Ailuropoda_melanoleuca | bird | 0.055 |
| Ailurus_fulgens | bird | 0.058 |
| Arctictis_binturong | bird | 0.042 |
| Arctonyx_collaris | bird | 0.083 |
| Atilax_paludinosus | bird | 0.033 |
| Bassariscus_astutus | bird | 0.071 |
| Bdeogale_jacksoni | bird | 0.058 |
| Canis_adustus | bird | 0.014 |
| Canis_aureus | bird | 0.022 |
| Canis_latrans | bird | 0.052 |
| Canis_lupus | bird | 0.023 |
| Canis_mesomelas | bird | 0.034 |
| Canis_simensis | bird | 0.030 |
| Caracal_caracal | bird | 0.074 |
| Catopuma_temminckii | bird | 0.026 |
| Cerdocyon_thous | bird | 0.044 |
| Chrysocyon_brachyurus | bird | 0.015 |
| Civettictis_civetta | bird | 0.069 |
| Conepatus_mesoleucus | bird | 0.099 |
| Crocuta_crocuta | bird | 0.070 |
| Cryptoprocta_ferox | bird | 0.089 |
| Cuon_alpinus | bird | 0.043 |
| Cynictis_penicillata | bird | 0.076 |
| Eira_barbara | bird | 0.034 |
| Enhydra_lutris | bird | 0.047 |
| Felis_silvestris | bird | 0.073 |
| Fossa_fossana | bird | 0.067 |
| Galictis_cuja | bird | 0.072 |
| Gulo_gulo | bird | 0.044 |
| Helarctos_malayanus | bird | 0.035 |
| Helogale_parvula | bird | 0.037 |
| Hyaena_hyaena | bird | 0.071 |
| Hydrictis_maculicollis | bird | 0.074 |
| Ichneumia_albicauda | bird | 0.038 |
| Leopardus_wiedii | bird | 0.078 |
| Leptailurus_serval | bird | 0.032 |
| Lontra_canadensis | bird | 0.056 |
| Lutra_lutra | bird | 0.046 |
| Lutrogale_perspicillata | bird | 0.053 |
| Lycalopex_culpaeus | bird | 0.019 |

| **Species** | **Food Item** | **Pareto-*k*** |
|---|---|---|
| Lycalopex_gymnocercus | bird | 0.021 |
| Lycalopex_sechurae | bird | 0.048 |
| Lycalopex_vetulus | bird | 0.019 |
| Lycaon_pictus | bird | 0.049 |
| Lynx_canadensis | bird | 0.062 |
| Martes_americana | bird | 0.083 |
| Mellivora_capensis | bird | 0.053 |
| Melursus_ursinus | bird | 0.035 |
| Mephitis_mephitis | bird | 0.067 |
| Mungotictis_decemlineata | bird | 0.025 |
| Nandinia_binotata | bird | 0.076 |
| Nasua_nasua | bird | 0.059 |
| Nasuella_olivacea | bird | 0.049 |
| Neofelis_nebulosa | bird | 0.083 |
| Neovison_vison | bird | 0.088 |
| Nyctereutes_procyonoides | bird | 0.046 |
| Otocyon_megalotis | bird | 0.049 |
| Paguma_larvata | bird | 0.100 |
| Panthera_leo | bird | 0.040 |
| Panthera_onca | bird | 0.053 |
| Panthera_pardus | bird | 0.034 |
| Parahyaena_brunnea | bird | 0.053 |
| Pekania_pennanti | bird | 0.043 |
| Potos_flavus | bird | 0.063 |
| Prionailurus_bengalensis | bird | 0.040 |
| Procyon_cancrivorus | bird | 0.051 |
| Procyon_lotor | bird | 0.038 |
| Proteles_cristatus | bird | 0.054 |
| Pteronura_brasiliensis | bird | 0.057 |
| Puma_concolor | bird | 0.019 |
| Speothos_venaticus | bird | 0.033 |
| Spilogale_putorius | bird | 0.077 |
| Taxidea_taxus | bird | 0.071 |
| Tremarctos_ornatus | bird | 0.064 |
| Uncia_uncia | bird | 0.041 |
| Urocyon_cinereoargenteus | bird | 0.035 |
| Ursus_arctos | bird | 0.067 |
| Ursus_maritimus | bird | 0.062 |
| Viverra_zibetha | bird | 0.047 |
| Viverricula_indica | bird | 0.057 |
| Vulpes_chama | bird | 0.008 |
| Vulpes_lagopus | bird | 0.064 |
| Vulpes_macrotis | bird | 0.053 |
| Vulpes_rueppellii | bird | 0.003 |
| Vulpes_velox | bird | 0.021 |
| Vulpes_vulpes | bird | 0.002 |
| Vulpes_zerda | bird | 0.016 |
| Acinonyx_jubatus | carrion | 0.051 |

| **Species** | **Food Item** | **Pareto-*k*** |
|---|---|---|
| Ailuropoda_melanoleuca | carrion | 0.087 |
| Ailurus_fulgens | carrion | 0.096 |
| Arctictis_binturong | carrion | 0.077 |
| Arctonyx_collaris | carrion | 0.090 |
| Atilax_paludinosus | carrion | 0.073 |
| Bassariscus_astutus | carrion | 0.081 |
| Bdeogale_jacksoni | carrion | 0.081 |
| Canis_adustus | carrion | 0.065 |
| Canis_aureus | carrion | 0.055 |
| Canis_latrans | carrion | 0.064 |
| Canis_lupus | carrion | 0.066 |
| Canis_mesomelas | carrion | 0.050 |
| Canis_simensis | carrion | 0.040 |
| Caracal_caracal | carrion | 0.087 |
| Catopuma_temminckii | carrion | 0.076 |
| Cerdocyon_thous | carrion | 0.025 |
| Chrysocyon_brachyurus | carrion | 0.068 |
| Civettictis_civetta | carrion | 0.135 |
| Conepatus_mesoleucus | carrion | 0.083 |
| Crocuta_crocuta | carrion | 0.063 |
| Cryptoprocta_ferox | carrion | 0.109 |
| Cuon_alpinus | carrion | 0.066 |
| Cynictis_penicillata | carrion | 0.050 |
| Eira_barbara | carrion | 0.072 |
| Enhydra_lutris | carrion | 0.072 |
| Felis_silvestris | carrion | 0.066 |
| Fossa_fossana | carrion | 0.095 |
| Galictis_cuja | carrion | 0.055 |
| Gulo_gulo | carrion | 0.064 |
| Helarctos_malayanus | carrion | 0.064 |
| Helogale_parvula | carrion | 0.098 |
| Hyaena_hyaena | carrion | 0.051 |
| Hydrictis_maculicollis | carrion | 0.052 |
| Ichneumia_albicauda | carrion | 0.049 |
| Leopardus_wiedii | carrion | 0.074 |
| Leptailurus_serval | carrion | 0.050 |
| Lontra_canadensis | carrion | 0.041 |
| Lutra_lutra | carrion | 0.052 |
| Lutrogale_perspicillata | carrion | 0.050 |
| Lycalopex_culpaeus | carrion | 0.044 |
| Lycalopex_gymnocercus | carrion | 0.062 |
| Lycalopex_sechurae | carrion | 0.031 |
| Lycalopex_vetulus | carrion | 0.056 |
| Lycaon_pictus | carrion | 0.064 |
| Lynx_canadensis | carrion | 0.058 |
| Martes_americana | carrion | 0.044 |
| Mellivora_capensis | carrion | 0.095 |
| Melursus_ursinus | carrion | 0.073 |

| **Species** | **Food Item** | **Pareto-$k$** |
|---|---|---|
| Mephitis_mephitis | carrion | 0.087 |
| Mungotictis_decemlineata | carrion | 0.056 |
| Nandinia_binotata | carrion | 0.114 |
| Nasua_nasua | carrion | 0.070 |
| Nasuella_olivacea | carrion | 0.106 |
| Neofelis_nebulosa | carrion | 0.046 |
| Neovison_vison | carrion | 0.098 |
| Nyctereutes_procyonoides | carrion | 0.081 |
| Otocyon_megalotis | carrion | 0.095 |
| Paguma_larvata | carrion | 0.068 |
| Panthera_leo | carrion | 0.074 |
| Panthera_onca | carrion | 0.064 |
| Panthera_pardus | carrion | 0.049 |
| Parahyaena_brunnea | carrion | 0.068 |
| Pekania_pennanti | carrion | 0.083 |
| Potos_flavus | carrion | 0.075 |
| Prionailurus_bengalensis | carrion | 0.087 |
| Procyon_cancrivorus | carrion | 0.038 |
| Procyon_lotor | carrion | 0.080 |
| Proteles_cristatus | carrion | 0.108 |
| Pteronura_brasiliensis | carrion | 0.048 |
| Puma_concolor | carrion | 0.030 |
| Speothos_venaticus | carrion | 0.075 |
| Spilogale_putorius | carrion | 0.097 |
| Taxidea_taxus | carrion | 0.054 |
| Tremarctos_ornatus | carrion | 0.082 |
| Uncia_uncia | carrion | 0.046 |
| Urocyon_cinereoargenteus | carrion | 0.068 |
| Ursus_arctos | carrion | 0.070 |
| Ursus_maritimus | carrion | 0.067 |
| Viverra_zibetha | carrion | 0.064 |
| Viverricula_indica | carrion | 0.072 |
| Vulpes_chama | carrion | 0.047 |
| Vulpes_lagopus | carrion | 0.037 |
| Vulpes_macrotis | carrion | 0.056 |
| Vulpes_rueppellii | carrion | 0.051 |
| Vulpes_velox | carrion | 0.067 |
| Vulpes_vulpes | carrion | 0.086 |
| Vulpes_zerda | carrion | 0.060 |
| Acinonyx_jubatus | egg | 0.058 |
| Ailuropoda_melanoleuca | egg | 0.073 |
| Ailurus_fulgens | egg | 0.063 |
| Arctictis_binturong | egg | 0.081 |
| Arctonyx_collaris | egg | 0.065 |
| Atilax_paludinosus | egg | 0.061 |
| Bassariscus_astutus | egg | 0.060 |
| Bdeogale_jacksoni | egg | 0.034 |
| Canis_adustus | egg | 0.047 |

| **Species** | **Food Item** | **Pareto-*k*** |
|---|---|---|
| Canis_aureus | egg | 0.009 |
| Canis_latrans | egg | 0.027 |
| Canis_lupus | egg | 0.058 |
| Canis_mesomelas | egg | 0.032 |
| Canis_simensis | egg | 0.074 |
| Caracal_caracal | egg | 0.026 |
| Catopuma_temminckii | egg | 0.025 |
| Cerdocyon_thous | egg | 0.046 |
| Chrysocyon_brachyurus | egg | -0.005 |
| Civettictis_civetta | egg | 0.060 |
| Conepatus_mesoleucus | egg | 0.057 |
| Crocuta_crocuta | egg | 0.051 |
| Cryptoprocta_ferox | egg | 0.048 |
| Cuon_alpinus | egg | 0.036 |
| Cynictis_penicillata | egg | 0.067 |
| Eira_barbara | egg | 0.091 |
| Enhydra_lutris | egg | 0.059 |
| Felis_silvestris | egg | 0.042 |
| Fossa_fossana | egg | 0.067 |
| Galictis_cuja | egg | 0.069 |
| Gulo_gulo | egg | 0.072 |
| Helarctos_malayanus | egg | 0.057 |
| Helogale_parvula | egg | 0.090 |
| Hyaena_hyaena | egg | 0.063 |
| Hydrictis_maculicollis | egg | 0.058 |
| Ichneumia_albicauda | egg | 0.067 |
| Leopardus_wiedii | egg | 0.048 |
| Leptailurus_serval | egg | 0.033 |
| Lontra_canadensis | egg | 0.047 |
| Lutra_lutra | egg | 0.036 |
| Lutrogale_perspicillata | egg | 0.040 |
| Lycalopex_culpaeus | egg | 0.037 |
| Lycalopex_gymnocercus | egg | 0.055 |
| Lycalopex_sechurae | egg | 0.044 |
| Lycalopex_vetulus | egg | 0.044 |
| Lycaon_pictus | egg | 0.045 |
| Lynx_canadensis | egg | 0.017 |
| Martes_americana | egg | 0.050 |
| Mellivora_capensis | egg | 0.050 |
| Melursus_ursinus | egg | 0.045 |
| Mephitis_mephitis | egg | 0.059 |
| Mungotictis_decemlineata | egg | 0.062 |
| Nandinia_binotata | egg | 0.061 |
| Nasua_nasua | egg | 0.081 |
| Nasuella_olivacea | egg | 0.036 |
| Neofelis_nebulosa | egg | 0.046 |
| Neovison_vison | egg | 0.102 |
| Nyctereutes_procyonoides | egg | 0.045 |

| Species | Food Item | Pareto-$k$ |
|---|---|---|
| Otocyon_megalotis | egg | 0.055 |
| Paguma_larvata | egg | 0.089 |
| Panthera_leo | egg | 0.087 |
| Panthera_onca | egg | 0.094 |
| Panthera_pardus | egg | 0.072 |
| Parahyaena_brunnea | egg | 0.087 |
| Pekania_pennanti | egg | 0.053 |
| Potos_flavus | egg | 0.079 |
| Prionailurus_bengalensis | egg | 0.063 |
| Procyon_cancrivorus | egg | 0.038 |
| Procyon_lotor | egg | 0.054 |
| Proteles_cristatus | egg | 0.045 |
| Pteronura_brasiliensis | egg | 0.044 |
| Puma_concolor | egg | 0.042 |
| Speothos_venaticus | egg | 0.054 |
| Spilogale_putorius | egg | 0.044 |
| Taxidea_taxus | egg | 0.091 |
| Tremarctos_ornatus | egg | 0.052 |
| Uncia_uncia | egg | 0.051 |
| Urocyon_cinereoargenteus | egg | 0.071 |
| Ursus_arctos | egg | 0.061 |
| Ursus_maritimus | egg | 0.088 |
| Viverra_zibetha | egg | 0.083 |
| Viverricula_indica | egg | 0.073 |
| Vulpes_chama | egg | 0.033 |
| Vulpes_lagopus | egg | 0.071 |
| Vulpes_macrotis | egg | 0.041 |
| Vulpes_rueppellii | egg | 0.049 |
| Vulpes_velox | egg | 0.055 |
| Vulpes_vulpes | egg | 0.037 |
| Vulpes_zerda | egg | 0.041 |
| Acinonyx_jubatus | fish | 0.066 |
| Ailuropoda_melanoleuca | fish | 0.079 |
| Ailurus_fulgens | fish | 0.083 |
| Arctictis_binturong | fish | 0.099 |
| Arctonyx_collaris | fish | 0.037 |
| Atilax_paludinosus | fish | 0.104 |
| Bassariscus_astutus | fish | 0.078 |
| Bdeogale_jacksoni | fish | 0.046 |
| Canis_adustus | fish | 0.053 |
| Canis_aureus | fish | 0.051 |
| Canis_latrans | fish | 0.081 |
| Canis_lupus | fish | 0.059 |
| Canis_mesomelas | fish | 0.015 |
| Canis_simensis | fish | 0.063 |
| Caracal_caracal | fish | 0.083 |
| Catopuma_temminckii | fish | 0.028 |
| Cerdocyon_thous | fish | 0.082 |

| Species | Food Item | Pareto-$k$ |
|---|---|---|
| Chrysocyon_brachyurus | fish | 0.050 |
| Civettictis_civetta | fish | 0.053 |
| Conepatus_mesoleucus | fish | 0.069 |
| Crocuta_crocuta | fish | 0.152 |
| Cryptoprocta_ferox | fish | 0.041 |
| Cuon_alpinus | fish | 0.070 |
| Cynictis_penicillata | fish | 0.089 |
| Eira_barbara | fish | 0.080 |
| Enhydra_lutris | fish | 0.050 |
| Felis_silvestris | fish | 0.137 |
| Fossa_fossana | fish | 0.103 |
| Galictis_cuja | fish | 0.119 |
| Gulo_gulo | fish | 0.060 |
| Helarctos_malayanus | fish | 0.065 |
| Helogale_parvula | fish | 0.097 |
| Hyaena_hyaena | fish | 0.079 |
| Hydrictis_maculicollis | fish | 0.051 |
| Ichneumia_albicauda | fish | 0.083 |
| Leopardus_wiedii | fish | 0.061 |
| Leptailurus_serval | fish | 0.044 |
| Lontra_canadensis | fish | 0.045 |
| Lutra_lutra | fish | 0.091 |
| Lutrogale_perspicillata | fish | 0.058 |
| Lycalopex_culpaeus | fish | 0.062 |
| Lycalopex_gymnocercus | fish | 0.076 |
| Lycalopex_sechurae | fish | 0.067 |
| Lycalopex_vetulus | fish | 0.019 |
| Lycaon_pictus | fish | 0.050 |
| Lynx_canadensis | fish | 0.028 |
| Martes_americana | fish | 0.060 |
| Mellivora_capensis | fish | 0.075 |
| Melursus_ursinus | fish | 0.057 |
| Mephitis_mephitis | fish | 0.135 |
| Mungotictis_decemlineata | fish | 0.108 |
| Nandinia_binotata | fish | 0.081 |
| Nasua_nasua | fish | 0.151 |
| Nasuella_olivacea | fish | 0.125 |
| Neofelis_nebulosa | fish | 0.092 |
| Neovison_vison | fish | 0.058 |
| Nyctereutes_procyonoides | fish | 0.069 |
| Otocyon_megalotis | fish | 0.116 |
| Paguma_larvata | fish | 0.081 |
| Panthera_leo | fish | 0.061 |
| Panthera_onca | fish | 0.065 |
| Panthera_pardus | fish | 0.088 |
| Parahyaena_brunnea | fish | 0.048 |
| Pekania_pennanti | fish | 0.076 |
| Potos_flavus | fish | 0.092 |

| Species | Food Item | Pareto-$k$ |
|---|---|---|
| Prionailurus_bengalensis | fish | 0.043 |
| Procyon_cancrivorus | fish | 0.054 |
| Procyon_lotor | fish | 0.049 |
| Proteles_cristatus | fish | 0.076 |
| Pteronura_brasiliensis | fish | 0.084 |
| Puma_concolor | fish | 0.094 |
| Speothos_venaticus | fish | 0.067 |
| Spilogale_putorius | fish | 0.078 |
| Taxidea_taxus | fish | 0.099 |
| Tremarctos_ornatus | fish | 0.073 |
| Uncia_uncia | fish | 0.066 |
| Urocyon_cinereoargenteus | fish | 0.105 |
| Ursus_arctos | fish | 0.067 |
| Ursus_maritimus | fish | 0.088 |
| Viverra_zibetha | fish | 0.070 |
| Viverricula_indica | fish | 0.056 |
| Vulpes_chama | fish | 0.018 |
| Vulpes_lagopus | fish | 0.050 |
| Vulpes_macrotis | fish | 0.030 |
| Vulpes_rueppellii | fish | 0.068 |
| Vulpes_velox | fish | 0.028 |
| Vulpes_vulpes | fish | 0.050 |
| Vulpes_zerda | fish | 0.029 |
| Acinonyx_jubatus | fruit | 0.044 |
| Ailuropoda_melanoleuca | fruit | 0.133 |
| Ailurus_fulgens | fruit | 0.132 |
| Arctictis_binturong | fruit | 0.056 |
| Arctonyx_collaris | fruit | 0.111 |
| Atilax_paludinosus | fruit | 0.096 |
| Bassariscus_astutus | fruit | 0.107 |
| Bdeogale_jacksoni | fruit | 0.102 |
| Canis_adustus | fruit | 0.092 |
| Canis_aureus | fruit | 0.029 |
| Canis_latrans | fruit | 0.050 |
| Canis_lupus | fruit | 0.040 |
| Canis_mesomelas | fruit | 0.060 |
| Canis_simensis | fruit | 0.077 |
| Caracal_caracal | fruit | 0.058 |
| Catopuma_temminckii | fruit | 0.065 |
| Cerdocyon_thous | fruit | 0.044 |
| Chrysocyon_brachyurus | fruit | 0.084 |
| Civettictis_civetta | fruit | 0.076 |
| Conepatus_mesoleucus | fruit | 0.093 |
| Crocuta_crocuta | fruit | 0.088 |
| Cryptoprocta_ferox | fruit | 0.077 |
| Cuon_alpinus | fruit | 0.038 |
| Cynictis_penicillata | fruit | 0.079 |
| Eira_barbara | fruit | 0.120 |

| **Species** | **Food Item** | **Pareto-$k$** |
|---|---|---|
| Enhydra_lutris | fruit | 0.059 |
| Felis_silvestris | fruit | 0.085 |
| Fossa_fossana | fruit | 0.094 |
| Galictis_cuja | fruit | 0.094 |
| Gulo_gulo | fruit | 0.059 |
| Helarctos_malayanus | fruit | 0.106 |
| Helogale_parvula | fruit | 0.110 |
| Hyaena_hyaena | fruit | 0.096 |
| Hydrictis_maculicollis | fruit | 0.060 |
| Ichneumia_albicauda | fruit | 0.071 |
| Leopardus_wiedii | fruit | 0.097 |
| Leptailurus_serval | fruit | 0.072 |
| Lontra_canadensis | fruit | 0.077 |
| Lutra_lutra | fruit | 0.094 |
| Lutrogale_perspicillata | fruit | 0.059 |
| Lycalopex_culpaeus | fruit | 0.040 |
| Lycalopex_gymnocercus | fruit | 0.051 |
| Lycalopex_sechurae | fruit | 0.048 |
| Lycalopex_vetulus | fruit | 0.076 |
| Lycaon_pictus | fruit | 0.083 |
| Lynx_canadensis | fruit | 0.068 |
| Martes_americana | fruit | 0.092 |
| Mellivora_capensis | fruit | 0.088 |
| Melursus_ursinus | fruit | 0.081 |
| Mephitis_mephitis | fruit | 0.109 |
| Mungotictis_decemlineata | fruit | 0.094 |
| Nandinia_binotata | fruit | 0.105 |
| Nasua_nasua | fruit | 0.130 |
| Nasuella_olivacea | fruit | 0.084 |
| Neofelis_nebulosa | fruit | 0.094 |
| Neovison_vison | fruit | 0.147 |
| Nyctereutes_procyonoides | fruit | 0.094 |
| Otocyon_megalotis | fruit | 0.083 |
| Paguma_larvata | fruit | 0.093 |
| Panthera_leo | fruit | 0.063 |
| Panthera_onca | fruit | 0.034 |
| Panthera_pardus | fruit | 0.030 |
| Parahyaena_brunnea | fruit | 0.092 |
| Pekania_pennanti | fruit | 0.083 |
| Potos_flavus | fruit | 0.103 |
| Prionailurus_bengalensis | fruit | 0.072 |
| Procyon_cancrivorus | fruit | 0.067 |
| Procyon_lotor | fruit | 0.078 |
| Proteles_cristatus | fruit | 0.086 |
| Pteronura_brasiliensis | fruit | 0.050 |
| Puma_concolor | fruit | 0.070 |
| Speothos_venaticus | fruit | 0.136 |
| Spilogale_putorius | fruit | 0.113 |

| Species | Food Item | Pareto-$k$ |
|---|---|---|
| Taxidea_taxus | fruit | 0.077 |
| Tremarctos_ornatus | fruit | 0.089 |
| Uncia_uncia | fruit | 0.046 |
| Urocyon_cinereoargenteus | fruit | 0.086 |
| Ursus_arctos | fruit | 0.072 |
| Ursus_maritimus | fruit | 0.106 |
| Viverra_zibetha | fruit | 0.066 |
| Viverricula_indica | fruit | 0.082 |
| Vulpes_chama | fruit | 0.044 |
| Vulpes_lagopus | fruit | 0.049 |
| Vulpes_macrotis | fruit | 0.052 |
| Vulpes_rueppellii | fruit | 0.071 |
| Vulpes_velox | fruit | 0.050 |
| Vulpes_vulpes | fruit | 0.073 |
| Vulpes_zerda | fruit | 0.058 |
| Acinonyx_jubatus | hard_invert | 0.047 |
| Ailuropoda_melanoleuca | hard_invert | 0.092 |
| Ailurus_fulgens | hard_invert | 0.104 |
| Arctictis_binturong | hard_invert | 0.080 |
| Arctonyx_collaris | hard_invert | 0.063 |
| Atilax_paludinosus | hard_invert | 0.045 |
| Bassariscus_astutus | hard_invert | 0.072 |
| Bdeogale_jacksoni | hard_invert | 0.063 |
| Canis_adustus | hard_invert | 0.054 |
| Canis_aureus | hard_invert | 0.030 |
| Canis_latrans | hard_invert | 0.017 |
| Canis_lupus | hard_invert | 0.038 |
| Canis_mesomelas | hard_invert | 0.073 |
| Canis_simensis | hard_invert | 0.045 |
| Caracal_caracal | hard_invert | 0.052 |
| Catopuma_temminckii | hard_invert | 0.063 |
| Cerdocyon_thous | hard_invert | 0.052 |
| Chrysocyon_brachyurus | hard_invert | 0.065 |
| Civettictis_civetta | hard_invert | 0.086 |
| Conepatus_mesoleucus | hard_invert | 0.094 |
| Crocuta_crocuta | hard_invert | 0.077 |
| Cryptoprocta_ferox | hard_invert | 0.122 |
| Cuon_alpinus | hard_invert | 0.062 |
| Cynictis_penicillata | hard_invert | 0.056 |
| Eira_barbara | hard_invert | 0.068 |
| Enhydra_lutris | hard_invert | 0.052 |
| Felis_silvestris | hard_invert | 0.070 |
| Fossa_fossana | hard_invert | 0.072 |
| Galictis_cuja | hard_invert | 0.084 |
| Gulo_gulo | hard_invert | 0.082 |
| Helarctos_malayanus | hard_invert | 0.069 |
| Helogale_parvula | hard_invert | 0.069 |
| Hyaena_hyaena | hard_invert | 0.059 |

| Species | Food Item | Pareto-$k$ |
|---|---|---|
| Hydrictis_maculicollis | hard_invert | 0.059 |
| Ichneumia_albicauda | hard_invert | 0.082 |
| Leopardus_wiedii | hard_invert | 0.076 |
| Leptailurus_serval | hard_invert | 0.069 |
| Lontra_canadensis | hard_invert | 0.067 |
| Lutra_lutra | hard_invert | 0.053 |
| Lutrogale_perspicillata | hard_invert | 0.089 |
| Lycalopex_culpaeus | hard_invert | 0.055 |
| Lycalopex_gymnocercus | hard_invert | 0.031 |
| Lycalopex_sechurae | hard_invert | 0.018 |
| Lycalopex_vetulus | hard_invert | 0.047 |
| Lycaon_pictus | hard_invert | 0.060 |
| Lynx_canadensis | hard_invert | 0.049 |
| Martes_americana | hard_invert | 0.090 |
| Mellivora_capensis | hard_invert | 0.074 |
| Melursus_ursinus | hard_invert | 0.062 |
| Mephitis_mephitis | hard_invert | 0.109 |
| Mungotictis_decemlineata | hard_invert | 0.083 |
| Nandinia_binotata | hard_invert | 0.104 |
| Nasua_nasua | hard_invert | 0.086 |
| Nasuella_olivacea | hard_invert | 0.085 |
| Neofelis_nebulosa | hard_invert | 0.064 |
| Neovison_vison | hard_invert | 0.070 |
| Nyctereutes_procyonoides | hard_invert | 0.065 |
| Otocyon_megalotis | hard_invert | 0.085 |
| Paguma_larvata | hard_invert | 0.086 |
| Panthera_leo | hard_invert | 0.085 |
| Panthera_onca | hard_invert | 0.063 |
| Panthera_pardus | hard_invert | 0.076 |
| Parahyaena_brunnea | hard_invert | 0.080 |
| Pekania_pennanti | hard_invert | 0.059 |
| Potos_flavus | hard_invert | 0.103 |
| Prionailurus_bengalensis | hard_invert | 0.057 |
| Procyon_cancrivorus | hard_invert | 0.088 |
| Procyon_lotor | hard_invert | 0.075 |
| Proteles_cristatus | hard_invert | 0.042 |
| Pteronura_brasiliensis | hard_invert | 0.087 |
| Puma_concolor | hard_invert | 0.041 |
| Speothos_venaticus | hard_invert | 0.095 |
| Spilogale_putorius | hard_invert | 0.069 |
| Taxidea_taxus | hard_invert | 0.110 |
| Tremarctos_ornatus | hard_invert | 0.090 |
| Uncia_uncia | hard_invert | 0.053 |
| Urocyon_cinereoargenteus | hard_invert | 0.071 |
| Ursus_arctos | hard_invert | 0.055 |
| Ursus_maritimus | hard_invert | 0.066 |
| Viverra_zibetha | hard_invert | 0.098 |
| Viverricula_indica | hard_invert | 0.072 |

| **Species** | **Food Item** | **Pareto-*k*** |
|---|---|---|
| Vulpes_chama | hard_invert | 0.045 |
| Vulpes_lagopus | hard_invert | 0.039 |
| Vulpes_macrotis | hard_invert | 0.055 |
| Vulpes_rueppellii | hard_invert | 0.040 |
| Vulpes_velox | hard_invert | 0.043 |
| Vulpes_vulpes | hard_invert | 0.050 |
| Vulpes_zerda | hard_invert | 0.041 |
| Acinonyx_jubatus | herptile | 0.056 |
| Ailuropoda_melanoleuca | herptile | 0.057 |
| Ailurus_fulgens | herptile | 0.065 |
| Arctictis_binturong | herptile | 0.067 |
| Arctonyx_collaris | herptile | 0.061 |
| Atilax_paludinosus | herptile | 0.056 |
| Bassariscus_astutus | herptile | 0.065 |
| Bdeogale_jacksoni | herptile | 0.073 |
| Canis_adustus | herptile | 0.029 |
| Canis_aureus | herptile | 0.020 |
| Canis_latrans | herptile | 0.016 |
| Canis_lupus | herptile | 0.022 |
| Canis_mesomelas | herptile | 0.018 |
| Canis_simensis | herptile | 0.050 |
| Caracal_caracal | herptile | 0.040 |
| Catopuma_temminckii | herptile | 0.015 |
| Cerdocyon_thous | herptile | 0.038 |
| Chrysocyon_brachyurus | herptile | 0.022 |
| Civettictis_civetta | herptile | 0.074 |
| Conepatus_mesoleucus | herptile | 0.065 |
| Crocuta_crocuta | herptile | 0.046 |
| Cryptoprocta_ferox | herptile | 0.056 |
| Cuon_alpinus | herptile | 0.035 |
| Cynictis_penicillata | herptile | 0.055 |
| Eira_barbara | herptile | 0.082 |
| Enhydra_lutris | herptile | 0.065 |
| Felis_silvestris | herptile | 0.039 |
| Fossa_fossana | herptile | 0.074 |
| Galictis_cuja | herptile | 0.036 |
| Gulo_gulo | herptile | 0.067 |
| Helarctos_malayanus | herptile | 0.043 |
| Helogale_parvula | herptile | 0.048 |
| Hyaena_hyaena | herptile | 0.056 |
| Hydrictis_maculicollis | herptile | 0.042 |
| Ichneumia_albicauda | herptile | 0.060 |
| Leopardus_wiedii | herptile | 0.012 |
| Leptailurus_serval | herptile | 0.038 |
| Lontra_canadensis | herptile | 0.039 |
| Lutra_lutra | herptile | 0.053 |
| Lutrogale_perspicillata | herptile | 0.028 |
| Lycalopex_culpaeus | herptile | 0.012 |

| Species | Food Item | Pareto-$k$ |
|---|---|---|
| Lycalopex_gymnocercus | herptile | 0.018 |
| Lycalopex_sechurae | herptile | 0.038 |
| Lycalopex_vetulus | herptile | 0.026 |
| Lycaon_pictus | herptile | 0.051 |
| Lynx_canadensis | herptile | 0.057 |
| Martes_americana | herptile | 0.041 |
| Mellivora_capensis | herptile | 0.032 |
| Melursus_ursinus | herptile | 0.060 |
| Mephitis_mephitis | herptile | 0.102 |
| Mungotictis_decemlineata | herptile | 0.050 |
| Nandinia_binotata | herptile | 0.061 |
| Nasua_nasua | herptile | 0.069 |
| Nasuella_olivacea | herptile | 0.071 |
| Neofelis_nebulosa | herptile | 0.051 |
| Neovison_vison | herptile | 0.033 |
| Nyctereutes_procyonoides | herptile | 0.043 |
| Otocyon_megalotis | herptile | 0.066 |
| Paguma_larvata | herptile | 0.070 |
| Panthera_leo | herptile | 0.051 |
| Panthera_onca | herptile | 0.066 |
| Panthera_pardus | herptile | 0.019 |
| Parahyaena_brunnea | herptile | 0.038 |
| Pekania_pennanti | herptile | 0.056 |
| Potos_flavus | herptile | 0.079 |
| Prionailurus_bengalensis | herptile | 0.062 |
| Procyon_cancrivorus | herptile | 0.057 |
| Procyon_lotor | herptile | 0.050 |
| Proteles_cristatus | herptile | 0.045 |
| Pteronura_brasiliensis | herptile | 0.074 |
| Puma_concolor | herptile | 0.017 |
| Speothos_venaticus | herptile | 0.044 |
| Spilogale_putorius | herptile | 0.031 |
| Taxidea_taxus | herptile | 0.069 |
| Tremarctos_ornatus | herptile | 0.055 |
| Uncia_uncia | herptile | 0.051 |
| Urocyon_cinereoargenteus | herptile | 0.029 |
| Ursus_arctos | herptile | 0.040 |
| Ursus_maritimus | herptile | 0.057 |
| Viverra_zibetha | herptile | 0.062 |
| Viverricula_indica | herptile | 0.047 |
| Vulpes_chama | herptile | 0.031 |
| Vulpes_lagopus | herptile | 0.046 |
| Vulpes_macrotis | herptile | 0.032 |
| Vulpes_rueppellii | herptile | 0.034 |
| Vulpes_velox | herptile | 0.017 |
| Vulpes_vulpes | herptile | 0.047 |
| Vulpes_zerda | herptile | 0.016 |
| Acinonyx_jubatus | large_mammal | 0.047 |

| Species | Food Item | Pareto-$k$ |
|---|---|---|
| Ailuropoda_melanoleuca | large_mammal | 0.046 |
| Ailurus_fulgens | large_mammal | 0.084 |
| Arctictis_binturong | large_mammal | 0.071 |
| Arctonyx_collaris | large_mammal | 0.086 |
| Atilax_paludinosus | large_mammal | 0.059 |
| Bassariscus_astutus | large_mammal | 0.088 |
| Bdeogale_jacksoni | large_mammal | 0.070 |
| Canis_adustus | large_mammal | 0.031 |
| Canis_aureus | large_mammal | 0.043 |
| Canis_latrans | large_mammal | 0.036 |
| Canis_lupus | large_mammal | 0.041 |
| Canis_mesomelas | large_mammal | 0.061 |
| Canis_simensis | large_mammal | 0.062 |
| Caracal_caracal | large_mammal | 0.049 |
| Catopuma_temminckii | large_mammal | 0.072 |
| Cerdocyon_thous | large_mammal | 0.054 |
| Chrysocyon_brachyurus | large_mammal | 0.053 |
| Civettictis_civetta | large_mammal | 0.052 |
| Conepatus_mesoleucus | large_mammal | 0.093 |
| Crocuta_crocuta | large_mammal | 0.065 |
| Cryptoprocta_ferox | large_mammal | 0.062 |
| Cuon_alpinus | large_mammal | 0.032 |
| Cynictis_penicillata | large_mammal | 0.084 |
| Eira_barbara | large_mammal | 0.051 |
| Enhydra_lutris | large_mammal | 0.065 |
| Felis_silvestris | large_mammal | 0.040 |
| Fossa_fossana | large_mammal | 0.074 |
| Galictis_cuja | large_mammal | 0.092 |
| Gulo_gulo | large_mammal | 0.065 |
| Helarctos_malayanus | large_mammal | 0.056 |
| Helogale_parvula | large_mammal | 0.093 |
| Hyaena_hyaena | large_mammal | 0.058 |
| Hydrictis_maculicollis | large_mammal | 0.052 |
| Ichneumia_albicauda | large_mammal | 0.069 |
| Leopardus_wiedii | large_mammal | 0.057 |
| Leptailurus_serval | large_mammal | 0.048 |
| Lontra_canadensis | large_mammal | 0.042 |
| Lutra_lutra | large_mammal | 0.053 |
| Lutrogale_perspicillata | large_mammal | 0.026 |
| Lycalopex_culpaeus | large_mammal | 0.051 |
| Lycalopex_gymnocercus | large_mammal | 0.037 |
| Lycalopex_sechurae | large_mammal | 0.037 |
| Lycalopex_vetulus | large_mammal | 0.041 |
| Lycaon_pictus | large_mammal | 0.051 |
| Lynx_canadensis | large_mammal | 0.063 |
| Martes_americana | large_mammal | 0.063 |
| Mellivora_capensis | large_mammal | 0.076 |
| Melursus_ursinus | large_mammal | 0.054 |

| Species | Food Item | Pareto-$k$ |
|---|---|---|
| Mephitis_mephitis | large_mammal | 0.076 |
| Mungotictis_decemlineata | large_mammal | 0.087 |
| Nandinia_binotata | large_mammal | 0.094 |
| Nasua_nasua | large_mammal | 0.087 |
| Nasuella_olivacea | large_mammal | 0.086 |
| Neofelis_nebulosa | large_mammal | 0.059 |
| Neovison_vison | large_mammal | 0.051 |
| Nyctereutes_procyonoides | large_mammal | 0.054 |
| Otocyon_megalotis | large_mammal | 0.061 |
| Paguma_larvata | large_mammal | 0.077 |
| Panthera_leo | large_mammal | 0.049 |
| Panthera_onca | large_mammal | 0.058 |
| Panthera_pardus | large_mammal | 0.030 |
| Parahyaena_brunnea | large_mammal | 0.061 |
| Pekania_pennanti | large_mammal | 0.058 |
| Potos_flavus | large_mammal | 0.108 |
| Prionailurus_bengalensis | large_mammal | 0.061 |
| Procyon_cancrivorus | large_mammal | 0.080 |
| Procyon_lotor | large_mammal | 0.058 |
| Proteles_cristatus | large_mammal | 0.087 |
| Pteronura_brasiliensis | large_mammal | 0.074 |
| Puma_concolor | large_mammal | 0.046 |
| Speothos_venaticus | large_mammal | 0.038 |
| Spilogale_putorius | large_mammal | 0.085 |
| Taxidea_taxus | large_mammal | 0.071 |
| Tremarctos_ornatus | large_mammal | 0.100 |
| Uncia_uncia | large_mammal | 0.050 |
| Urocyon_cinereoargenteus | large_mammal | 0.042 |
| Ursus_arctos | large_mammal | 0.068 |
| Ursus_maritimus | large_mammal | 0.069 |
| Viverra_zibetha | large_mammal | 0.066 |
| Viverricula_indica | large_mammal | 0.062 |
| Vulpes_chama | large_mammal | 0.018 |
| Vulpes_lagopus | large_mammal | 0.078 |
| Vulpes_macrotis | large_mammal | 0.073 |
| Vulpes_rueppellii | large_mammal | 0.024 |
| Vulpes_velox | large_mammal | 0.048 |
| Vulpes_vulpes | large_mammal | 0.059 |
| Vulpes_zerda | large_mammal | 0.029 |
| Acinonyx_jubatus | plant | 0.059 |
| Ailuropoda_melanoleuca | plant | 0.096 |
| Ailurus_fulgens | plant | 0.073 |
| Arctictis_binturong | plant | 0.075 |
| Arctonyx_collaris | plant | 0.115 |
| Atilax_paludinosus | plant | 0.058 |
| Bassariscus_astutus | plant | 0.086 |
| Bdeogale_jacksoni | plant | 0.059 |
| Canis_adustus | plant | 0.063 |

| Species | Food Item | Pareto-*k* |
|---|---|---|
| Canis_aureus | plant | 0.055 |
| Canis_latrans | plant | 0.052 |
| Canis_lupus | plant | 0.062 |
| Canis_mesomelas | plant | 0.055 |
| Canis_simensis | plant | 0.033 |
| Caracal_caracal | plant | 0.087 |
| Catopuma_temminckii | plant | 0.089 |
| Cerdocyon_thous | plant | 0.065 |
| Chrysocyon_brachyurus | plant | 0.076 |
| Civettictis_civetta | plant | 0.079 |
| Conepatus_mesoleucus | plant | 0.085 |
| Crocuta_crocuta | plant | 0.136 |
| Cryptoprocta_ferox | plant | 0.070 |
| Cuon_alpinus | plant | 0.040 |
| Cynictis_penicillata | plant | 0.100 |
| Eira_barbara | plant | 0.061 |
| Enhydra_lutris | plant | 0.042 |
| Felis_silvestris | plant | 0.059 |
| Fossa_fossana | plant | 0.083 |
| Galictis_cuja | plant | 0.061 |
| Gulo_gulo | plant | 0.052 |
| Helarctos_malayanus | plant | 0.071 |
| Helogale_parvula | plant | 0.065 |
| Hyaena_hyaena | plant | 0.073 |
| Hydrictis_maculicollis | plant | 0.061 |
| Ichneumia_albicauda | plant | 0.073 |
| Leopardus_wiedii | plant | 0.096 |
| Leptailurus_serval | plant | 0.078 |
| Lontra_canadensis | plant | 0.072 |
| Lutra_lutra | plant | 0.046 |
| Lutrogale_perspicillata | plant | 0.052 |
| Lycalopex_culpaeus | plant | 0.045 |
| Lycalopex_gymnocercus | plant | 0.042 |
| Lycalopex_sechurae | plant | 0.042 |
| Lycalopex_vetulus | plant | 0.034 |
| Lycaon_pictus | plant | 0.089 |
| Lynx_canadensis | plant | 0.046 |
| Martes_americana | plant | 0.052 |
| Mellivora_capensis | plant | 0.085 |
| Melursus_ursinus | plant | 0.064 |
| Mephitis_mephitis | plant | 0.111 |
| Mungotictis_decemlineata | plant | 0.080 |
| Nandinia_binotata | plant | 0.078 |
| Nasua_nasua | plant | 0.082 |
| Nasuella_olivacea | plant | 0.070 |
| Neofelis_nebulosa | plant | 0.069 |
| Neovison_vison | plant | 0.089 |
| Nyctereutes_procyonoides | plant | 0.073 |

| Species | Food Item | Pareto-$k$ |
|---|---|---|
| Otocyon_megalotis | plant | 0.065 |
| Paguma_larvata | plant | 0.089 |
| Panthera_leo | plant | 0.074 |
| Panthera_onca | plant | 0.048 |
| Panthera_pardus | plant | 0.047 |
| Parahyaena_brunnea | plant | 0.080 |
| Pekania_pennanti | plant | 0.056 |
| Potos_flavus | plant | 0.121 |
| Prionailurus_bengalensis | plant | 0.072 |
| Procyon_cancrivorus | plant | 0.073 |
| Procyon_lotor | plant | 0.081 |
| Proteles_cristatus | plant | 0.078 |
| Pteronura_brasiliensis | plant | 0.056 |
| Puma_concolor | plant | 0.058 |
| Speothos_venaticus | plant | 0.030 |
| Spilogale_putorius | plant | 0.116 |
| Taxidea_taxus | plant | 0.091 |
| Tremarctos_ornatus | plant | 0.089 |
| Uncia_uncia | plant | 0.053 |
| Urocyon_cinereoargenteus | plant | 0.069 |
| Ursus_arctos | plant | 0.080 |
| Ursus_maritimus | plant | 0.060 |
| Viverra_zibetha | plant | 0.090 |
| Viverricula_indica | plant | 0.102 |
| Vulpes_chama | plant | 0.050 |
| Vulpes_lagopus | plant | 0.039 |
| Vulpes_macrotis | plant | 0.051 |
| Vulpes_rueppellii | plant | 0.069 |
| Vulpes_velox | plant | 0.049 |
| Vulpes_vulpes | plant | 0.048 |
| Vulpes_zerda | plant | 0.053 |
| Acinonyx_jubatus | root | 0.055 |
| Ailuropoda_melanoleuca | root | 0.079 |
| Ailurus_fulgens | root | 0.077 |
| Arctictis_binturong | root | 0.046 |
| Arctonyx_collaris | root | 0.043 |
| Atilax_paludinosus | root | 0.057 |
| Bassariscus_astutus | root | 0.099 |
| Bdeogale_jacksoni | root | 0.071 |
| Canis_adustus | root | 0.050 |
| Canis_aureus | root | 0.047 |
| Canis_latrans | root | 0.029 |
| Canis_lupus | root | 0.052 |
| Canis_mesomelas | root | 0.058 |
| Canis_simensis | root | 0.033 |
| Caracal_caracal | root | 0.057 |
| Catopuma_temminckii | root | 0.054 |
| Cerdocyon_thous | root | 0.048 |

| Species | Food Item | Pareto-$k$ |
|---|---|---|
| Chrysocyon_brachyurus | root | 0.061 |
| Civettictis_civetta | root | 0.093 |
| Conepatus_mesoleucus | root | 0.092 |
| Crocuta_crocuta | root | 0.053 |
| Cryptoprocta_ferox | root | 0.071 |
| Cuon_alpinus | root | 0.061 |
| Cynictis_penicillata | root | 0.042 |
| Eira_barbara | root | 0.055 |
| Enhydra_lutris | root | 0.097 |
| Felis_silvestris | root | 0.056 |
| Fossa_fossana | root | 0.058 |
| Galictis_cuja | root | 0.065 |
| Gulo_gulo | root | 0.056 |
| Helarctos_malayanus | root | 0.057 |
| Helogale_parvula | root | 0.035 |
| Hyaena_hyaena | root | 0.039 |
| Hydrictis_maculicollis | root | 0.054 |
| Ichneumia_albicauda | root | 0.065 |
| Leopardus_wiedii | root | 0.026 |
| Leptailurus_serval | root | 0.039 |
| Lontra_canadensis | root | 0.048 |
| Lutra_lutra | root | 0.062 |
| Lutrogale_perspicillata | root | 0.057 |
| Lycalopex_culpaeus | root | 0.069 |
| Lycalopex_gymnocercus | root | 0.038 |
| Lycalopex_sechurae | root | 0.030 |
| Lycalopex_vetulus | root | 0.023 |
| Lycaon_pictus | root | 0.071 |
| Lynx_canadensis | root | 0.050 |
| Martes_americana | root | 0.068 |
| Mellivora_capensis | root | 0.066 |
| Melursus_ursinus | root | 0.048 |
| Mephitis_mephitis | root | 0.056 |
| Mungotictis_decemlineata | root | 0.060 |
| Nandinia_binotata | root | 0.102 |
| Nasua_nasua | root | 0.058 |
| Nasuella_olivacea | root | 0.092 |
| Neofelis_nebulosa | root | 0.035 |
| Neovison_vison | root | 0.034 |
| Nyctereutes_procyonoides | root | 0.080 |
| Otocyon_megalotis | root | 0.060 |
| Paguma_larvata | root | 0.093 |
| Panthera_leo | root | 0.071 |
| Panthera_onca | root | 0.061 |
| Panthera_pardus | root | 0.049 |
| Parahyaena_brunnea | root | 0.033 |
| Pekania_pennanti | root | 0.055 |
| Potos_flavus | root | 0.078 |

| Species | Food Item | Pareto-$k$ |
|---|---|---|
| Prionailurus_bengalensis | root | 0.032 |
| Procyon_cancrivorus | root | 0.092 |
| Procyon_lotor | root | 0.080 |
| Proteles_cristatus | root | 0.052 |
| Pteronura_brasiliensis | root | 0.045 |
| Puma_concolor | root | 0.056 |
| Speothos_venaticus | root | 0.040 |
| Spilogale_putorius | root | 0.080 |
| Taxidea_taxus | root | 0.098 |
| Tremarctos_ornatus | root | 0.087 |
| Uncia_uncia | root | 0.014 |
| Urocyon_cinereoargenteus | root | 0.103 |
| Ursus_arctos | root | 0.063 |
| Ursus_maritimus | root | 0.071 |
| Viverra_zibetha | root | 0.088 |
| Viverricula_indica | root | 0.067 |
| Vulpes_chama | root | 0.049 |
| Vulpes_lagopus | root | 0.079 |
| Vulpes_macrotis | root | 0.031 |
| Vulpes_rueppellii | root | 0.062 |
| Vulpes_velox | root | 0.029 |
| Vulpes_vulpes | root | 0.051 |
| Vulpes_zerda | root | 0.105 |
| Acinonyx_jubatus | seed | 0.045 |
| Ailuropoda_melanoleuca | seed | 0.075 |
| Ailurus_fulgens | seed | 0.074 |
| Arctictis_binturong | seed | 0.057 |
| Arctonyx_collaris | seed | 0.092 |
| Atilax_paludinosus | seed | 0.091 |
| Bassariscus_astutus | seed | 0.076 |
| Bdeogale_jacksoni | seed | 0.028 |
| Canis_adustus | seed | 0.046 |
| Canis_aureus | seed | 0.058 |
| Canis_latrans | seed | 0.043 |
| Canis_lupus | seed | 0.049 |
| Canis_mesomelas | seed | 0.034 |
| Canis_simensis | seed | 0.036 |
| Caracal_caracal | seed | 0.037 |
| Catopuma_temminckii | seed | 0.051 |
| Cerdocyon_thous | seed | 0.044 |
| Chrysocyon_brachyurus | seed | 0.043 |
| Civettictis_civetta | seed | 0.048 |
| Conepatus_mesoleucus | seed | 0.080 |
| Crocuta_crocuta | seed | 0.070 |
| Cryptoprocta_ferox | seed | 0.079 |
| Cuon_alpinus | seed | 0.034 |
| Cynictis_penicillata | seed | 0.087 |
| Eira_barbara | seed | 0.063 |

| Species | Food Item | Pareto-*k* |
|---|---|---|
| Enhydra_lutris | seed | 0.037 |
| Felis_silvestris | seed | 0.049 |
| Fossa_fossana | seed | 0.063 |
| Galictis_cuja | seed | 0.047 |
| Gulo_gulo | seed | 0.067 |
| Helarctos_malayanus | seed | 0.053 |
| Helogale_parvula | seed | 0.063 |
| Hyaena_hyaena | seed | 0.058 |
| Hydrictis_maculicollis | seed | 0.024 |
| Ichneumia_albicauda | seed | 0.064 |
| Leopardus_wiedii | seed | 0.041 |
| Leptailurus_serval | seed | 0.034 |
| Lontra_canadensis | seed | 0.059 |
| Lutra_lutra | seed | 0.054 |
| Lutrogale_perspicillata | seed | 0.046 |
| Lycalopex_culpaeus | seed | 0.035 |
| Lycalopex_gymnocercus | seed | 0.047 |
| Lycalopex_sechurae | seed | 0.051 |
| Lycalopex_vetulus | seed | 0.025 |
| Lycaon_pictus | seed | 0.056 |
| Lynx_canadensis | seed | 0.064 |
| Martes_americana | seed | 0.060 |
| Mellivora_capensis | seed | 0.080 |
| Melursus_ursinus | seed | 0.066 |
| Mephitis_mephitis | seed | 0.066 |
| Mungotictis_decemlineata | seed | 0.047 |
| Nandinia_binotata | seed | 0.093 |
| Nasua_nasua | seed | 0.071 |
| Nasuella_olivacea | seed | 0.073 |
| Neofelis_nebulosa | seed | 0.041 |
| Neovison_vison | seed | 0.056 |
| Nyctereutes_procyonoides | seed | 0.059 |
| Otocyon_megalotis | seed | 0.043 |
| Paguma_larvata | seed | 0.059 |
| Panthera_leo | seed | 0.069 |
| Panthera_onca | seed | 0.041 |
| Panthera_pardus | seed | 0.039 |
| Parahyaena_brunnea | seed | 0.063 |
| Pekania_pennanti | seed | 0.049 |
| Potos_flavus | seed | 0.070 |
| Prionailurus_bengalensis | seed | 0.045 |
| Procyon_cancrivorus | seed | 0.058 |
| Procyon_lotor | seed | 0.116 |
| Proteles_cristatus | seed | 0.056 |
| Pteronura_brasiliensis | seed | 0.046 |
| Puma_concolor | seed | 0.040 |
| Speothos_venaticus | seed | 0.053 |
| Spilogale_putorius | seed | 0.057 |

| Species | Food Item | Pareto-$k$ |
|---|---|---|
| Taxidea_taxus | seed | 0.066 |
| Tremarctos_ornatus | seed | 0.041 |
| Uncia_uncia | seed | 0.048 |
| Urocyon_cinereoargenteus | seed | 0.047 |
| Ursus_arctos | seed | 0.091 |
| Ursus_maritimus | seed | 0.055 |
| Viverra_zibetha | seed | 0.045 |
| Viverricula_indica | seed | 0.075 |
| Vulpes_chama | seed | 0.041 |
| Vulpes_lagopus | seed | 0.061 |
| Vulpes_macrotis | seed | 0.045 |
| Vulpes_rueppellii | seed | 0.036 |
| Vulpes_velox | seed | 0.048 |
| Vulpes_vulpes | seed | 0.041 |
| Vulpes_zerda | seed | 0.062 |
| Acinonyx_jubatus | small_mammal | 0.079 |
| Ailuropoda_melanoleuca | small_mammal | 0.077 |
| Ailurus_fulgens | small_mammal | 0.080 |
| Arctictis_binturong | small_mammal | 0.101 |
| Arctonyx_collaris | small_mammal | 0.067 |
| Atilax_paludinosus | small_mammal | 0.073 |
| Bassariscus_astutus | small_mammal | 0.068 |
| Bdeogale_jacksoni | small_mammal | 0.088 |
| Canis_adustus | small_mammal | 0.042 |
| Canis_aureus | small_mammal | 0.048 |
| Canis_latrans | small_mammal | 0.019 |
| Canis_lupus | small_mammal | 0.048 |
| Canis_mesomelas | small_mammal | 0.054 |
| Canis_simensis | small_mammal | 0.046 |
| Caracal_caracal | small_mammal | 0.065 |
| Catopuma_temminckii | small_mammal | 0.025 |
| Cerdocyon_thous | small_mammal | 0.036 |
| Chrysocyon_brachyurus | small_mammal | 0.045 |
| Civettictis_civetta | small_mammal | 0.097 |
| Conepatus_mesoleucus | small_mammal | 0.051 |
| Crocuta_crocuta | small_mammal | 0.034 |
| Cryptoprocta_ferox | small_mammal | 0.065 |
| Cuon_alpinus | small_mammal | 0.037 |
| Cynictis_penicillata | small_mammal | 0.076 |
| Eira_barbara | small_mammal | 0.065 |
| Enhydra_lutris | small_mammal | 0.107 |
| Felis_silvestris | small_mammal | 0.035 |
| Fossa_fossana | small_mammal | 0.063 |
| Galictis_cuja | small_mammal | 0.061 |
| Gulo_gulo | small_mammal | 0.088 |
| Helarctos_malayanus | small_mammal | 0.041 |
| Helogale_parvula | small_mammal | 0.039 |
| Hyaena_hyaena | small_mammal | 0.020 |

| Species | Food Item | Pareto-$k$ |
|---|---|---|
| Hydrictis_maculicollis | small_mammal | 0.055 |
| Ichneumia_albicauda | small_mammal | 0.064 |
| Leopardus_wiedii | small_mammal | 0.032 |
| Leptailurus_serval | small_mammal | 0.033 |
| Lontra_canadensis | small_mammal | 0.075 |
| Lutra_lutra | small_mammal | 0.075 |
| Lutrogale_perspicillata | small_mammal | 0.077 |
| Lycalopex_culpaeus | small_mammal | 0.019 |
| Lycalopex_gymnocercus | small_mammal | 0.031 |
| Lycalopex_sechurae | small_mammal | 0.027 |
| Lycalopex_vetulus | small_mammal | 0.055 |
| Lycaon_pictus | small_mammal | 0.045 |
| Lynx_canadensis | small_mammal | 0.035 |
| Martes_americana | small_mammal | 0.057 |
| Mellivora_capensis | small_mammal | 0.074 |
| Melursus_ursinus | small_mammal | 0.065 |
| Mephitis_mephitis | small_mammal | 0.056 |
| Mungotictis_decemlineata | small_mammal | 0.075 |
| Nandinia_binotata | small_mammal | 0.070 |
| Nasua_nasua | small_mammal | 0.038 |
| Nasuella_olivacea | small_mammal | 0.078 |
| Neofelis_nebulosa | small_mammal | 0.014 |
| Neovison_vison | small_mammal | 0.059 |
| Nyctereutes_procyonoides | small_mammal | 0.068 |
| Otocyon_megalotis | small_mammal | 0.095 |
| Paguma_larvata | small_mammal | 0.101 |
| Panthera_leo | small_mammal | 0.054 |
| Panthera_onca | small_mammal | 0.022 |
| Panthera_pardus | small_mammal | 0.053 |
| Parahyaena_brunnea | small_mammal | 0.089 |
| Pekania_pennanti | small_mammal | 0.046 |
| Potos_flavus | small_mammal | 0.091 |
| Prionailurus_bengalensis | small_mammal | 0.049 |
| Procyon_cancrivorus | small_mammal | 0.075 |
| Procyon_lotor | small_mammal | 0.095 |
| Proteles_cristatus | small_mammal | 0.056 |
| Pteronura_brasiliensis | small_mammal | 0.060 |
| Puma_concolor | small_mammal | 0.010 |
| Speothos_venaticus | small_mammal | 0.039 |
| Spilogale_putorius | small_mammal | 0.067 |
| Taxidea_taxus | small_mammal | 0.096 |
| Tremarctos_ornatus | small_mammal | 0.056 |
| Uncia_uncia | small_mammal | 0.067 |
| Urocyon_cinereoargenteus | small_mammal | 0.062 |
| Ursus_arctos | small_mammal | 0.034 |
| Ursus_maritimus | small_mammal | 0.026 |
| Viverra_zibetha | small_mammal | 0.065 |
| Viverricula_indica | small_mammal | 0.062 |

| **Species** | **Food Item** | **Pareto-*k*** |
|---|---|---|
| Vulpes_chama | small_mammal | 0.027 |
| Vulpes_lagopus | small_mammal | 0.064 |
| Vulpes_macrotis | small_mammal | 0.042 |
| Vulpes_rueppellii | small_mammal | 0.033 |
| Vulpes_velox | small_mammal | 0.044 |
| Vulpes_vulpes | small_mammal | 0.050 |
| Vulpes_zerda | small_mammal | 0.037 |
| Acinonyx_jubatus | soft_invert | 0.116 |
| Ailuropoda_melanoleuca | soft_invert | 0.106 |
| Ailurus_fulgens | soft_invert | 0.114 |
| Arctictis_binturong | soft_invert | 0.114 |
| Arctonyx_collaris | soft_invert | 0.099 |
| Atilax_paludinosus | soft_invert | 0.104 |
| Bassariscus_astutus | soft_invert | 0.168 |
| Bdeogale_jacksoni | soft_invert | 0.138 |
| Canis_adustus | soft_invert | 0.104 |
| Canis_aureus | soft_invert | 0.094 |
| Canis_latrans | soft_invert | 0.130 |
| Canis_lupus | soft_invert | 0.062 |
| Canis_mesomelas | soft_invert | 0.082 |
| Canis_simensis | soft_invert | 0.063 |
| Caracal_caracal | soft_invert | 0.069 |
| Catopuma_temminckii | soft_invert | 0.059 |
| Cerdocyon_thous | soft_invert | 0.070 |
| Chrysocyon_brachyurus | soft_invert | 0.106 |
| Civettictis_civetta | soft_invert | 0.096 |
| Conepatus_mesoleucus | soft_invert | 0.085 |
| Crocuta_crocuta | soft_invert | 0.096 |
| Cryptoprocta_ferox | soft_invert | 0.185 |
| Cuon_alpinus | soft_invert | 0.053 |
| Cynictis_penicillata | soft_invert | 0.076 |
| Eira_barbara | soft_invert | 0.089 |
| Enhydra_lutris | soft_invert | 0.180 |
| Felis_silvestris | soft_invert | 0.089 |
| Fossa_fossana | soft_invert | 0.196 |
| Galictis_cuja | soft_invert | 0.109 |
| Gulo_gulo | soft_invert | 0.118 |
| Helarctos_malayanus | soft_invert | 0.109 |
| Helogale_parvula | soft_invert | 0.111 |
| Hyaena_hyaena | soft_invert | 0.073 |
| Hydrictis_maculicollis | soft_invert | 0.070 |
| Ichneumia_albicauda | soft_invert | 0.117 |
| Leopardus_wiedii | soft_invert | 0.057 |
| Leptailurus_serval | soft_invert | 0.071 |
| Lontra_canadensis | soft_invert | 0.105 |
| Lutra_lutra | soft_invert | 0.073 |
| Lutrogale_perspicillata | soft_invert | 0.129 |
| Lycalopex_culpaeus | soft_invert | 0.053 |

| Species | Food Item | Pareto-$k$ |
|---|---|---|
| Lycalopex_gymnocercus | soft_invert | 0.084 |
| Lycalopex_sechurae | soft_invert | 0.083 |
| Lycalopex_vetulus | soft_invert | 0.069 |
| Lycaon_pictus | soft_invert | 0.071 |
| Lynx_canadensis | soft_invert | 0.067 |
| Martes_americana | soft_invert | 0.087 |
| Mellivora_capensis | soft_invert | 0.107 |
| Melursus_ursinus | soft_invert | 0.111 |
| Mephitis_mephitis | soft_invert | 0.119 |
| Mungotictis_decemlineata | soft_invert | 0.142 |
| Nandinia_binotata | soft_invert | 0.189 |
| Nasua_nasua | soft_invert | 0.138 |
| Nasuella_olivacea | soft_invert | 0.098 |
| Neofelis_nebulosa | soft_invert | 0.067 |
| Neovison_vison | soft_invert | 0.116 |
| Nyctereutes_procyonoides | soft_invert | 0.117 |
| Otocyon_megalotis | soft_invert | 0.083 |
| Paguma_larvata | soft_invert | 0.142 |
| Panthera_leo | soft_invert | 0.116 |
| Panthera_onca | soft_invert | 0.112 |
| Panthera_pardus | soft_invert | 0.110 |
| Parahyaena_brunnea | soft_invert | 0.148 |
| Pekania_pennanti | soft_invert | 0.113 |
| Potos_flavus | soft_invert | 0.148 |
| Prionailurus_bengalensis | soft_invert | 0.169 |
| Procyon_cancrivorus | soft_invert | 0.100 |
| Procyon_lotor | soft_invert | 0.093 |
| Proteles_cristatus | soft_invert | 0.130 |
| Pteronura_brasiliensis | soft_invert | 0.137 |
| Puma_concolor | soft_invert | 0.090 |
| Speothos_venaticus | soft_invert | 0.116 |
| Spilogale_putorius | soft_invert | 0.103 |
| Taxidea_taxus | soft_invert | 0.149 |
| Tremarctos_ornatus | soft_invert | 0.134 |
| Uncia_uncia | soft_invert | 0.097 |
| Urocyon_cinereoargenteus | soft_invert | 0.124 |
| Ursus_arctos | soft_invert | 0.111 |
| Ursus_maritimus | soft_invert | 0.125 |
| Viverra_zibetha | soft_invert | 0.143 |
| Viverricula_indica | soft_invert | 0.122 |
| Vulpes_chama | soft_invert | 0.058 |
| Vulpes_lagopus | soft_invert | 0.096 |
| Vulpes_macrotis | soft_invert | 0.056 |
| Vulpes_rueppellii | soft_invert | 0.099 |
| Vulpes_velox | soft_invert | 0.082 |
| Vulpes_vulpes | soft_invert | 0.053 |
| Vulpes_zerda | soft_invert | 0.095 |

Table S3: Ordinal regression model outputs. Each cell provides the mean value with the 89% credible intervals in parentheses. Cells are NA if the model did not include the parameter. This table is available in the Dryad repo and on GitHub as `Supp_model_results_table.csv`.

| **Metric** | **Parameter** | **Bird** | **Hard Invert** | **Large Mammal** | **Plant** | **Carrion** | **Egg** | **Herptile** | **Seed** | **Small Mammal** | **Soft Invert** | **Fish** | **Fruit** | **Root** |
|---|---|---|---|---|---|---|---|---|---|---|---|---|---|---|
| ALL | Intercept$_1$ | NA | NA | 0.48 (-0.38, 1.23) | -0.74 (-2.11, 0.45) | NA | NA | NA | NA | NA | NA | NA | NA | NA |
| ALL | Intercept$_2$ | NA | NA | 1.97 (1.12, 2.82) | 1.96 (0.66, 3.21) | NA | NA | NA | NA | NA | NA | NA | NA | NA |
| ALL | Intercept$_3$ | NA | NA | 2.88 (1.95, 3.86) | 3.62 (2.23, 5.13) | NA | NA | NA | NA | NA | NA | NA | NA | NA |
| ALL | Mass | NA | NA | 2.28 (1.49, 3.09) | 0.24 (-0.42, 0.92) | NA | NA | NA | NA | NA | NA | NA | NA | NA |
| ALL | Phylo$_s d$ | NA | NA | 0.82 (0.08, 1.92) | 2.01 (0.89, 3.1) | NA | NA | NA | NA | NA | NA | NA | NA | NA |
| ALL | RLGA | NA | NA | -1.57 (-2.63, -0.48) | 0.26 (-0.67, 1.2) | NA | NA | NA | NA | NA | NA | NA | NA | NA |
| ALL | RLGA:Mass | NA | NA | -0.01 (-1.11, 1.08) | -0.34 (-1.32, 0.67) | NA | NA | NA | NA | NA | NA | NA | NA | NA |
| ALL | m1$_R FI$ | NA | NA | 0.22 (-0.46, 0.9) | 0.38 (-0.29, 1.06) | NA | NA | NA | NA | NA | NA | NA | NA | NA |
| ALL | m1$_R FI$ : *Mass* | NA | NA | 0.71 (-0.03, 1.47) | 0.44 (-0.17, 1.06) | NA | NA | NA | NA | NA | NA | NA | NA | NA |
| ALL | m2$_R FI$ | NA | NA | 0.68 (-0.3, 1.65) | -0.17 (-1.15, 0.83) | NA | NA | NA | NA | NA | NA | NA | NA | NA |
| ALL | m2$_R FI$ : *Mass* | NA | NA | 0.41 (-0.64, 1.49) | -0.36 (-1.39, 0.67) | NA | NA | NA | NA | NA | NA | NA | NA | NA |
| ALL | m1$_O PCr$ | NA | NA | -0.08 (-0.83, 0.66) | 0.64 (-0.1, 1.36) | NA | NA | NA | NA | NA | NA | NA | NA | NA |
| ALL | m1$_O PCr$ : *Mass* | NA | NA | -0.25 (-1.09, 0.6) | 0.22 (-0.54, 1.01) | NA | NA | NA | NA | NA | NA | NA | NA | NA |
| ALL | m2$_O PCr$ | NA | NA | -0.11 (-1.18, 1) | 0.18 (-0.96, 1.25) | NA | NA | NA | NA | NA | NA | NA | NA | NA |
| ALL | m2$_O PCr$ : *Mass* | NA | NA | 0.2 (-0.68, 1.09) | 0.77 (-0.23, 1.8) | NA | NA | NA | NA | NA | NA | NA | NA | NA |

| Metric | Parameter | Bird | Hard Invert | Large Mammal | Plant | Carrion | Egg | Herptile | Seed | Small Mammal | Soft Invert | Fish | Fruit | Root |
|---|---|---|---|---|---|---|---|---|---|---|---|---|---|---|
| ALL | $m1_a DNE$ | NA | NA | -0.55 (-1.4, 0.27) | 0.66 (-0.1, 1.46) | NA | NA | NA | NA | NA | NA | NA | NA | NA |
| ALL | $m1_a DNE : Mass$ | NA | NA | -0.04 (-0.94, 0.88) | -0.99 (-1.84, -0.15) | NA | NA | NA | NA | NA | NA | NA | NA | NA |
| ALL | $m2_a DNE$ | NA | NA | -0.13 (-1.27, 0.94) | -0.27 (-1.41, 0.86) | NA | NA | NA | NA | NA | NA | NA | NA | NA |
| ALL | $m2_a DNE : Mass$ | NA | NA | -0.01 (-1.22, 1.2) | 0.56 (-0.62, 1.75) | NA | NA | NA | NA | NA | NA | NA | NA | NA |
| DNE | $Intercept_1$ | NA | -2.62 (-3.69, -1.65) | NA | -0.71 (-1.99, 0.31) | NA | 0.39 (-0.29, 0.99) | -1.45 (-2.07, -0.88) | 1.73 (0.99, 2.42) | -3.19 (-4.45, -2) | -0.31 (-1.09, 0.46) | NA | NA | NA |
| DNE | $Intercept_2$ | NA | -0.46 (-1.41, 0.44) | NA | 1.71 (0.62, 2.76) | NA | 2.49 (1.75, 3.24) | 0.75 (0.23, 1.31) | 2.86 (2.12, 3.62) | -0.8 (-1.85, 0.34) | 1.28 (0.54, 2.07) | NA | NA | NA |
| DNE | $Intercept_3$ | NA | 1.06 (0.15, 2.04) | NA | 3.07 (1.9, 4.28) | NA | 4.47 (3.3, 5.87) | 3.05 (2.3, 3.89) | 3.99 (3.1, 5) | 0.72 (-0.32, 1.94) | 2.34 (1.52, 3.22) | NA | NA | NA |
| DNE | Mass | NA | -1.42 (-2.06, -0.81) | NA | 0.39 (-0.18, 0.95) | NA | -0.29 (-0.76, 0.17) | -0.84 (-1.3, -0.39) | 0.06 (-0.6, 0.7) | -0.94 (-1.53, -0.37) | -1.12 (-1.7, -0.56) | NA | NA | NA |
| DNE | $Phylo_s d$ | NA | 1.29 (0.48, 2.28) | NA | 1.6 (0.49, 2.7) | NA | 0.56 (0.05, 1.39) | 0.41 (0.03, 1.06) | 0.69 (0.06, 1.67) | 1.68 (0.78, 2.68) | 1.44 (0.57, 2.38) | NA | NA | NA |
| DNE | $m1_a DNE$ | NA | 0.69 (0.03, 1.36) | NA | 0.66 (-0.05, 1.38) | NA | -0.49 (-1.11, 0.13) | 0.84 (0.28, 1.41) | 0.18 (-0.53, 0.91) | -0.23 (-0.91, 0.45) | 0.23 (-0.4, 0.86) | NA | NA | NA |
| DNE | $m1_a DNE : Mass$ | NA | -1.2 (-1.93, -0.5) | NA | -0.68 (-1.38, 0.03) | NA | 0.18 (-0.45, 0.81) | 0.51 (-0.1, 1.15) | 0.07 (-0.67, 0.8) | 0.3 (-0.36, 1) | -0.98 (-1.67, -0.31) | NA | NA | NA |
| DNE | $m2_a DNE$ | NA | 0.56 (-0.16, 1.3) | NA | 0.12 (-0.64, 0.9) | NA | 0.94 (0.28, 1.6) | -0.75 (-1.33, -0.17) | 0.94 (0.11, 1.79) | -0.82 (-1.61, -0.03) | 0.39 (-0.34, 1.09) | NA | NA | NA |
| DNE | $m2_a DNE : Mass$ | NA | 0.22 (-0.38, 0.83) | NA | 0.49 (-0.2, 1.15) | NA | -0.37 (-1.01, 0.27) | -0.43 (-1.06, 0.17) | -0.15 (-0.96, 0.63) | 0.65 (-0.07, 1.43) | 0.41 (-0.28, 1.09) | NA | NA | NA |
| OPCr | $Intercept_1$ | NA | NA | NA | -0.76 (-1.97, 0.18) | -0.95 (-2.11, 0.02) | NA | NA | NA | NA | NA | NA | -2.58 (-4.08, -1.14) | NA |
| OPCr | $Intercept_2$ | NA | NA | NA | 1.6 (0.55, 2.54) | 1.29 (0.26, 2.31) | NA | NA | NA | NA | NA | NA | -0.2 (-1.59, 1.17) | NA |
| OPCr | $Intercept_3$ | NA | NA | NA | 3 (1.82, 4.23) | 3.29 (2.11, 4.6) | NA | NA | NA | NA | NA | NA | 1.55 (0.16, 2.99) | NA |

| Metric | Parameter | Bird | Hard Invert | Large Mammal | Plant | Carrion | Egg | Herptile | Seed | Small Mammal | Soft Invert | Fish | Fruit | Root |
|---|---|---|---|---|---|---|---|---|---|---|---|---|---|---|
| OPCr | Mass | NA | NA | NA | 0.21 (-0.31, 0.72) | 0.36 (-0.16, 0.88) | NA | NA | NA | NA | NA | NA | -0.3 (-0.96, 0.32) | NA |
| OPCr | $Phylo_sd$ | NA | NA | NA | 1.43 (0.4, 2.46) | 1.59 (0.59, 2.61) | NA | NA | NA | NA | NA | NA | 2.84 (1.9, 3.8) | NA |
| OPCr | $m1_OPCr$ | NA | NA | NA | 0.43 (-0.2, 1.08) | 0.33 (-0.3, 0.98) | NA | NA | NA | NA | NA | NA | 0.41 (-0.25, 1.08) | NA |
| OPCr | $m1_OPCr$ : *Mass* | NA | NA | NA | -0.11 (-0.81, 0.61) | 0.48 (-0.23, 1.16) | NA | NA | NA | NA | NA | NA | 0.43 (-0.3, 1.16) | NA |
| OPCr | $m2_OPCr$ | NA | NA | NA | 0.49 (-0.33, 1.3) | -0.25 (-1.09, 0.58) | NA | NA | NA | NA | NA | NA | 0.76 (-0.17, 1.73) | NA |
| OPCr | $m2_OPCr$ : *Mass* | NA | NA | NA | 0.2 (-0.53, 0.93) | -0.42 (-1.15, 0.3) | NA | NA | NA | NA | NA | NA | -0.71 (-1.53, 0.08) | NA |
| RFI | $Intercept_1$ | NA | NA | NA | NA | NA | 0.53 (-0.22, 1.18) | NA | 1.82 (1.03, 2.59) | NA | NA | NA | NA | NA |
| RFI | $Intercept_2$ | NA | NA | NA | NA | NA | 2.67 (1.89, 3.48) | NA | 3.04 (2.14, 4.01) | NA | NA | NA | NA | NA |
| RFI | $Intercept_3$ | NA | NA | NA | NA | NA | 4.66 (3.43, 6.08) | NA | 4.59 (3.23, 6.25) | NA | NA | NA | NA | NA |
| RFI | Mass | NA | NA | NA | NA | NA | 0.02 (-0.47, 0.51) | NA | 0.02 (-0.68, 0.69) | NA | NA | NA | NA | NA |
| RFI | $Phylo_sd$ | NA | NA | NA | NA | NA | 0.6 (0.05, 1.47) | NA | 0.75 (0.08, 1.72) | NA | NA | NA | NA | NA |
| RFI | $m1_RFI$ | NA | NA | NA | NA | NA | 0.13 (-0.33, 0.6) | NA | 0.08 (-0.49, 0.66) | NA | NA | NA | NA | NA |
| RFI | $m1_RFI$ : *Mass* | NA | NA | NA | NA | NA | 0.12 (-0.26, 0.51) | NA | 0.02 (-0.48, 0.55) | NA | NA | NA | NA | NA |
| RFI | $m2_RFI$ | NA | NA | NA | NA | NA | 0.97 (0.38, 1.63) | NA | 1.2 (0.36, 2.09) | NA | NA | NA | NA | NA |
| RFI | $m2_RFI$ : *Mass* | NA | NA | NA | NA | NA | -0.46 (-0.98, 0.06) | NA | 0.16 (-0.67, 1.07) | NA | NA | NA | NA | NA |
| RLGA | $Intercept_1$ | -2.47 (-3.27, -1.76) | -2.84 (-3.96, -1.9) | 0.66 (-0.14, 1.4) | NA | -0.63 (-1.68, 0.31) | NA | -1.31 (-1.9, -0.73) | 1.62 (0.94, 2.31) | -3.68 (-4.7, -2.73) | -0.97 (-2.05, -0.08) | 0.46 (-0.8, 1.74) | -2.33 (-3.73, -1.01) | 2.44 (1.7, 3.27) |

| Metric | Parameter | Bird | Hard Invert | Large Mammal | Plant | Carrion | Egg | Herptile | Seed | Small Mammal | Soft Invert | Fish | Fruit | Root |
|---|---|---|---|---|---|---|---|---|---|---|---|---|---|---|
| RLGA | Intercept$_2$ | 0.57 (-0.04, 1.17) | -0.7 (-1.58, 0.17) | 2.03 (1.25, 2.86) | NA | 1.61 (0.65, 2.64) | NA | 0.71 (0.16, 1.28) | 2.83 (2.04, 3.7) | -1.16 (-1.85, -0.41) | 0.63 (-0.36, 1.54) | 2.7 (1.41, 4.12) | -0.12 (-1.41, 1.15) | 3.6 (2.61, 4.76) |
| RLGA | Intercept$_3$ | 3.58 (2.66, 4.63) | 0.76 (-0.11, 1.75) | 2.83 (1.99, 3.79) | NA | 3.55 (2.39, 4.83) | NA | 2.93 (2.18, 3.79) | 4.35 (3.06, 5.9) | 0.35 (-0.32, 1.11) | 1.66 (0.65, 2.63) | 3.82 (2.4, 5.45) | 1.57 (0.29, 2.92) | NA |
| RLGA | Mass | -0.86 (-1.28, -0.44) | -1.08 (-1.64, -0.59) | 1.84 (1.24, 2.49) | NA | 0.24 (-0.22, 0.7) | NA | -1.03 (-1.45, -0.62) | -0.46 (-1.08, 0.15) | -1.04 (-1.5, -0.59) | -0.98 (-1.53, -0.48) | 0.57 (-0.01, 1.18) | -0.11 (-0.74, 0.48) | 0.02 (-0.72, 0.77) |
| RLGA | Phylo$_{sd}$ | 0.62 (0.08, 1.33) | 1.15 (0.19, 2.41) | 1.07 (0.3, 1.95) | NA | 1.51 (0.64, 2.49) | NA | 0.52 (0.04, 1.24) | 0.76 (0.1, 1.68) | 0.77 (0.12, 1.61) | 1.52 (0.54, 2.55) | 2.1 (1.16, 3.18) | 2.46 (1.53, 3.48) | 0.71 (0.06, 1.78) |
| RLGA | RLGA | -1.25 (-1.75, -0.77) | 1.35 (0.75, 2) | -1.67 (-2.43, -0.96) | NA | -0.17 (-0.76, 0.39) | NA | -0.37 (-0.79, 0.04) | 0.56 (-0.04, 1.15) | -1.84 (-2.45, -1.26) | 0.52 (-0.11, 1.12) | -0.13 (-0.84, 0.55) | 1.38 (0.62, 2.16) | 0.74 (0.03, 1.44) |
| RLGA | RLGA:Mass | 0.72 (0.32, 1.13) | -0.25 (-0.72, 0.22) | -0.06 (-0.65, 0.51) | NA | 0.15 (-0.3, 0.62) | NA | 0.02 (-0.39, 0.41) | 0.6 (-0.03, 1.28) | 0.82 (0.34, 1.31) | 0.11 (-0.43, 0.66) | 0.34 (-0.2, 0.9) | -0.06 (-0.66, 0.54) | 0.37 (-0.3, 1.07) |
